# A transcriptome-based phylogeny of Scarabaeoidea confirms the sister group relationship of dung beetles and phytophagous pleurostict scarabs (Coleoptera)

**DOI:** 10.1101/2023.03.11.532172

**Authors:** Lars Dietz, Matthias Seidel, Jonas Eberle, Bernhard Misof, Thaynara L. Pacheco, Lars Podsiadlowski, Sasanka Ranasinghe, Nicole L. Gunter, Oliver Niehuis, Christoph Mayer, Dirk Ahrens

## Abstract

Scarab beetles (Scarabaeidae) are a diverse and ecologically important group of angiosperm-associated insects. As conventionally understood, scarab beetles comprise two major lineages: dung beetles and the phytophagous Pleurosticti. However, previous phylogenetic analyses have not been able to convincingly answer the question whether or not the two lineages form a monophyletic group. Here we report our results from phylogenetic analyses of more than 4,000 genes mined from transcriptomes of more than 50 species of Scarabaeidae and other Scarabaeoidea. Our results provide convincing support for the monophyly of Scarabaeidae, confirming the debated sister group relationship of dung beetles and phytophagous pleurostict scarabs. Supermatrix-based maximum likelihood and multispecies coalescent phylogenetic analyses strongly imply the subfamily Melolonthinae as currently understood being paraphyletic. We consequently suggest various changes in the systematics of Melolonthinae: Sericinae Kirby, 1837 stat. rest. and sensu n. to include the tribes Sericini, Ablaberini and Diphucephalini, and Sericoidinae Erichson, 1847 stat. rest. and sensu n. to include the tribes Automoliini, Heteronychini, Liparetrini, Maechidiini, Scitalini, Sericoidini, and Phyllotocini. Both subfamilies appear to consistently form a monophyletic sister group to all remaining subfamilies so far included within pleurostict scarabs except Orphninae. Our results represent a major step towards understanding the diversification history of one of the largest angiosperm-associated radiations of beetles.

## Introduction

The evolution of large parts of extant terrestrial biodiversity has been driven by the evolutionary success of angiosperm plants; these radiations have been linked to increased productivity and growth rates of angiosperm vegetation (de Boer et al. 2012), the rise of ectomycorrhiza enhancing chemical weathering of soils (Taylor et al. 2011, 2012), and the promotion of soil nutrient release by angiosperm litter that is easily decomposed (Berendse & Scheffer 2009). While the diversification of many insects, and especially that of beetles, was directly or indirectly fostered by that of angiosperms (Hunt et al. 2007; Ahrens et al. 2014; McKenna et al. 2019), the evolutionary mechanisms and timescales of angiosperm-dependent radiations have remained poorly understood, as the phylogenetic relationships of many lineages remained insufficiently known. This is especially true for scarab beetles (Scarabaeidae), which represent a diverse lineage of beetles feeding predominantly on either angiosperm plants or mammal dung. Although traditionally grouped into a single family (Scholtz & Grebennikov 2005), it is divided into two major lineages: 1) the plant-feeding lineage Pleurosticti, which include, for example, rose chafers, rhinoceros beetles, and Christmas beetles, and 2) a clade of taxa feeding on mammal dung (Aphodiinae + Scarabaeinae).

Molecular phylogenetic analyses of the Scarabaeidae have been controversial, with only nine out of 21 recently published studies reporting Scarabaeidae being monophyletic (Table 1). Scarabaeidae are part of a wider clade (Scarabaeoidea) which also includes Lucanidae (stag beetles), Geotrupidae (earth-boring dung beetles) and several other families (Scholtz & Grebennikov 2005), and the monophyly of this superfamily has been confirmed by all major molecular studies (e.g., Zhang et al 2018, McKenna et al. 2019). The current classification of Scarabaeoidea (Scholtz & Grebennikov 2005) is founded on morphological evidence (Lawrence & Newton 1995). Yet, despite extensive research on the morphology of Scarabaeoidea by Browne & Scholtz (1998, 1999), the monophyly of Scarabaeidae has yet to be adequately tested. The analysis of Browne & Scholtz (1998) included only lineages of “Scarabaeidae” which were rooted with a single outgroup, while the analysis of Browne & Scholtz (1999) coded “Scarabaeidae” as a single terminal taxon. A recent cladistic analysis based on morphology in a wider systematic framework (Lawrence et al. 2011) did not recover Scarabaeidae as a monophyletic group.

**Table 1.**
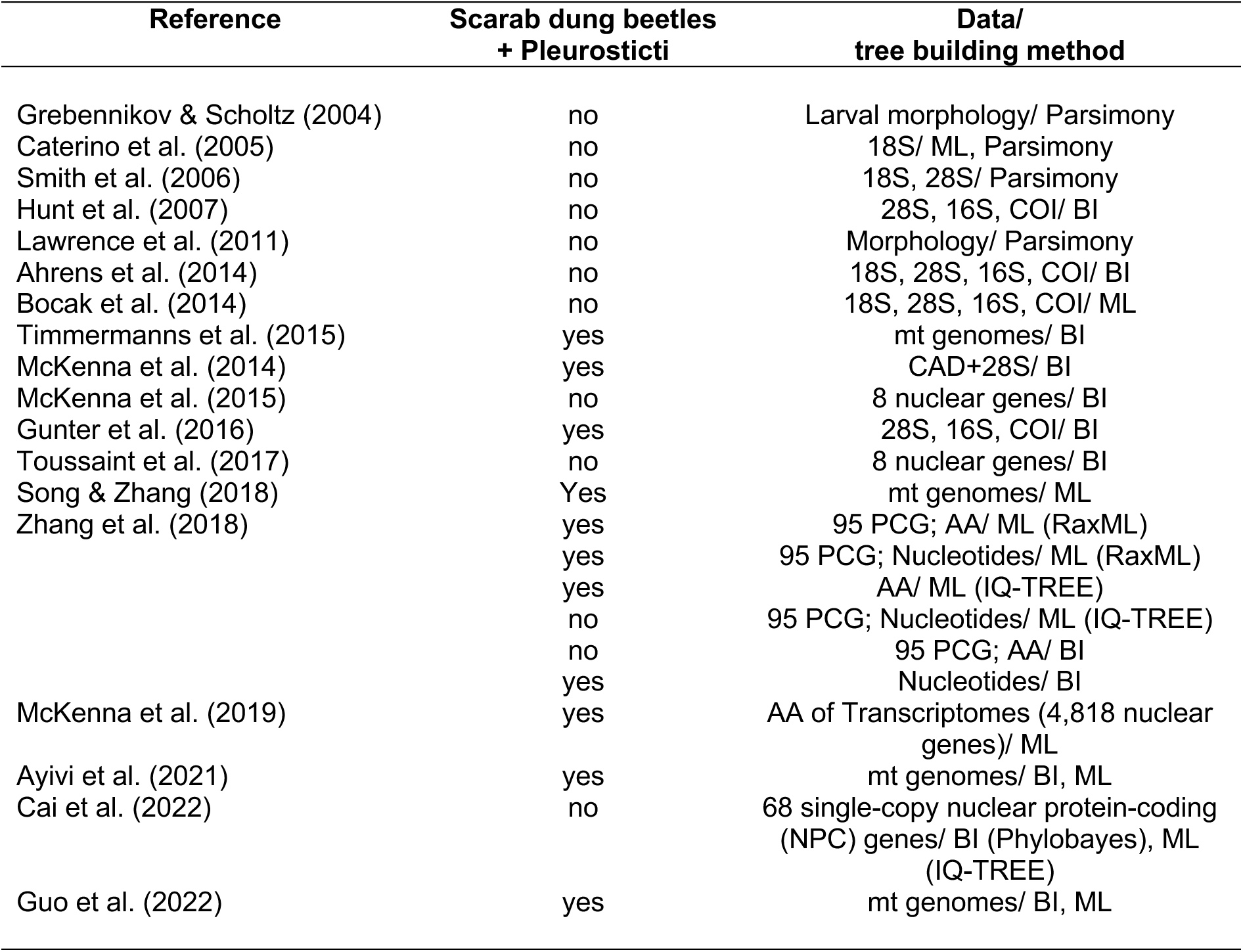
Overview of results on monophyly of Scarabaeidae retrieved in different previous phylogenetic analyses.

| Reference | Scarab dung beetles<br>+ Pleurosticti | Data/<br>tree building method |
| --- | --- | --- |
| Grebennikov & Scholtz (2004) | no | Larval morphology/ Parsimony |
| Caterino et al. (2005) | no | 18S/ ML, Parsimony |
| Smith et al. (2006) | no | 18S, 28S/ Parsimony |
| Hunt et al. (2007) | no | 28S, 16S, COI/ BI |
| Lawrence et al. (2011) | no | Morphology/ Parsimony |
| Ahrens et al. (2014) | no | 18S, 28S, 16S, COI/ BI |
| Bocak et al. (2014) | no | 18S, 28S, 16S, COI/ ML |
| Timmermanns et al. (2015) | yes | mt genomes/ BI |
| McKenna et al. (2014) | yes | CAD+28S/ BI |
| McKenna et al. (2015) | no | 8 nuclear genes/ BI |
| Gunter et al. (2016) | yes | 28S, 16S, COI/ BI |
| Toussaint et al. (2017) | no | 8 nuclear genes/ BI |
| Song & Zhang (2018) | Yes | mt genomes/ ML |
| Zhang et al. (2018) | yes | 95 PCG; AA/ ML (RaxML) |
|  | yes | 95 PCG; Nucleotides/ ML (RaxML) |
|  | yes | AA/ ML (IQ-TREE) |
|  | no | 95 PCG; Nucleotides/ ML (IQ-TREE) |
|  | no | 95 PCG; AA/ BI |
|  | yes | Nucleotides/ BI |
| McKenna et al. (2019) | yes | AA of Transcriptomes (4,818 nuclear genes)/ ML |
| Ayivi et al. (2021) | yes | mt genomes/ BI, ML |
| Cai et al. (2022) | no | 68 single-copy nuclear protein-coding (NPC) genes/ BI (Phylobayes), ML (IQ-TREE) |
| Guo et al. (2022) | yes | mt genomes/ BI, ML |

The uncertainty surrounding the monophyly of Scarabaeidae prompted discussions about the classification of the family (Kohlmann & Morón 2003). Apart from many unresolved issues of classification within the group, the question arose whether to split scarab dung beetles and pleurostict scarabs (Erichson 1848) in two or more families (Morón 1997; Cherman & Morón 2014). Prior to the era of molecular phylogenies, there has been an impressive array of classification schemes on Scarabaeidae and related families (Jansens 1949; Crowson 1955; Balthasar 1963; Endroedi 1966; Medvedev 1976; Iablokoff-Khnzorian 1977; Howden 1982; Lawrence & Newton 1982, 1995; Paulian & Baraud 1982; Morón 1984, 1997, 2003; Scholtz 1990; Browne & Scholtz 1995; Nikolaev 1995; Kohlmann & Morón 2003; Ratcliffe & Jameson 2004; Scholtz & Grebennikov 2005; Smith 2006; Smith et al. 2006). As robust and convincing evidence is lacking for most of these classification schemes, many systematic and taxonomic studies arbitrarily followed one of these classifications according to the author’s opinion or geographic provenience (e.g., Cherman & Morón 2014).

Here, we readdress the question about the controversial sister group relationship of scarab dung beetles and pleurostict scarabs (Ahrens et al. 2014), which is fundamental to gain a more complete understanding of the evolutionary impact of Angiosperms on the diversification of Scarabaeidae. If the monophyly of Scarabaeidae is confirmed, then it raises the question of why angiosperms and their related follow-up radiations (e.g., that of large herbivore mammals; see Ahrens et al. 2014) seemingly had a more significant impact on the radiation of this lineage than on any other scarabaeoid beetle lineage. To this end, we expand the phylogenomic data set compiled by McKenna et al. (2019) within additional taxa of Scarabaeoidea. The expanded taxonomic sampling is used to assess which aspects of the data could result in the inference of incompatible topologies (e.g., Zhang et al. 2018; Cai et al. 2022).

## Material and methods

### Taxon sampling and new transcriptome data

We analyzed 57 transcriptomes that covered almost all major families of Scarabaeoidea and two outgroup taxa (Table S1). Fifteen of these transcriptomes had been published by McKenna et al. (2019). Two transcriptomes were sequenced in context of the 1KITE project but had not been analyzed and published before. For details relating to the extraction of total mRNA and fragmentation, construction of cDNA libraries, and tagging of these two datasets see Peters et al. (2017). 40 additional transcriptomes were specifically generated in context of this study (Table S1).

All mRNA libraries were sequenced with Illumina HiSeq 2000 sequencers (Illumina, San Diego, CA, USA), using paired-end 150-bp reads.

Extraction of RNA, complementary deoxyribonucleic acid (cDNA) library construction, library normalization, and Illumina sequencing were carried out by a commercial sequencing company (Starseq, Mainz, Germany).

All raw nucleotide sequences are deposited at the National Center for Biotechnology Information (NCBI), Sequence Read Archive (see Table S1 for accession numbers). The raw data of previously published transcriptomes of 17 Scarabaeoidea (see above) and of two outgroup taxa (*Ocypus brunnipes* (Staphylinidae) and *Helophorus nanus* (Helophoridae)) (McKenna et al. 2019) were downloaded from NCBI. All raw nucleotide reads from both newly sequenced and published transcriptomes were trimmed with TrimGalore 0.6.6 (Krueger et al. 2021) and assembled with Trinity 2.11.0 (Grabherr et al. 2011) using the software’s default settings. Transcriptome assemblies (see Table S1 for accession numbers) are deposited at the Transcriptome Shotgun Assembly (TSA) Database, NCBI Bioproject ID PRJNA906571 for newly sequenced transcriptomes or PRJNA936991 for re-assemblies of published transcriptomes (http://www.ncbi.nlm.nih.gov/bioproject). The assemblies were filtered with a custom Perl script (trinity_longest_d.pl, see Supplement) to retain only the longest isoform per locus, as loci with multiple isoforms could otherwise be falsely discarded as paralogs in the gene orthology assessment step.

### Data extraction and alignment

Tab-delimited files were downloaded from the OrthoDB10 database (www.orthodb.org) for all groups of orthologous genes (= ortholog groups) at the hierarchical level Coleoptera that were present in at least eight of the nine coleopteran genomes in the database and single-copy in all of them. This included a total of 4,296 genes. This is similar to the principle of universal single-copy orthologs (USCOs) as used by the program BUSCO (Simão et al. 2015). However, USCOs have to be present and single-copy in at least 90% of all known genomes of a taxonomic group. In this case, this would mean that the genes would have to be present in all nine annotated coleopteran genomes available at the time. To avoid excluding genes that may be absent simply due to the incompleteness of one of the nine genome assemblies, we decided to include genes present in only eight of the nine coleopteran genomes.

Furthermore, we downloaded the official gene sets (OGS) for all nine available coleopteran genomes from OrthoDB. Tab-delimited files were modified for use in Orthograph 0.7.1 (Petersen et al. 2017) and used together with the OGS, to create a SQLite database of the genetic information with that program. Hidden Markov models (HMMs) were created with Orthograph from the available amino acid sequences of each ortholog group. HMMs were used then to extract the target genes from the filtered Trinity contigs of each specimen with Orthograph, using the software’s default settings.

USCO nucleotide and corresponding amino acid sequences that were identified and inferred with Orthograph were aligned in two different ways. We first aligned the inferred amino acid sequences against the HMMs from Orthograph with hmmalign (part of the HMMER 3.3 package; Eddy 2011; http://eddylab.org/software/hmmer/hmmer.org). The amino acid alignment was then used as a blueprint to align the corresponding nucleotide sequences with pal2nal 14.1 (Suyama et al. 2006). Alignment regions not covered by the HMMs were removed with a custom Perl script (hmmalign_cut2_d.pl, Supplement Files). We additionally aligned the amino acid sequences with MAFFT 7.305b (Katoh & Standley 2013) using the L-INS-i algorithm. The corresponding nucleotide sequence alignments were inferred using again the software pal2nal. Poorly aligned regions in the amino acid sequence alignments were identified with ALISCORE 2.0 (Misof & Misof 2009; Kück et al. 2010) and removed from the amino acid sequence alignments and corresponding nucleotide sequence alignments with the software ALICUT 2.31 (available from: https://github.com/PatrickKueck/AliCUT). Outlier sequences were identified and removed with the software OliInSeq 0.9.3 (https://github.com/cmayer/OliInSeq) using the software’s default parameters. As the third codon position is typically hypervariable and often exhibits inhomogeneous nucleotide frequencies, we also used a custom Perl script (extract_codpos_d.pl, see Supplement) to generate nucleotide sequence alignments in which the third codon position was removed.

To test the monophyly of Scarabaeidae with data from another dataset, we downloaded the OrthoDB10 Endopterygota data from BUSCO 4.0.6 (Simão et al. 2015; Manni et al. 2021), including Hidden Markov Models (HMMs) and information files, from the BUSCO website (busco.ezlab.org). This set comprises 2,120 genes that are present in single copy in at least 90% of all known genomes of Endopterygota (hereafter referred to as Endopterygota USCOs). The OGS for all 56 species in that dataset were downloaded from OrthoDB and used to create an SQLite database with Orthograph. Together with the HMMs from BUSCO, the information was used to extract the Endopterygota USCO genes of each specimen with Orthograph as described above.

### Phylogenetic tree inference

We inferred phylogenetic trees from the Coleoptera and Endopterygota data sets, analyzing the amino acid sequence alignments (AA) generated with hmmalign and MAFFT and the corresponding nucleotide sequence alignments (NT) inferred with pal2nal. We then considered two sets of nucleotide sequence alignments: those that included all three codon positions (NT123) and those that include only first and second codon positions (NT12). The multiple sequence alignments were analyzed using coalescent-based and concatenation-based tree inference methods. For conducting the concatenation-based phylogenetic analyses, the multiple sequence alignments of a given type (i.e., amino acid, nucleotide) of all genes were concatenated with a custom Perl script (concat_eogs_part_d.pl, see Supplement). The resulting super-alignments were then analyzed with IQ-TREE 2.1.2 (Minh et al. 2020). The datasets were partitioned by gene. The best-fitting model and partitioning scheme were inferred with ModelFinder, using the IQ-TREE option -m MFP+MERGE (Kalyaanamoorthy et al. 2017; Chernomor et al. 2016). Branch support was assessed with approximate likelihood ratio tests (aLRT) and via ultrafast bootstrapping (Hoang et al. 2018) applying 1,000 replicates and nearest neighbor interchange (NNI) as tree rearrangement method. For conducting coalescent-based phylogenetic analyses, we first calculated phylogenetic trees of each gene with IQ-TREE, determining the best-fitting model with ModelFinder. The resulting gene trees were used for analyses with ASTRAL 5.6.1 (Zhang et al. 2018) that we used to conduct the coalescent-based phylogenetic analysis. All trees were rooted with *Helophorus* (Helophoridae) and *Ocypus* (Staphylinidae) as outgroups.

To test the effect of missing data, we generated reduced versions of all nucleotide and amino-acid datasets in which positions with a taxon coverage of less than 70% were removed with a custom Perl script (removegaps_d.pl, Suppl. File 5). The previously described phylogenetic analyses were repeated using these reduced datasets.

Furthermore, we examined the effect of varying substitution rates between different genes. For this test, we divided datasets containing at least 70% complete positions into sets of fast and of slowly evolving genes. Using a custom script (pairwise_id2.pl, see Supplement Files) we calculated the pairwise sequence identity within each gene alignment according to Sharma et al. (2014). We then divided both the hmmalign- and MAFFT-inferred multiple sequence alignments of individual genes into two sets with high and with low pairwise sequence identity, each including 50% of the genes. We then conducted concatenation- and coalescent-based analyses on these sets using the same methods as described above.

### Topology tests

To assess support for the monophyly of Scarabaeidae in our datasets, we conducted a number of topology tests on the six different datasets (i.e., NT12, NT123 and AA, inferred using hmmalign or MAFFT). First, we computed the likelihood scores of maximum likelihood trees constrained to support different topologies for each dataset with IQ-TREE. The constrained trees covered all possible phylogenetic relationships of the following four taxa: Glaphyridae, Hybosoridae, Scarabaeinae + Aphodiinae, Pleurosticti. We compared the likelihood of the constraint trees using a variety of resampling tests in IQ-TREE using RELL approximation (Kishino et al. 1990) with 10,000 iterations, including bootstrap proportion (BP), Kishino-Hasegawa test (KH; Kishino & Hasegawa 1989), Shimodaira-Hasegawa test (SH; Shimodaira & Hasegawa 1999), expected likelihood weights (ELW; Strimmer and Rambaut 2002), and the approximately unbiased (AU) test (Shimodaira 2002).

We used the same strategy to assess support for all possible phylogenetic relationships between Dynastinae and the genera *Anomala* and *Adoretus* (both part of the potentially paraphyletic Rutelinae) as well as between Melolonthinae s. str., Hopliini, and Cetoniinae + Rutelinae + Dynastinae (hereafter referred to as CRD). We further tested the monophyly of Scarabaeidae using four-cluster likelihood mapping (Strimmer and von Haeseler 1997) in IQ-TREE. For this purpose, we divided the taxon set into Hybosoridae, Scarabaeinae + Aphodiinae, Pleurosticti, and all others. All 4,410 unique quartets containing one taxon of each quartet were tested for their support for the three possible four-taxon trees.

## Results

### Data completeness

Nucleotide sequences from all but one of 4,296 Coleoptera-specific genes and from all 2,120 Endopterygota USCO genes were successfully recovered. In total, the dataset of the Coleoptera-specific genes aligned with hmmalign comprised 6,960,192 nucleotide positions, of which 3,670,744 were parsimony informative. The overall alignment completeness was 62.7%. The corresponding MAFFT-aligned dataset comprised 3,929,520 nucleotides, of which 2,186,562 were parsimony informative. The alignment completeness was 79.6%. The dataset of Endopterygota USCO genes comprised 2,337,876 nucleotides aligned with hmmalign, of which 1,126,272 were parsimony informative. The alignment completeness was 62.8%. The corresponding dataset aligned with MAFFT was 1,649,496 nucleotide positions long, of which 893,640 were parsimony informative. The alignment completeness was 82.2%.

### Phylogenetic analyses

The choice of the alignment software (hmmalign vs. MAFFT) had little impact on the results of the phylogenetic analyses (Fig. 1; Suppl. Figs 1 and 2), and neither did the presence of incomplete alignment sites. However, the choice of the alignment software had an impact on the inferred position of Passalidae (Suppl. Figs 3 and 4). Both sets of genes yielded very similar phylogenies, except that in trees based on the smaller Endopterygota USCO dataset, the single species of passalid included in our analyses was consistently placed as sister to Glaphyridae + Hybosoridae + Scarabaeidae, while in the larger Coleoptera-specific set, its position differed in the various phylogenetic analyses. Including only fast or slowly evolving genes did not lead to noteworthy consistent differences in topology, although support values were generally lower than with the complete gene set. The most significant topological differences were found between trees inferred from datasets that included all three codon positions (NT123) and between trees inferred from datasets which included only nucleotides of the first and second codon position and those that were analyzed on the amino acid level. Whether a concatenation-based and coalescent-based approach was used to analyze the data had little impact on the phylogenetic results, except on the placement of the outgroup. All datasets yield well-resolved trees, in which most splits received maximal support.

**Fig. 1.**
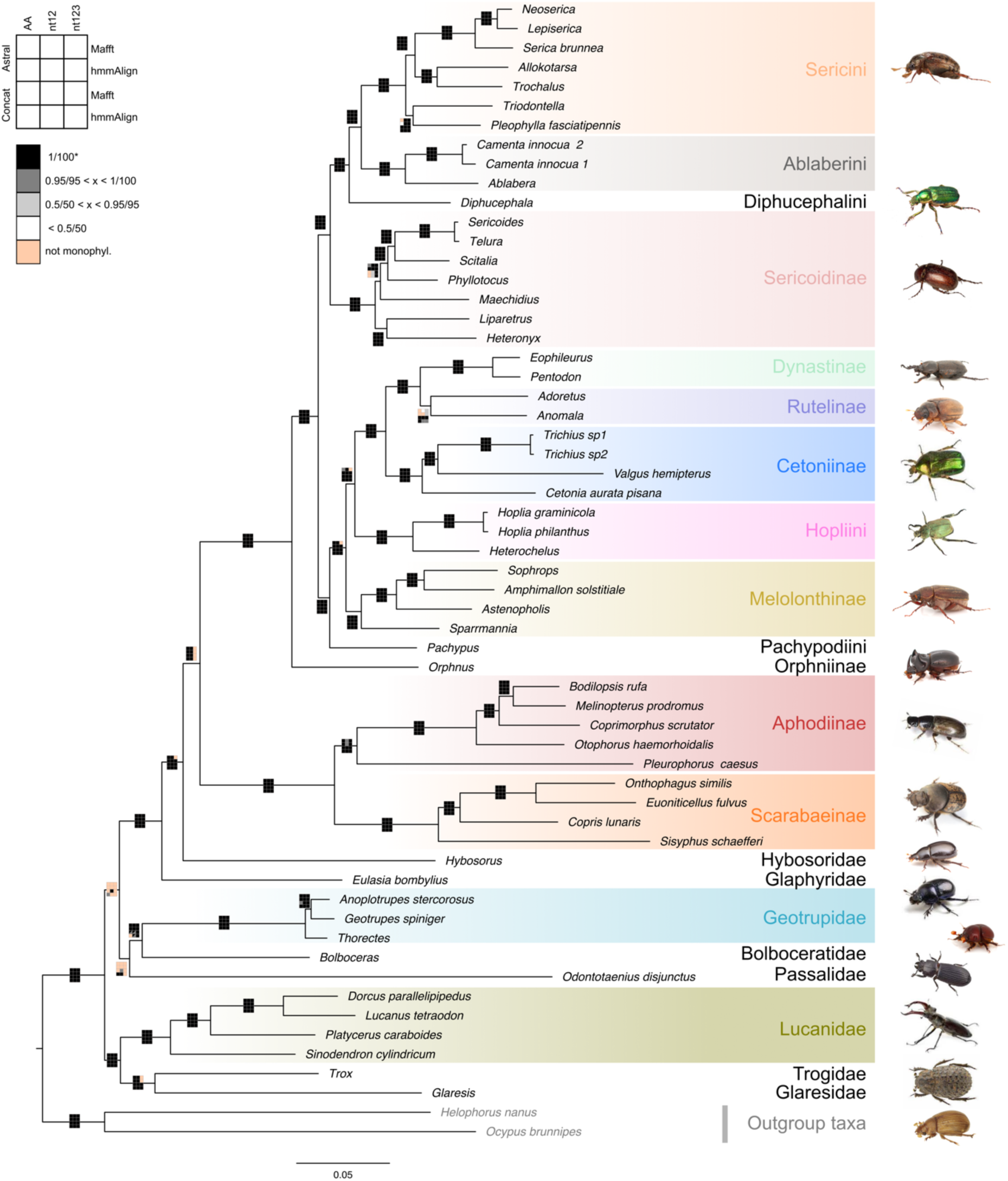
Phylogenetic tree from the concatenated data of the Mafft alignment of the nucleotide data containing (Coleoptera single copy orthologs, full) only the first and second base pair (nt12). Branch support (nn/ xx; see legend) and results of retrieval with other alignments and tree reconstruction approaches as well as the current systematic assignment of lineages are mapped onto the branches.

Scarabaeidae are strongly supported as being monophyletic by all datasets except the nucleotide datasets that include the 3^rd^ codon position (NT123) (Fig. 1). Analysis of the latter suggested Hybosoridae as the sister group of pleurostict Scarabaeidae. All remaining scarabaeoid families that were represented by more than one species in our datasets were consistently supported as being monophyletic.

Phylogenetic analysis of the NT12 and AA supermatrices suggested Lucanidae is the sister group of Glaresidae + Trogidae. They furthermore suggested Geotrupidae + Bolboceratidae is the sister group of a clade that comprised Glaphyridae, Hybosoridae, and Scarabaeidae. Passalidae was placed in various positions within that group depending on the specific analysis. Hybosoridae was found as the sister group of Scarabaeidae. The monophyly of Scarabaeidae was always maximally supported by our data (Fig. 1; Suppl. Figs 1-2). Coalescence-based summary trees of all gene trees differed from the supermatrix-based trees in suggesting Geotrupidae + Bolboceratidae being the sister group of Lucanidae, Glaresidae, and Trogidae. However, some the coalescence-based summary trees inferred from exclusively slowly evolving genes showed the same topology as the concatenation-based trees. The two subgroups of Scarabaeidae, Aphodiinae + Scarabaeinae and Pleurosticti were always strongly supported as being monophyletic, as were the two subfamilies, Aphodiinae and Scarabeinae. Within Pleurosticti, *Orphnus* (Orphninae) was consistently found as sister group of the remaining pleurostict scarabaeid lineages, that were divided into two major clades: the first clade comprised 1) Sericoidinae — Australasian and Neotropical taxa referred to by some authors as Southern World Melolonthinae (Ahrens & Vogler 2008; Ahrens et al. 2011, 2014; Šípek et al. 2016) or Liparetrinae (Lacroix 2007, 2014; Eberle et al. 2019; Pacheco et al. 2022) — and 2) Ablaberini + Sericini + the Australian genus *Diphucephala* (i.e., Diphucephalini), which represented the sister group of the two former tribes. The second clade comprised *Pachypus* (Pachypodini) and three major lineages, which *Pachypus* was found to be sister group of. The phylogenetic relationships among the latter three lineages differed among the inferred trees. These three lineages were: 1) Hopliini, 2) Melolonthini, including the genus *Sparrmannia* (currently placed within the probably polyphyletic Tanyproctini; Eberle et al. 2019), and 3) Cetoniinae + (Dynastinae + Rutelinae). Supermatrix-based phylogenetic analyses consistently provided support for Melolonthini being the sister group of the remaining two lineages. Coalescence-based analysis were inconsistent in regard of the phylogenetic arrangement of the three lineages. While the monophyly of Dynastinae + Rutelinae was consistently well supported, a monophyly of Rutelinae was strongly supported only in concatenation-based trees. In coalescence-based trees, the grouping was not consistently found, and if so, it typically received only low support — a pattern also confirmed by results of previous studies in which monophyly of Rutelinae did not result (e.g., Ahrens & Vogler 2008; Ahrens et al. 2014; Šipek et al. 2016; Neita-Moreno et al. 2019). With Cetoniinae + (Dynastinae + Rutelinae) consistently found nested within the lineages so far classified as Melolonthinae, our study confirmed the paraphyly of Melolonthinae (Ahrens & Vogler 2008; Ahrens et al. 2014; Gunter et al. 2016; Šipek et al. 2016; Neita-Moreno et al. 2019; McKenna et al. 2019).

**Fig. 2.**
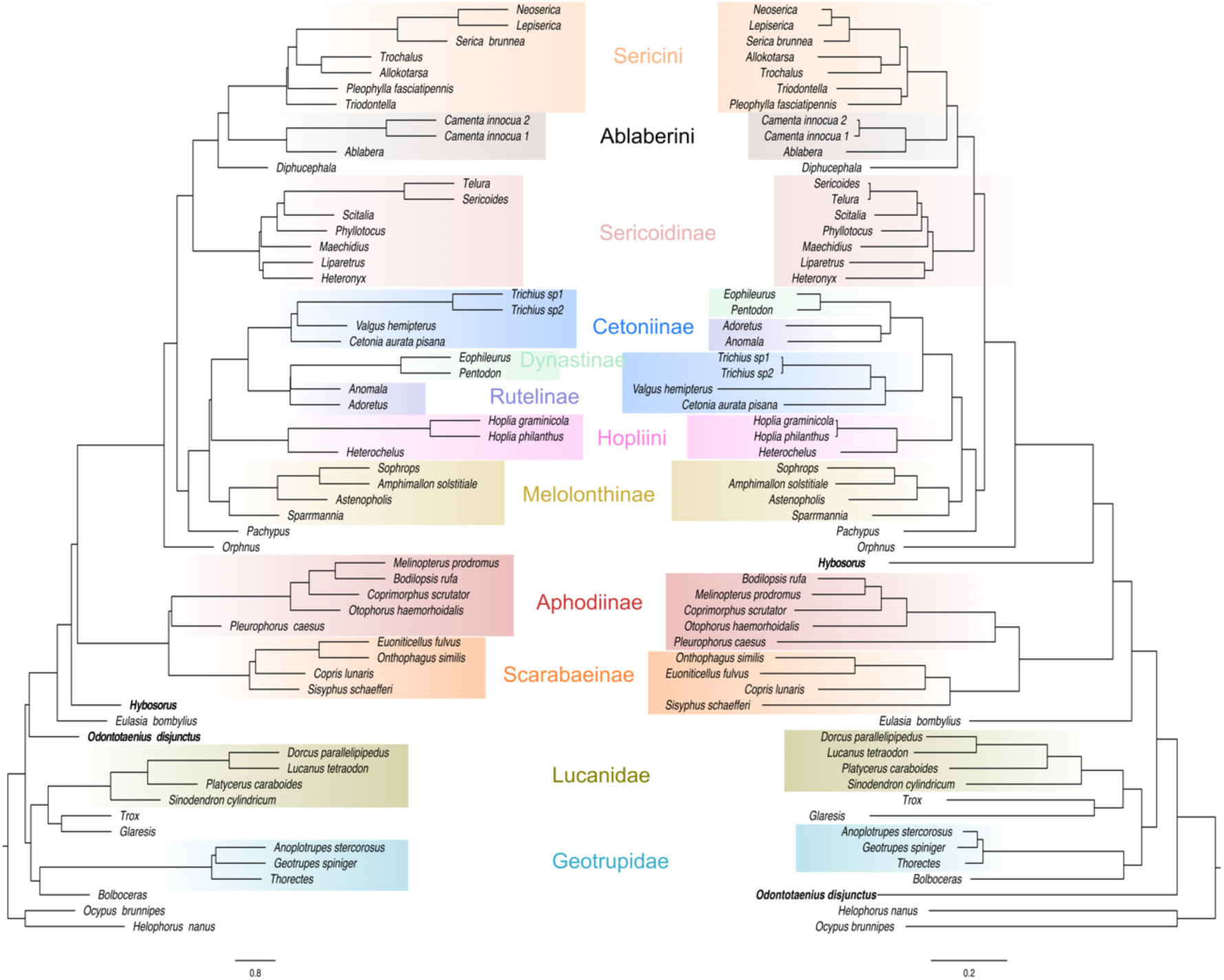
Contrasting tree topologies based on the Coleoptera single copy orthologs obtained with coalescent tree search with Astral (amino acid sequences; left side) and with concatenated data using all nucleotides (right side); major lineages are highlighted; single taxa changing topology are marked in bold.

### Topology tests

In tests assessing the support for monophyly of Scarabaeidae, the originally inferred topology (i.e., monophyly of Scarabaeidae) consistently received the highest support when analyzing the NT12 and AA supermatrices. Likewise, the monophyly of Hybosoridae + Pleurosticti consistently received the highest support when analyzing the NT123 datasets. Alternative trees were rejected with p < 0.001 (Table 2). The topology tests supported the sister group relationship of Hopliini and the CRD clade, rejecting alternative topologies with p < 0.001. Likelihood tests testing the monophyly of Rutelinae supported the latter irrespective of what dataset we analyzed, but often only with p between 0.01 and 0.05 (Tables 3, 4).

**Table 2.**
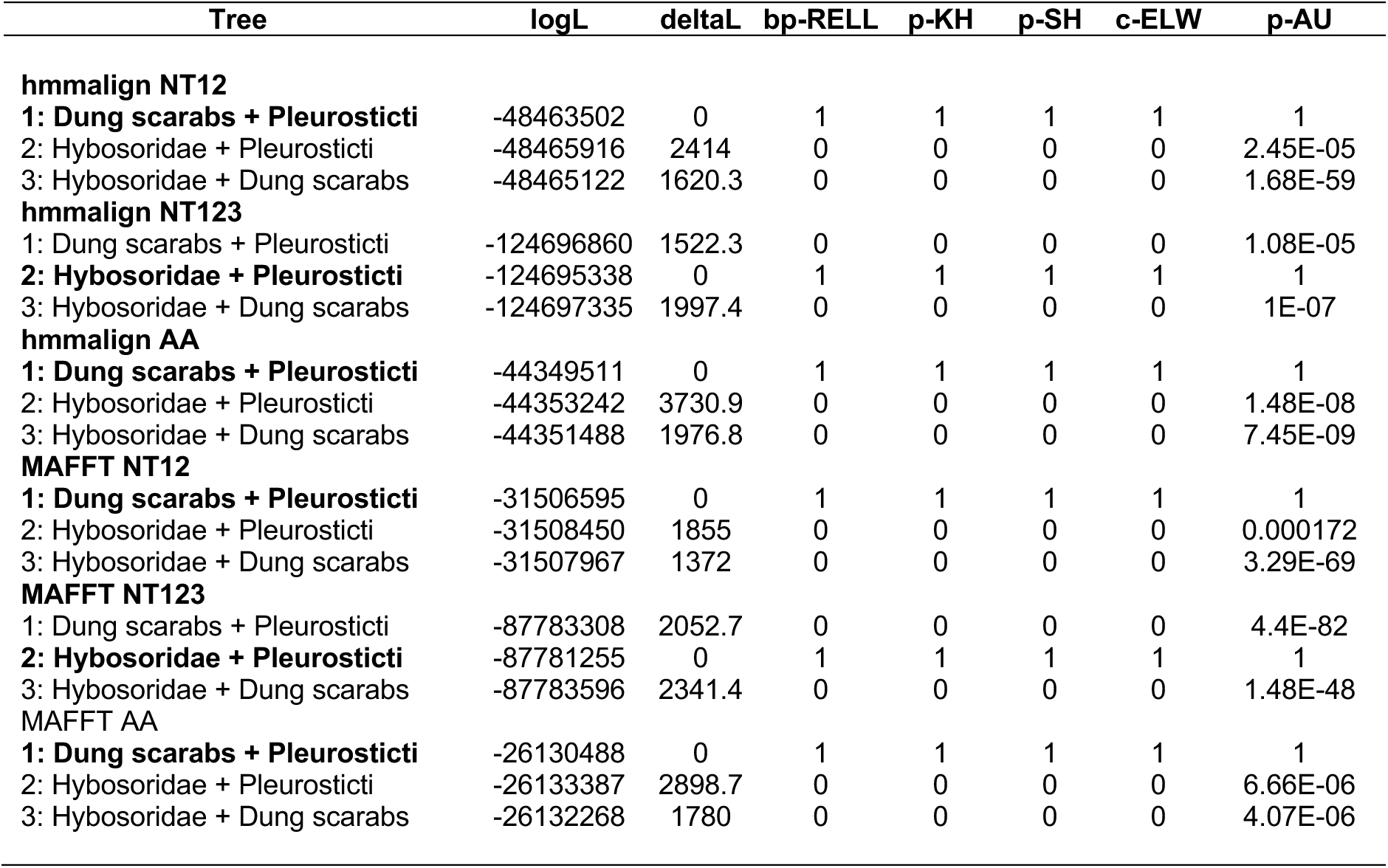
Likelihood tests regarding the monophyly of Scarabaeidae using the coleopteran-specific dataset, with constraint trees using a variety of resampling tests in IQ-TREE using the RELL approximation including bootstrap proportion (BP), Kishino-Hasegawa test (KH), Shimodaira-Hasegawa test (SH), expected likelihood weights (ELW), and the approximately unbiased (AU) test. The confirmed most likely topology is highlighted in bold.

| Tree | logL | deltaL | bp-RELL | p-KH | p-SH | c-ELW | p-AU |
| --- | --- | --- | --- | --- | --- | --- | --- |
| **hmmalign NT12** |  |  |  |  |  |  |  |
| **1: Dung scarabs + Pleurosticti** | -48463502 | 0 | 1 | 1 | 1 | 1 | 1 |
| 2: Hybosoridae + Pleurosticti | -48465916 | 2414 | 0 | 0 | 0 | 0 | 2.45E-05 |
| 3: Hybosoridae + Dung scarabs | -48465122 | 1620.3 | 0 | 0 | 0 | 0 | 1.68E-59 |
| **hmmalign NT123** |  |  |  |  |  |  |  |
| 1: Dung scarabs + Pleurosticti | -124696860 | 1522.3 | 0 | 0 | 0 | 0 | 1.08E-05 |
| **2: Hybosoridae + Pleurosticti** | -124695338 | 0 | 1 | 1 | 1 | 1 | 1 |
| 3: Hybosoridae + Dung scarabs | -124697335 | 1997.4 | 0 | 0 | 0 | 0 | 1E-07 |
| **hmmalign AA** |  |  |  |  |  |  |  |
| **1: Dung scarabs + Pleurosticti** | -44349511 | 0 | 1 | 1 | 1 | 1 | 1 |
| 2: Hybosoridae + Pleurosticti | -44353242 | 3730.9 | 0 | 0 | 0 | 0 | 1.48E-08 |
| 3: Hybosoridae + Dung scarabs | -44351488 | 1976.8 | 0 | 0 | 0 | 0 | 7.45E-09 |
| **MAFFT NT12** |  |  |  |  |  |  |  |
| **1: Dung scarabs + Pleurosticti** | -31506595 | 0 | 1 | 1 | 1 | 1 | 1 |
| 2: Hybosoridae + Pleurosticti | -31508450 | 1855 | 0 | 0 | 0 | 0 | 0.000172 |
| 3: Hybosoridae + Dung scarabs | -31507967 | 1372 | 0 | 0 | 0 | 0 | 3.29E-69 |
| **MAFFT NT123** |  |  |  |  |  |  |  |
| 1: Dung scarabs + Pleurosticti | -87783308 | 2052.7 | 0 | 0 | 0 | 0 | 4.4E-82 |
| **2: Hybosoridae + Pleurosticti** | -87781255 | 0 | 1 | 1 | 1 | 1 | 1 |
| 3: Hybosoridae + Dung scarabs | -87783596 | 2341.4 | 0 | 0 | 0 | 0 | 1.48E-48 |
| **MAFFT AA** |  |  |  |  |  |  |  |
| **1: Dung scarabs + Pleurosticti** | -26130488 | 0 | 1 | 1 | 1 | 1 | 1 |
| 2: Hybosoridae + Pleurosticti | -26133387 | 2898.7 | 0 | 0 | 0 | 0 | 6.66E-06 |
| 3: Hybosoridae + Dung scarabs | -26132268 | 1780 | 0 | 0 | 0 | 0 | 4.07E-06 |

**Table 3.** Likelihood tests regarding the monophyly of Rutelinae using the coleopteran-specific dataset, with constraint trees using a variety of resampling tests in IQ-TREE using the RELL approximation including bootstrap proportion (BP), Kishino-Hasegawa test (KH), Shimodaira-Hasegawa test (SH), expected likelihood weights (ELW), and the approximately unbiased (AU) test. The confirmed most likely topology is highlighted in bold.

| Tree | logL | deltaL | bp-RELL | p-KH | p-SH | c-ELW | p-AU |
| --- | --- | --- | --- | --- | --- | --- | --- |
| **hmmalign NT12** |  |  |  |  |  |  |  |
| **1: Adoretus + Anomala** | -48463550 | 0 | 0.986 | 0.983 | 1 | 0.986 | 0.991 |
| 2: Adoretus + Dynastinae | -48463962 | 412.1 | 0.0006 | 0.0012 | 0.0012 | 0.000607 | 0.0013 |
| 3: Anomala + Dynastinae | -48463842 | 292.12 | 0.0134 | 0.0168 | 0.0277 | 0.0135 | 0.0129 |
| **hmmalign NT123** |  |  |  |  |  |  |  |
| **1: Adoretus + Anomala** | -124695262 | 0 | 0.97 | 0.976 | 1 | 0.97 | 0.991 |
| 2: Adoretus + Dynastinae | -124695650 | 387.47 | 0.0053 | 0.0063 | 0.0129 | 0.00534 | 0.00767 |
| 3: Anomala + Dynastinae | -124695590 | 327.26 | 0.0244 | 0.0236 | 0.0407 | 0.0245 | 0.0198 |
| **hmmalign AA** |  |  |  |  |  |  |  |
| **1: Adoretus + Anomala** | -44349512 | 0 | 1 | 1 | 1 | 1 | 1 |
| 2: Adoretus + Dynastinae | -44349994 | 482.06 | 0.0002 | 0.0001 | 0.0002 | 0.000195 | 0.000127 |
| 3: Anomala + Dynastinae | -44350021 | 508.52 | 0.0002 | 0 | 0.0002 | 0.00019 | 0.000164 |
| **MAFFT NT12** |  |  |  |  |  |  |  |
| **1: Adoretus + Anomala** | -31506594 | 0 | 1 | 1 | 1 | 1 | 1 |
| 2: Adoretus + Dynastinae | -31507065 | 470.61 | 0.0001 | 0.0002 | 0.0003 | 0.0001 | 7.03E-05 |
| 3: Anomala + Dynastinae | -31507098 | 503.3 | 0 | 0 | 0 | 5.19E-05 | 0.000247 |
| **MAFFT NT123** |  |  |  |  |  |  |  |
| **1: Adoretus + Anomala** | -87781254 | 0 | 0.998 | 0.997 | 1 | 0.998 | 0.998 |
| 2: Adoretus + Dynastinae | -87781673 | 419.51 | 0.0021 | 0.0031 | 0.0048 | 0.00214 | 0.0029 |
| 3: Anomala + Dynastinae | -87781779 | 524.79 | 0.0001 | 0.0002 | 0.0005 | 0.00013 | 0.000376 |
| **MAFFT AA** |  |  |  |  |  |  |  |
| 1: Adoretus + Anomala | -26130489 | 0 | 1 | 1 | 1 | 1 | 1 |
| **2: Adoretus + Dynastinae** | -26130951 | 462.27 | 0.0003 | 0.0001 | 0.0005 | 0.000317 | 0.000697 |
| 3: Anomala + Dynastinae | -26130995 | 506.31 | 0.0002 | 0.0001 | 0.0002 | 0.000164 | 0.000818 |

**Table 4.** Likelihood tests regarding the monophyly sister group relationship of Hopliini and Melolonthinae s. str. (including Melolonthini and Sparrmannia, excluding Pachypus and Sericini) using the coleopteran-specific dataset, with constraint trees using a variety of resampling tests in IQ-TREE using the RELL approximation including bootstrap proportion (BP), Kishino-Hasegawa test (KH), Shimodaira-Hasegawa test (SH), expected likelihood weights (ELW), and the approximately unbiased (AU) test. The confirmed most likely topology is highlighted in bold.

| Tree | logL | deltaL | bp-RELL | p-KH | p-SH | c-ELW | p-AU |
| --- | --- | --- | --- | --- | --- | --- | --- |
| **hmmalign NT12** |  |  |  |  |  |  |  |
| 1: **Hopliini + Dynastinae etc.** | -48463522 | 0 | 1 | 1 | 1 | 1 | 1 |
| 2: Melolonthinae + Dynastinae etc. | -48466907 | 3384.2 | 0 | 0 | 0 | 0 | 9.98E-06 |
| 3: Melolonthinae + Hopliini | -48466288 | 2765.9 | 0 | 0 | 0 | 0 | 9.98E-06 |
| **hmmalign NT123** |  |  |  |  |  |  |  |
| 1: **Hopliini + Dynastinae etc.** | -124695344 | 0 | 1 | 1 | 1 | 1 | 1 |
| 2: Melolonthinae + Dynastinae etc. | -124703111 | 7766.7 | 0 | 0 | 0 | 0 | 1.01E-08 |
| 3: Melolonthinae + Hopliini | -124701830 | 6486 | 0 | 0 | 0 | 0 | 2.33E-43 |
| **hmmalign AA** |  |  |  |  |  |  |  |
| 1: **Hopliini + Dynastinae etc.** | -44349512 | 0 | 1 | 1 | 1 | 1 | 1 |
| 2: Melolonthinae + Dynastinae etc. | -44352720 | 3208.2 | 0 | 0 | 0 | 0 | 1.65E-52 |
| 3: Melolonthinae + Hopliini | -44352216 | 2704.6 | 0 | 0 | 0 | 0 | 1.22E-57 |
| **MAFFT NT12** |  |  |  |  |  |  |  |
| 1: **Hopliini + Dynastinae etc.** | -31506595 | 0 | 1 | 1 | 1 | 1 | 1 |
| 2: Melolonthinae + Dynastinae etc. | -31509134 | 2539.6 | 0 | 0 | 0 | 0 | 2.63E-44 |
| 3: Melolonthinae + Hopliini | -31508736 | 2141.3 | 0 | 0 | 0 | 0 | 1.64E-67 |
| **MAFFT NT123** |  |  |  |  |  |  |  |
| 1: **Hopliini + Dynastinae etc.** | -87781254 | 0 | 1 | 1 | 1 | 1 | 1 |
| 2: Melolonthinae + Dynastinae etc. | -87787291 | 6037.7 | 0 | 0 | 0 | 0 | 4.16E-50 |
| 3: Melolonthinae + Hopliini | -87786383 | 5129.7 | 0 | 0 | 0 | 0 | 4.65E-07 |
| **MAFFT AA** |  |  |  |  |  |  |  |
| 1: **Hopliini + Dynastinae etc.** | -26130487 | 0 | 1 | 1 | 1 | 1 | 1 |
| 2: Melolonthinae + Dynastinae etc. | -26132818 | 2331 | 0 | 0 | 0 | 0 | 1.09E-45 |
| 3: Melolonthinae + Hopliini | -26132612 | 2124.8 | 0 | 0 | 0 | 0 | 6.7E-09 |

Four-cluster likelihood mapping revealed support for a monophyly of Scarabaeidae when analyzing the NT12 and AA datasets (Fig. 3). When analyzing the NT12 datasets, a monophyly of Scarabaeidae was supported by 55–60% of the quartets. When analyzing the AA datasets, the support was > 80%. However, it should be noted that in all analyses almost all quartets that include the most remotely related outgroups *Helophorus* and *Ocypus* supported Hybosoridae + Pleurosticti, while among those containing the closest outgroup *Eulasia* (Glaphyridae), more than 90% supported monophyly of Scarabaeidae, even when analyzing the NT12 datasets. When analyzing the NT123 datasets, more than 70% of quartets supported Hybosoridae + Pleurosticti.

**Figure 3.**
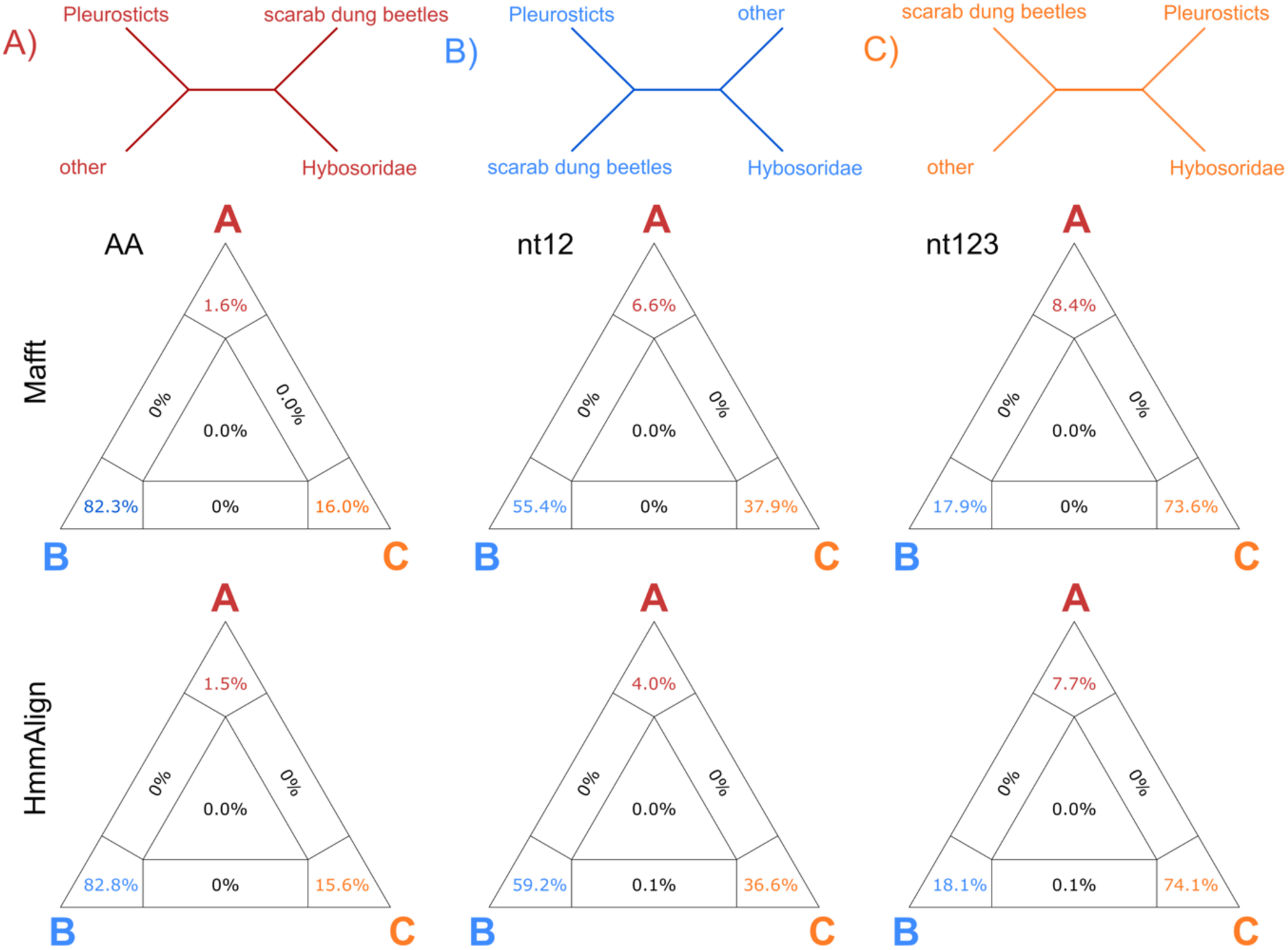
Quartet comparison of Coleoptera - single copy orthologs based on alignments with hmmalign and Mafft, both based on the AA, nt12, and nt123 data set.

## Discussion

The interfamilial results found in our study agree well with those of other multi-gene phylogenetic studies with less taxon and/or gene sampling (i.e., McKenna et al. 2019; Zhang et al. 2018; Cai et al. 2022). One exception is the position of Passalidae, which other studies found to be phylogenetically closely related to Geotrupidae + Bolboceratidae (McKenna et al. 2019; Beza-Beza et al. 2020). We found this phylogenetic position only in a few of our trees. In the majority of the inferred trees, we found Passalidae to be closely related to Scarabaeidae. As Passalidae had been represented by only few taxa on relatively long branches in all phylogenetic studies (including ours) conducted so far, a definitive statement on the phylogenetic position of Passalidae must await broader taxonomic sampling of the family itself and of related clades.

Our results confirm the early divergence of Orphninae within Pleurosticti, which had remained unresolved in some earlier molecular studies (e.g., Ahrens et al. 2014). However, an early divergence had been suspected based on morphological evidence (e.g., Browne & Scholtz 1998; Ahrens 2006). For example, Orphninae lack the derived conformation of spiracles after which Pleurosticti is named (Erichson 1848). The results of our study confirmed the sister group relationship of dung feeding scarabs and phytophagous pleurostict scarabs (i.e., monophyly of Scarabaeidae), which was not always found in previous studies with limited gene sampling (see Table 1). We found monophyly of Scarabaeidae with both supermatrix and coalescent-based tree reconstruction approaches. However, tree reconstruction results based on the nucleotide sequence data heavily depended on whether or not the 3^rd^ codon position was included — a phenomenon frequently observed in phylogenomic studies (Li et al. 2014).

The results of the likelihood mapping analyses strongly suggested that the non-monophyly of Scarabaeidae in the analyses of the NT123 data is an artifact. As this result is suggested primarily by quartets that include distantly related outgroup taxa, it can possibly be explained by long-branch attraction between the outgroup and the relatively long-branched dung-feeding scarabs. Likewise, long-branch attraction between the CRD clade and outgroups may have led to an artificial grouping of Melolonthinae and Hopliini.

The inferred monophyly of Scarabaeidae supports the initial hypothesis that the radiation of angiosperms had primarily affected species diversity, diversity of feeding habits, and morphological disparity of only a single lineage of Scarabaeoidea. This sheds new light on possible causes for their successful diversification, which might be highly lineage related, possibly also in regard to genome-driven events (e.g., McKenna et al. 2019). The successful diversification of other major herbivorous beetle lineages, such as the Phytophaga (Curculionoidea + Chrysomeloidea), was attributed to the genomic presence of plant cell wall-degrading enzymes obtained from bacteria and fungi (McKenna et al. 2019). However, the presence of PCWDEs in Scarabaeoidea was only limited to GH1 and GH9 that were expected to occur in most beetle species (McKenna et al. 2019), therefore other mechanisms are likely responsible for the successful radiation of Scarabaeidae. Another important factor that promoted diversification with angiosperm plants could be the efficient management of endosymbionts by the species of Scarabaeidae. As one key innovation in this regard can be seen the development of female accessory glands at the end of the digestive and genital duct, which are present in Aphodiinae and pleurostict scarabs (Ahrens 2006).

Accessory glands are known to have an important function in the transmission of endosymbiont bacteria that are known, beyond their active part in cellulose digestion (Martin 1983; Martin et al. 1991), to assist the production of pheromones (Hoyt et al. 1971). The latter could have had an important impact on the improvement of chemical communication in these groups (Pacheco et al. 2022). It should be noted that accessory glands are absent in Scarabaeinae, but other mechanisms for transmission of endosymbiotic bacteria have been documented (Estes et al. 2013). Nevertheless, the confirmed monophyly of Scarabaeidae allows us to interpret the reduction of accessory glands in Scarabaeinae dung beetles as a true loss and not as a parallelism between pleurostict scarabs and Aphodiinae.

The diverse Mesozoic fossil record of Hybosoridae (Lu et al. 2022) still provides important information for dating the onset of scarab divergences, given the poor fossil record of the Mesozoic Scarabaeidae (Krell 2007), particularly in amber which often allows a more accurate classification and a more robust systematic placement of its fossils. While we confirm the sister-group relationship between Scarabaeidae and Hybosoridae, the limited taxonomic sampling of this study restricts our ability to accurately place most of the known fossils into the inferred phylogenetic tree. Therefore, we explicitly refrained from inferring divergence time estimates.

### Implications on the classification of Scarabaeidae

Our phylogenetic analyses on Scarabaeidae revealed that there is — from a phylogenetic point of view — no necessity to split Scarabaeidae into two families. Given the monophyly of dung beetles and phytophagous pleurostict scarabs, such a splitting (Cherman & Morón 2014) would be rather arbitrary and in contrary to the aim of maintaining a stable nomenclature and classification, which are important backbones of all biodiversity-related databases (e.g., NCBI, GBIF). It should be noted, though, that our study did not include some taxonomic lineages (e.g., Ochodaeidae, Eremazinae) that earlier studies found (although with poor support) within a clade which included Glaphyridae, Hybosoridae, scarab dung beetles, and pleurostict Scarabaeidae (Ahrens et al. 2014; Neita-Moreno et al. 2019).

Interpretations for classification and evolution of the Scarabaeidae will be thus more robust when the position of these groups is better known.

The phylogenetic tree hypothesis supported by our study (Fig. 1) shows some examples of non-homogenous lineage classifications, in which sister taxa, according to the current classification, are either classified as tribes or as subfamilies (e.g., Pachypodini vs. Melolonthinae vs. Rutelinae/ Dynastinae/ Cetoniinae vs. Hopliini). Given the non-monophyly of Melolonthinae as currently understood (Smith 2006; Bouchard et al. 2011), the question arises to which clade the name “Melolonthinae” should be referred, with respective modifications to the current classification. Many unanswered questions remain due to the limited sampling here and contradictory tree topologies compared to and between previous studies (e.g., Ahrens et al. 2014; Eberle et al. 2019).

The clear phylogenetic separation of the monophyletic “Southern World” Melolonthinae and the clade containing Sericini, Ablaberini, and Diphucephalini from the remaining pleurostict scarabs (i.e., Cetoniinae, Rutelinae, Dynastinae, Melolonthinae etc.) makes it reasonable to treat both lineages as separate subfamilies. Both lineages are currently classified as a series of tribes within Melolonthinae (Smith 2006). A revised classification that elevates these clades to subfamilies would alleviate the problem, at least in part, of rendering Melolonthinae polyphyletic. It would also help to focus on the problem of whether Melolonthinae are monophyletic under inclusion of Hopliini and Macrodactylini (the latter not included in taxonomic sampling of this study). In regard to the sister group relationship Hopliini + Melolonthini, based on the current sampling there is good support that they do not form a monophyletic group (Table 4) although due to the limited sampling our results should not be regarded as fully conclusive.

Currently, the clade Melolonthini, which we refer here to a restricted interpretation of the subfamily “Melolonthinae”, includes several lineages currently circumscribed as subtribes, such as Rhizotrogina, Pyglina, Melolonthina, Enariina, Schizonychina, Leucopholina. It also includes several other minor lineages (Eberle et al. 2019) and could at least contain the genus *Sparrmannia*. The latter is so far assigned to Tanyproctini but the position of other genera of the polyphyletic Tanyproctini remains yet uncertain (Eberle et al. 2019). A restricted Melolonthinae (see Fig. 1) would be a starting point for a re-classification that would allow retaining well-established subfamily names (e.g., Dynastinae, Rutelinae, Cetoniinae). However, the exact extent of this clade Melolonthinae is yet to be identified, particularly with reference to the other lineages so far classified as “Melolonthinae” (e.g., Diplotaxini, Hopliini, Macrodactylini; e.g., Ahrens et al. 2014). To further address this topic, the taxonomic sampling needs to be extended, also to allow more robust statistical topology testing (Table 2-4; Fig. 3). The same applies to Rutelinae + Dynastinae: the monophyly of Rutelinae with respect to Dynastinae recovered by our results was often not supported in other studies (e.g., Ahrens & Vogler 2008; Ahrens et al. 2014; Šipek et al. 2016; Neita-Moreno et al. 2019), although Guo et al. (2022) also found a clade of Adoretini + Anomalini to the exclusion of Dynastinae using mitochondrial genomes, similar to our results. However, as our datasets contained only one representative each of two of the seven currently recognized tribes of Rutelinae, a decision on the classification of these two subfamilies should await further studies.

The following formal classification changes are proposed. Sericinae Kirby, 1837 **stat. rest.** and **sensu n.** is re-elevated to subfamily and revised to include the tribes Sericini, Ablaberini and Diphucephalini. Sericoidinae Erichson, 1847 **stat. rest.** and **sensu n.** is re-elevated to subfamily and revised to include the tribes Sericoidini, Liparetrini, Maechidiini, Heteronychini, Scitalini and Phyllotocini. Sericoidinae Erichson, 1847 has formally priority over the younger name Liparetrinae Burmeister, 1855 (and other tribal names within the lineage) and also includes the tribe Automoliini not included in this analysis but being confirmed in previous molecular phylogenies to be part of the same lineage (Ahrens & Vogler 2008; Ahrens et al. 2014). Based on morphological characters, other candidate members of this subfamily are Colymbomorphini, Comophorinini, Pachytrichini, and Phyllotocidiini, however, their phylogenetic placement has not been confirmed yet with molecular data.

Apart of some of our robust results regarding the monophyly of Scarabaeidae, we consider this work also as a primer and starting point for further and more detailed phylogenomic research in Scarabaeoidea. In this, the generated transcriptomic data will serve as backbone for other approaches such as DNA target enrichment approaches, which would possibly allow to considerably extend the taxon sampling. Because also dry museum specimens could be analyzed this way, we expect that even yet entirely obscure lineages (e.g., Dynamopodinae, Phaenomeridinae, Belohinidae), which never have been considered in any phylogenetic analysis, will find their place in a phylogenetic tree.

## Supplementary Material

**Supplement Files 1-12** (see Dryad, link: xxx; to be provided)

Supplement File 1: Perl script to filter Trinity contigs for the longest variant. (trinity_longest_d.pl).

Supplement File 2: Perl script to remove unaligned regions from hmmalign output. (hmmalign_cut2_d.pl).

Supplement File 3: Perl script to remove third codon positions from an alignment. (extract_codpos_d.pl).

Supplement File 4: Perl script to concatenate gene alignments into a partitioned superalignment. (concat_eogs_part_d.pl).

Supplement File 5: Perl script to remove alignment positions present in less than a given number of taxa. (removegaps_d.pl).

Supplement File 6: Perl script to calculate average pairwise identities between sequences within alignments. (pairwise_id2.pl).

Supplement File 7: Coalescent-based phylogenetic trees from ASTRAL analysis (astral_trees.zip) (see astral-trees-descriptions.txt for descriptions).

Supplement File 8: Concatenated sequence alignments of coding sequences of all loci in a dataset in FASTA format (concat_alignments.zip) (see concat-alignments-descriptions.txt for descriptions).

Supplement File 9: Maximum-likelihood phylogenetic trees created with IQ-TREE based on concatenated alignments of coding sequences of USCO loci (concat_trees.zip) (see concat-trees-descriptions.txt for descriptions).

Supplement File 10: Sequence alignments of coding sequences of individual loci in FASTA format (gene_alignments.zip) (see gene-alignments-descriptions.txt for descriptions).

Supplement File 11: Maximum-likelihood phylogenetic trees created with IQ-TREE for individual loci (gene_trees.zip) (see gene-trees-descriptions.txt for descriptions).

Supplement File 12: Partition files for concatenated gene alignments in NEXUS format (partition_files.zip) (see partition-files-descriptions.txt for descriptions).

## Acknowledgements

The study was funded by the German Research Foundation grants AH 175/6-1, AH175/6-2 (D.A.), MI 649/18-1 (B.M.), NI 1387/6-1 and 6-2 as well as by institutional funding of the Museum Koenig Bonn and the A. Koenig Stiftung Bonn.

---

**Figure S1.**
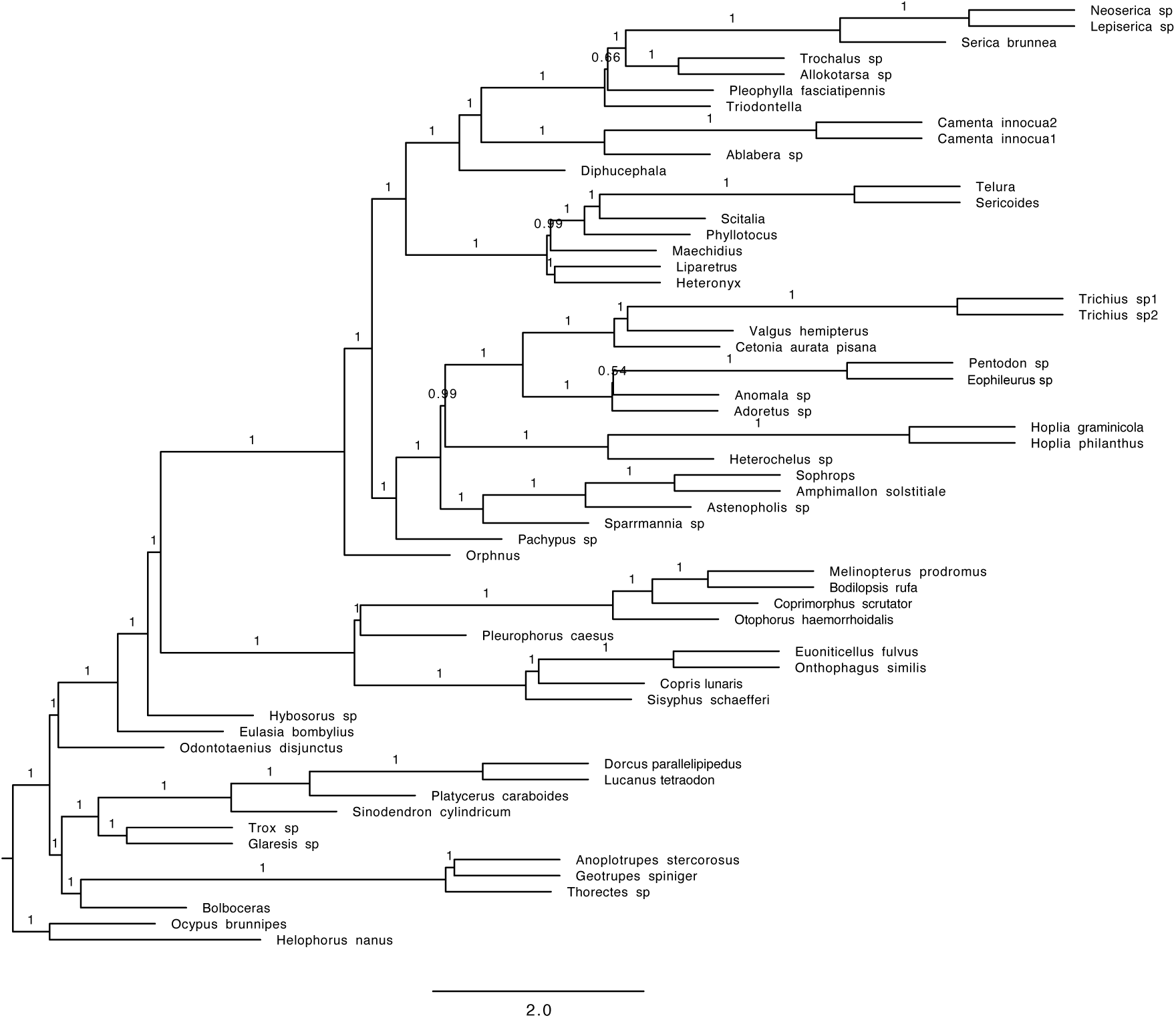
Phylogenetic tree obtained with the coalescent tree search with Astral and amino acid sequences (Mafft alignment; full dataset; Coleoptera orthologs).

**Figure S2.**
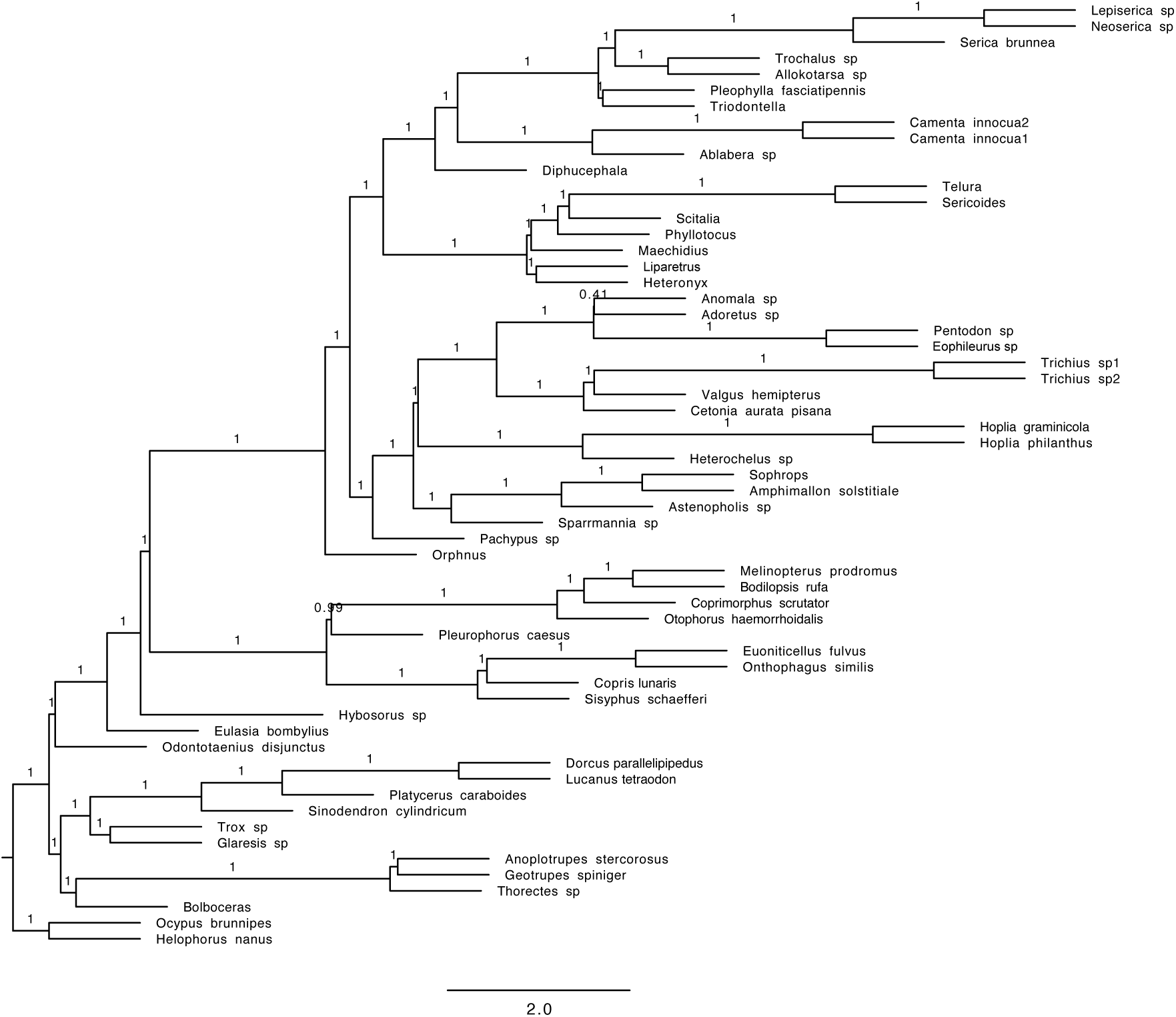
Phylogenetic tree obtained with the coalescent tree search with Astral and nucleotide data containing only the first and second base pair (nt12) (Mafft alignment; full dataset; Coleoptera orthologs).

**Figure S3.**
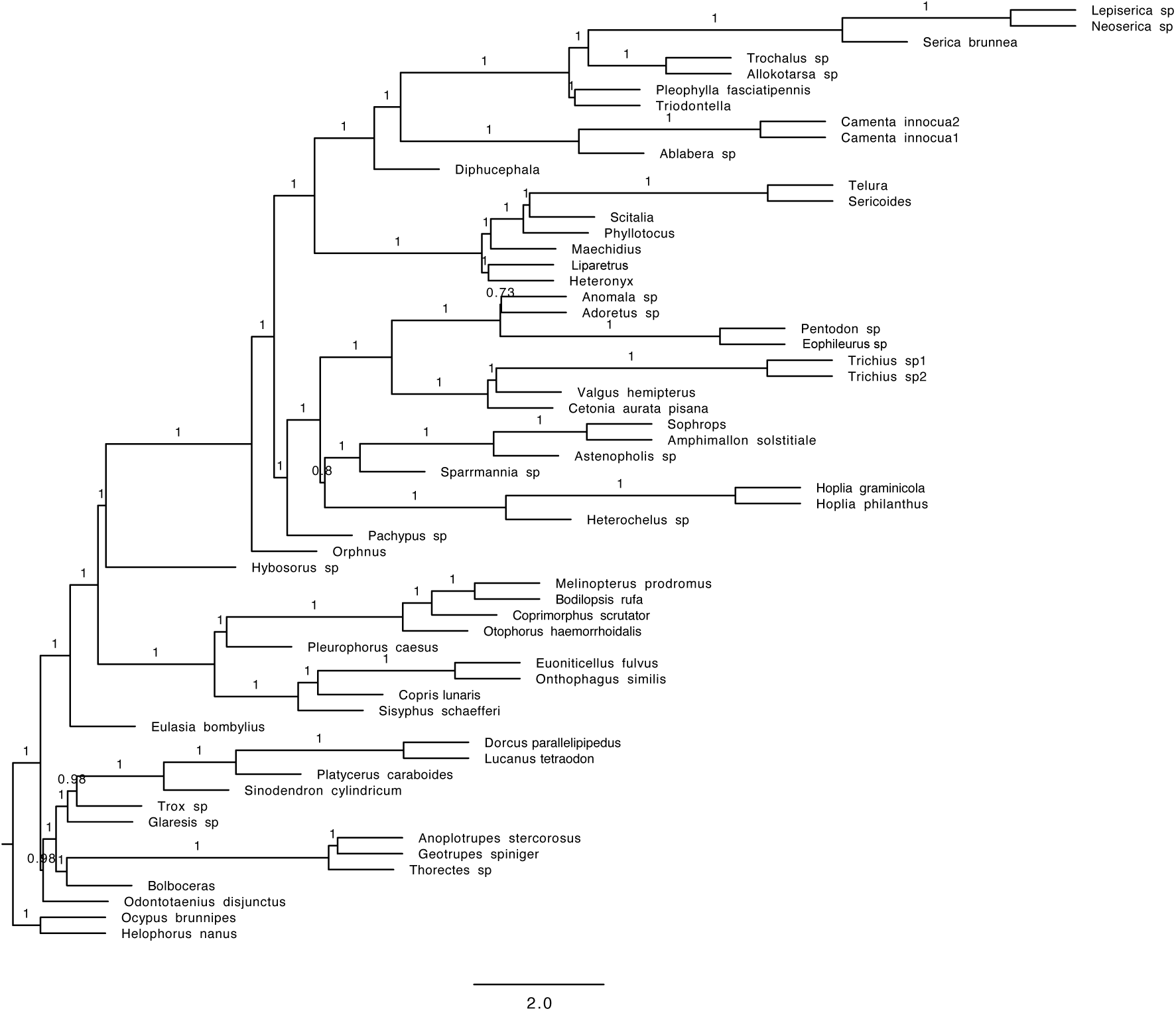
Phylogenetic tree obtained with the coalescent tree search with Astral and nucleotide data containing all three base pairs (nt123) (Mafft alignment; full dataset; Coleoptera orthologs).

**Figure S4.**
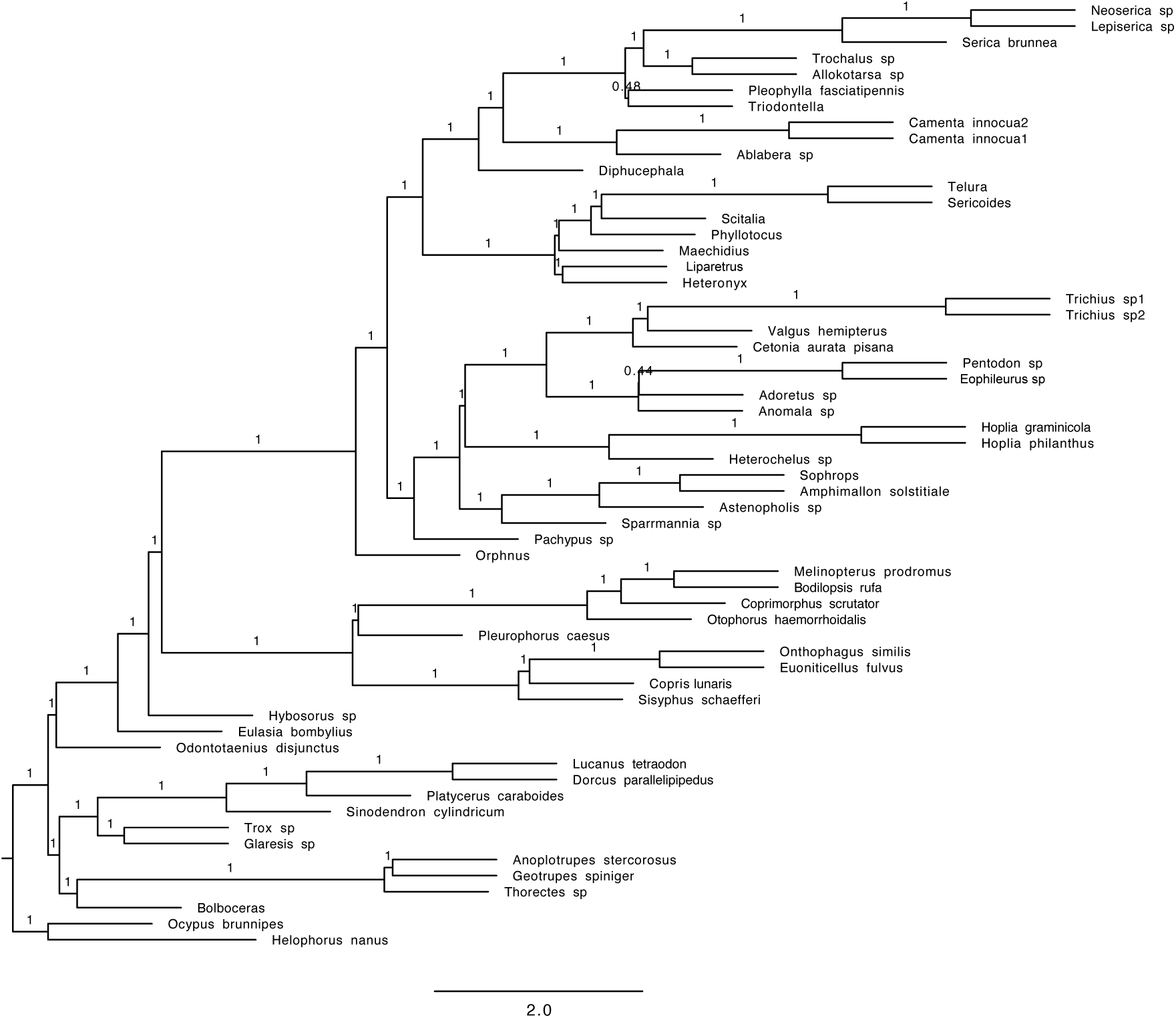
Phylogenetic tree obtained with the coalescent tree search with Astral and amino acid sequences (HmmAlign alignment; full dataset; Coleoptera orthologs).

**Figure S5.**
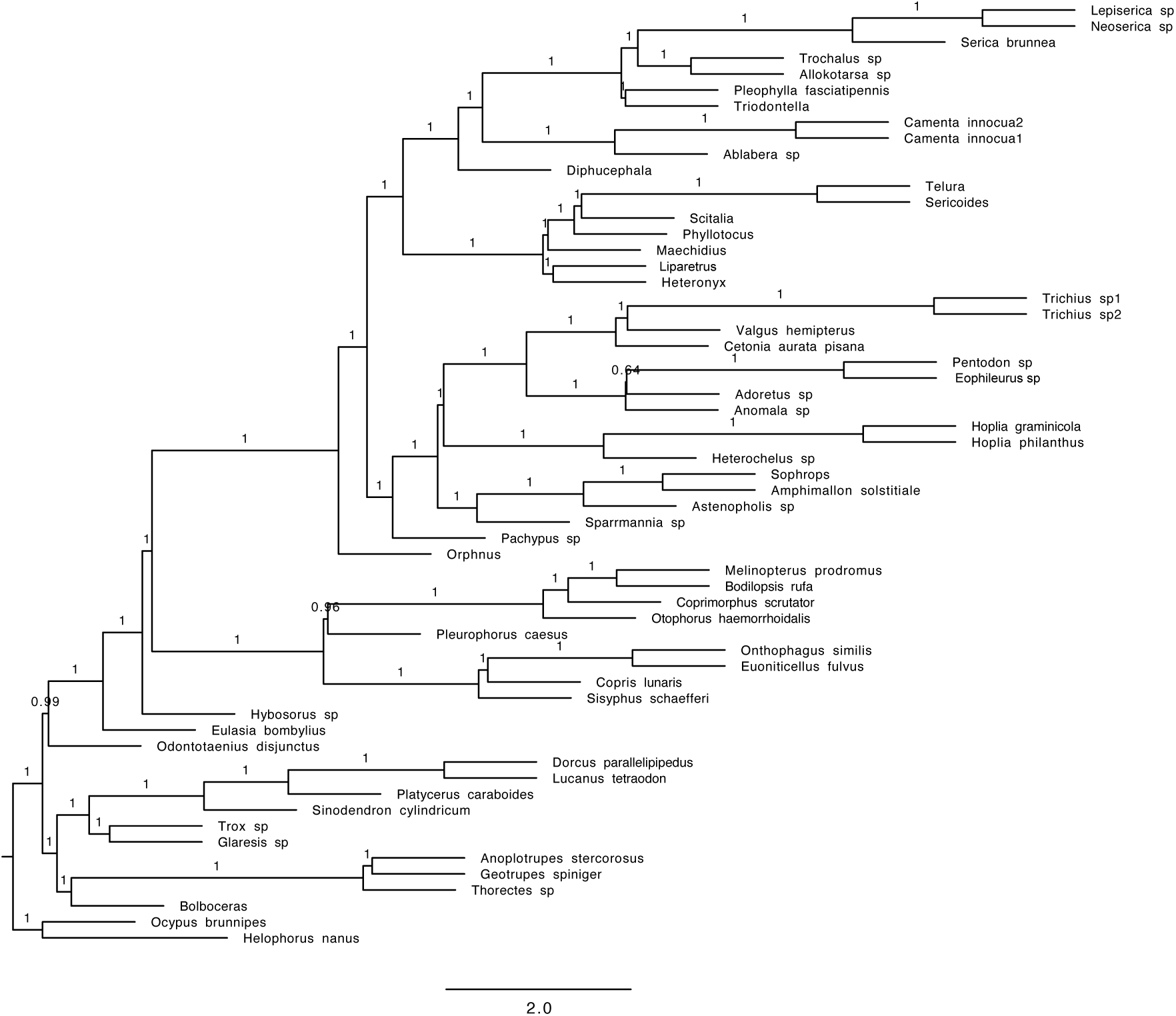
Phylogenetic tree obtained with the coalescent tree search with Astral and nucleotide data containing only the first and second base pair (nt12) (HmmAlign alignment; full dataset; Coleoptera orthologs).

**Figure S6.**
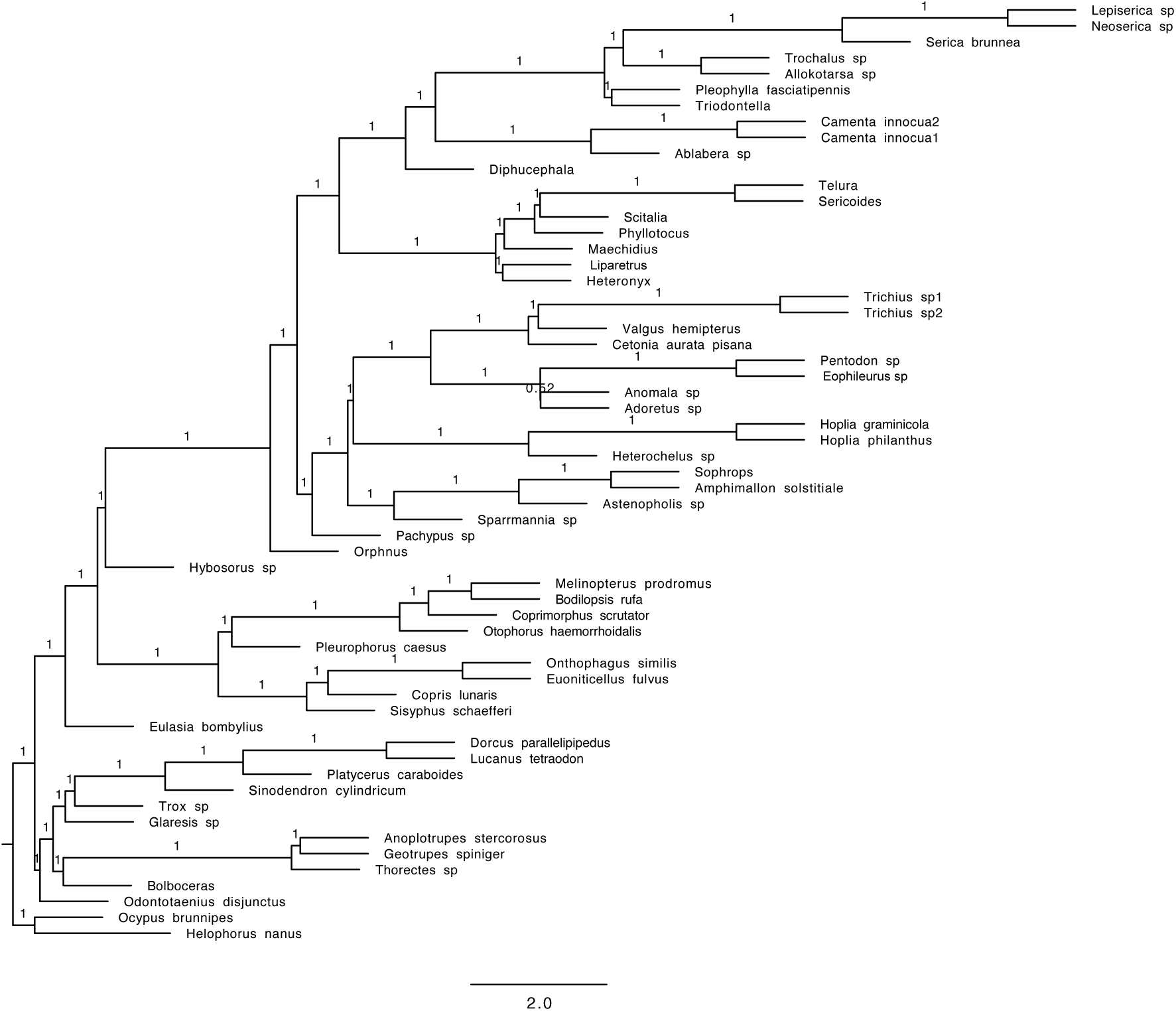
Phylogenetic tree obtained with the coalescent tree search with Astral and nucleotide data containing all three base pairs (nt123) (HmmAlign alignment; full dataset; Coleoptera orthologs).

**Figure S7.**
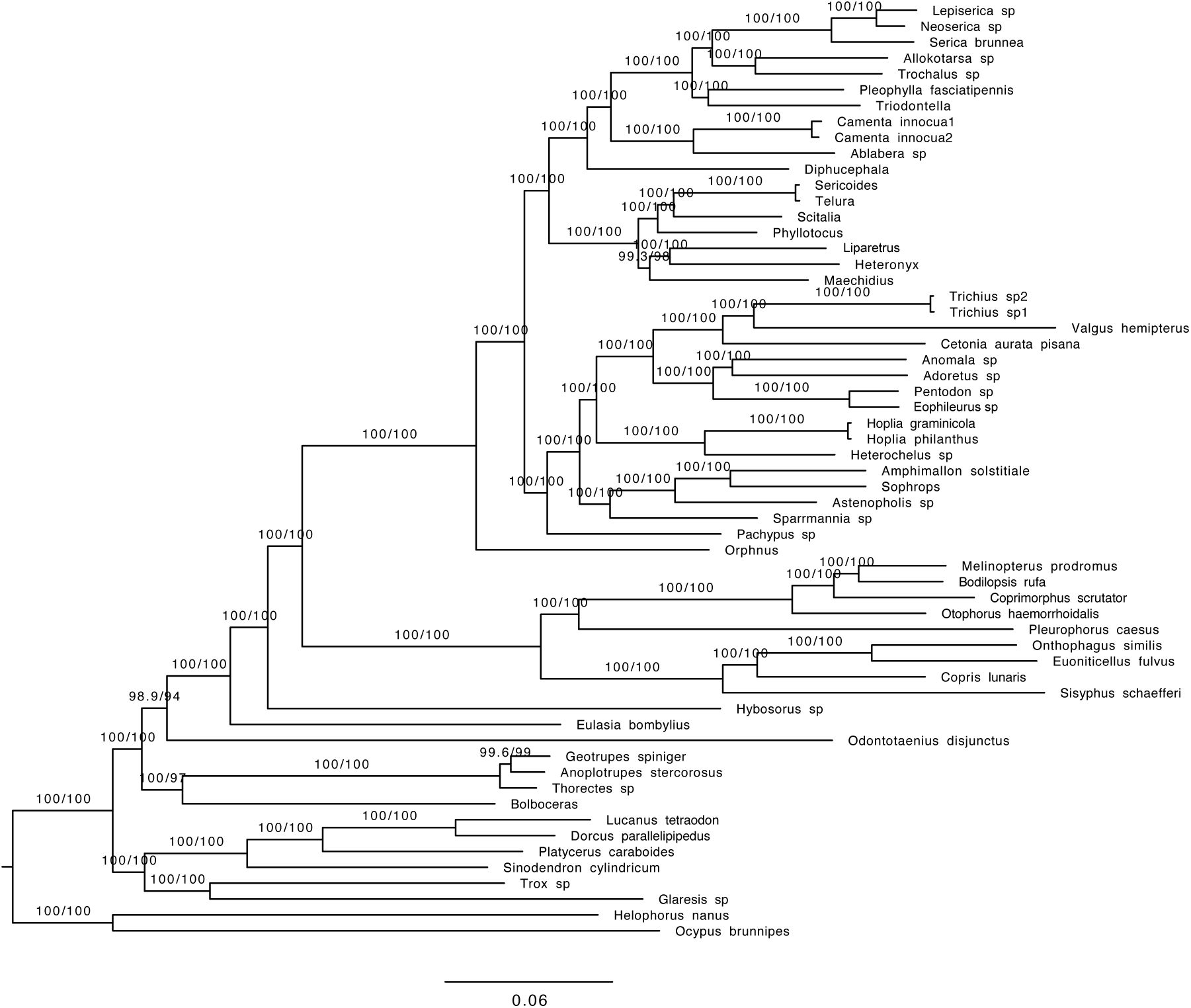
Phylogenetic tree obtained with the concatenated data and amino acid sequences (AA) (Mafft alignment; full dataset; Coleoptera orthologs).

**Figure S8.**
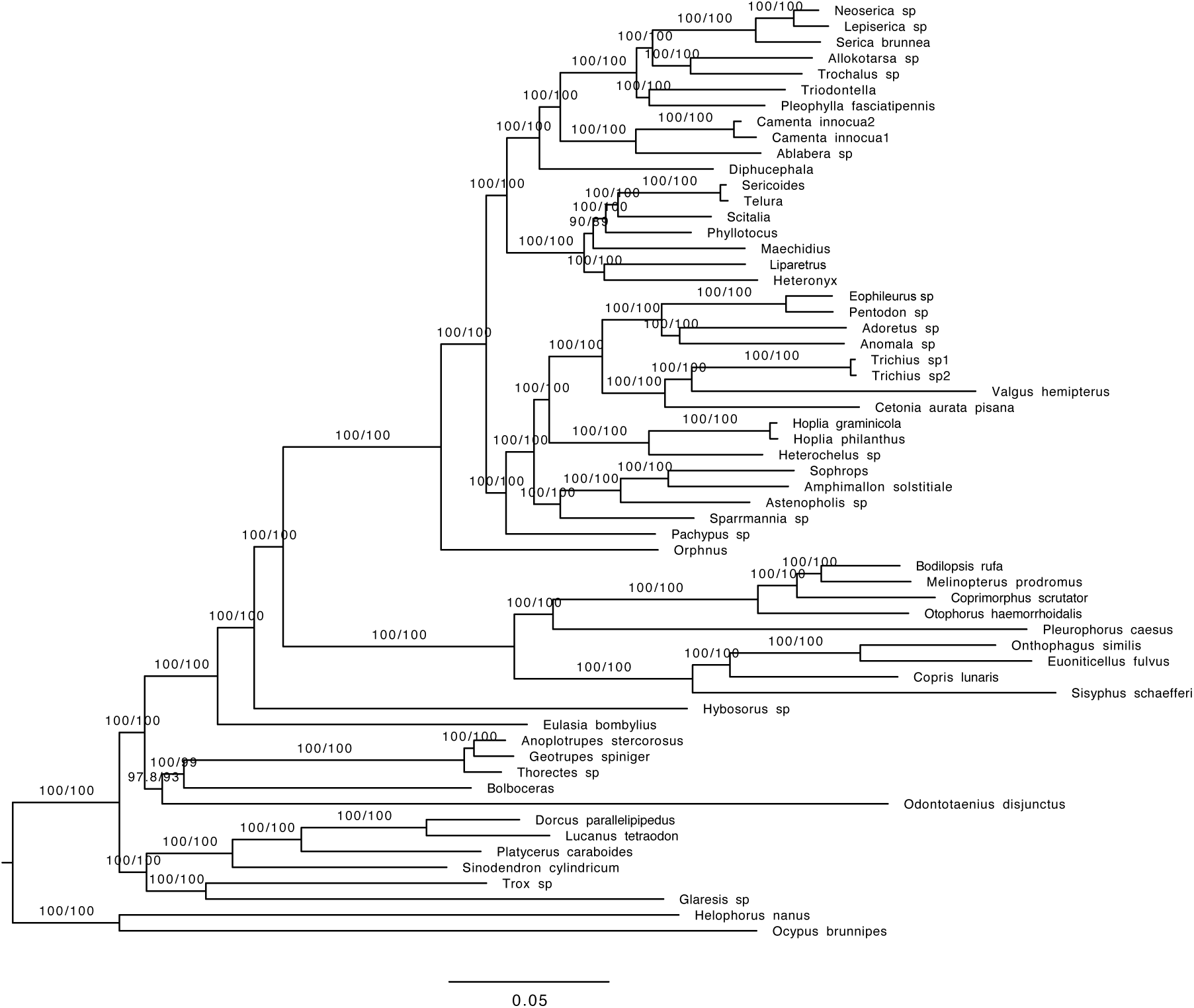
Phylogenetic tree obtained with the concatenated data and nucleotide data containing only the first and second base pair (nt12) (Mafft alignment; full dataset; Coleoptera orthologs).

**Figure S9.**
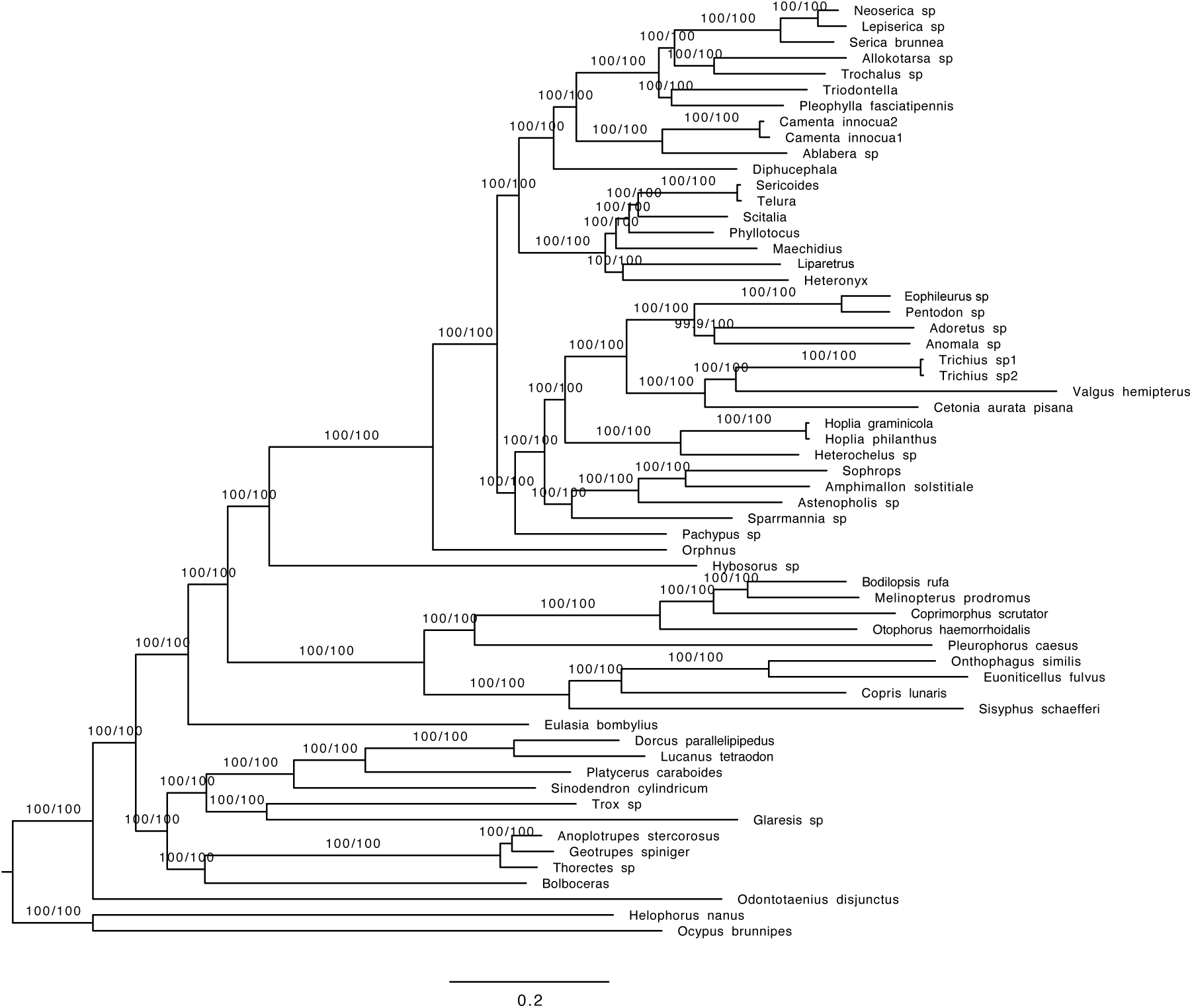
Phylogenetic tree obtained with the concatenated data and nucleotide data containing all three base pairs (nt123) (Mafft alignment; full dataset; Coleoptera orthologs).

**Figure S10.**
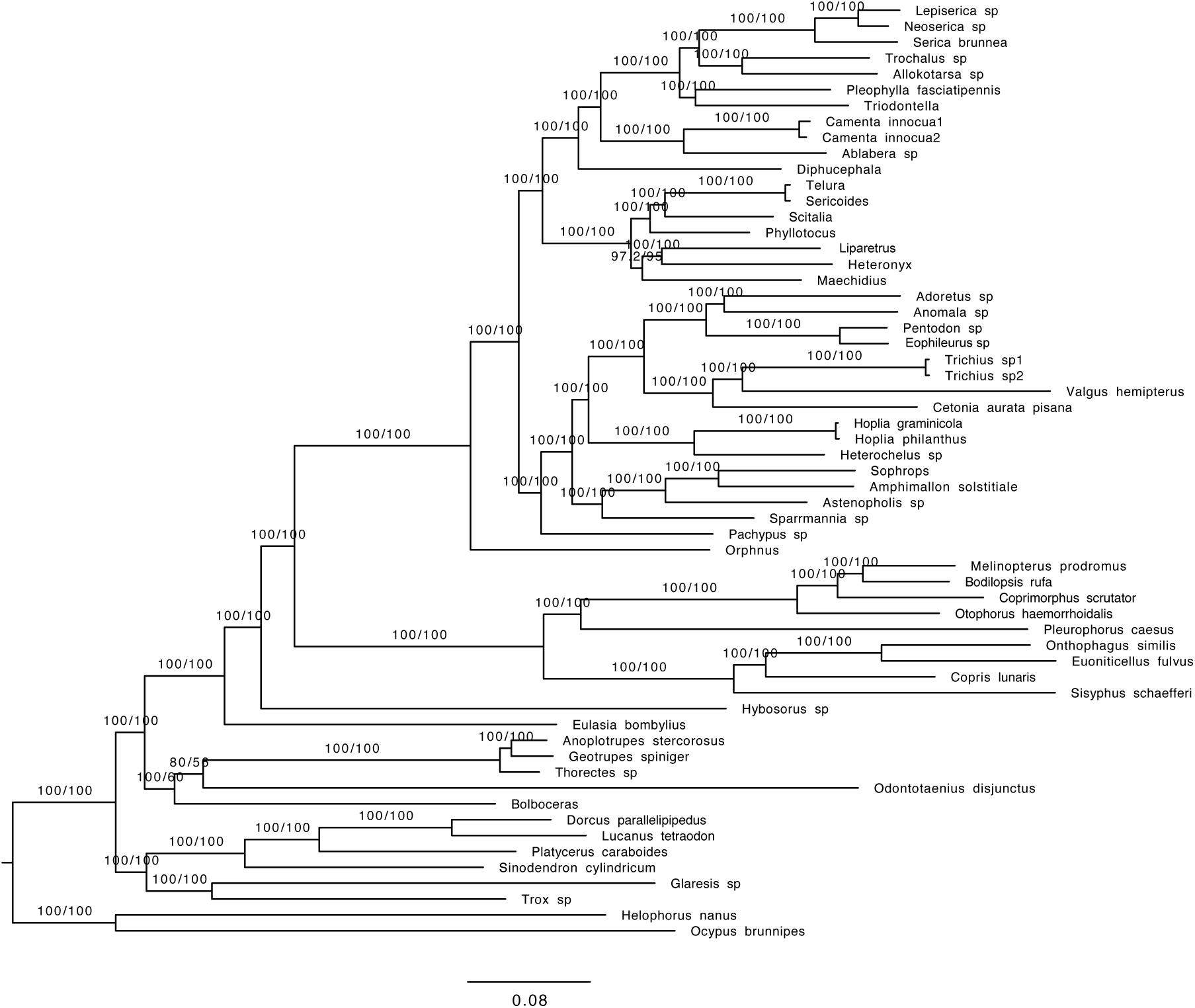
Phylogenetic tree obtained with the concatenated data and amino acid sequences (AA) (HmmAlign alignment; full dataset; Coleoptera orthologs).

**Figure S11.**
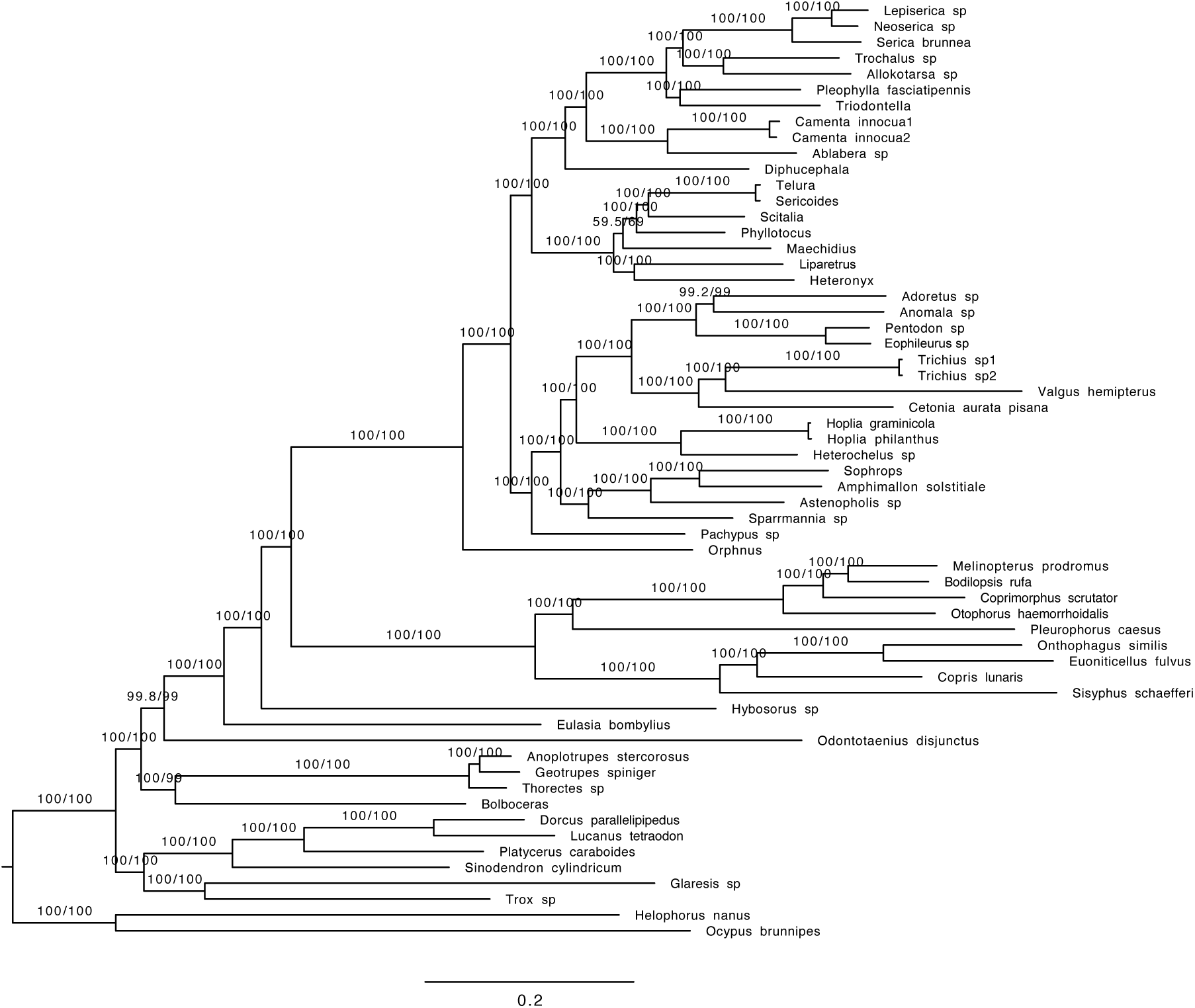
Phylogenetic tree obtained with the concatenated data and nucleotide data containing only the first and second base pair (nt12) (HmmAlign alignment; full dataset; Coleoptera orthologs).

**Figure S12.**
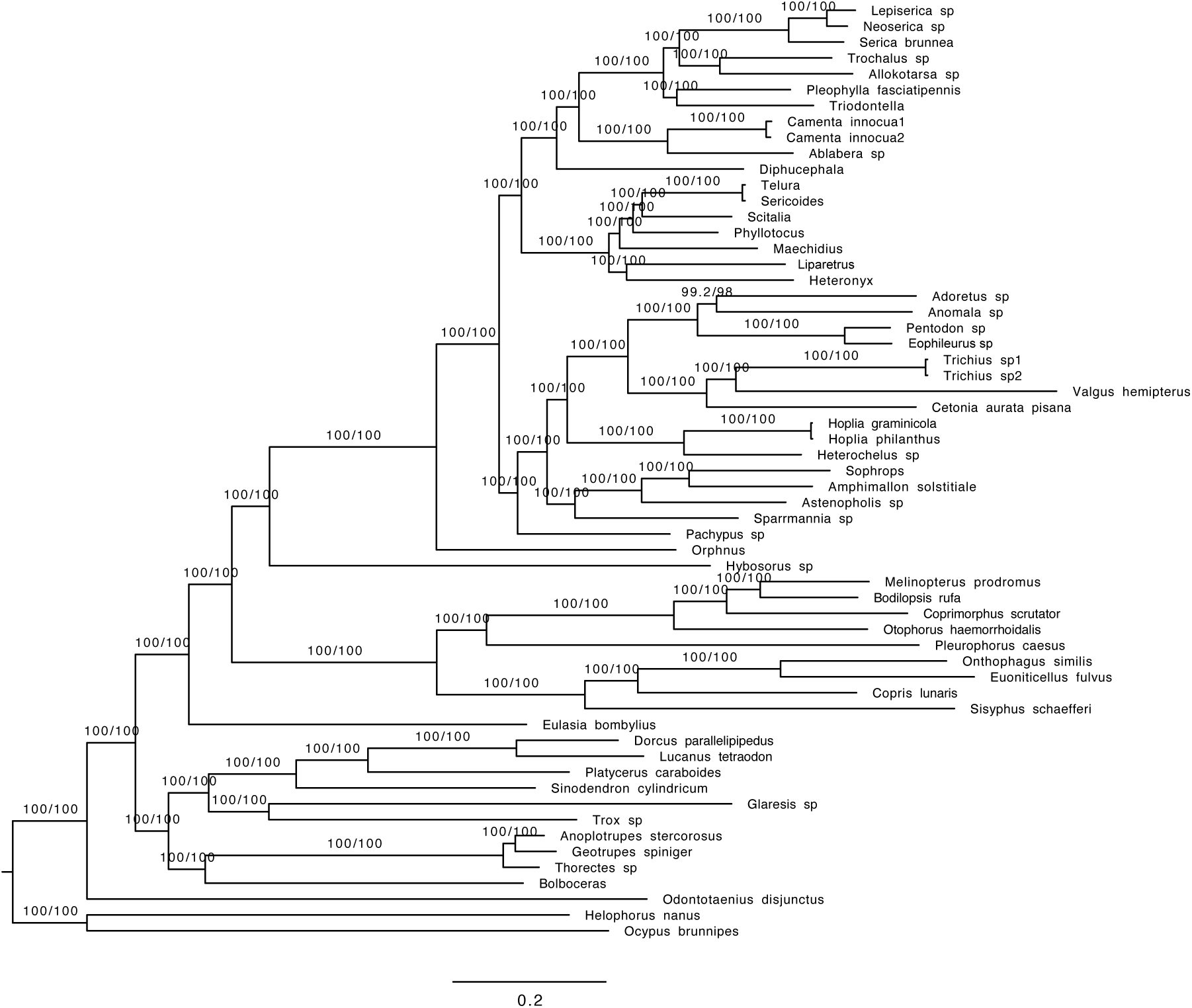
Phylogenetic tree obtained with the concatenated data and nucleotide data containing all three base pairs (nt123) (HmmAlign alignment; full dataset; Coleoptera orthologs).

**Figure S13.**
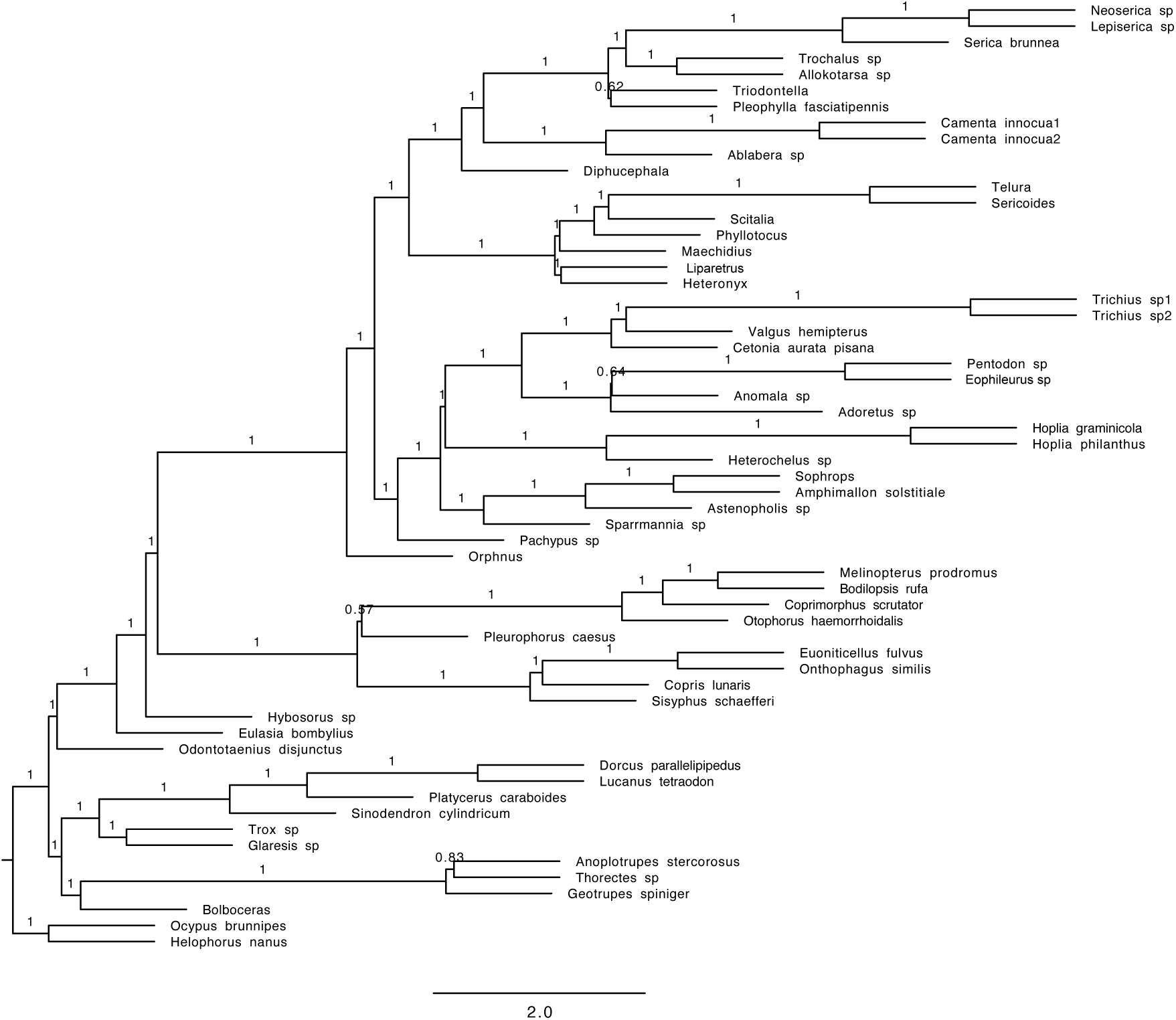
Phylogenetic tree obtained with the coalescent tree search with Astral and amino acid sequences (Mafft alignment; dataset with 70% complete data; Coleoptera orthologs).

**Figure S14.**
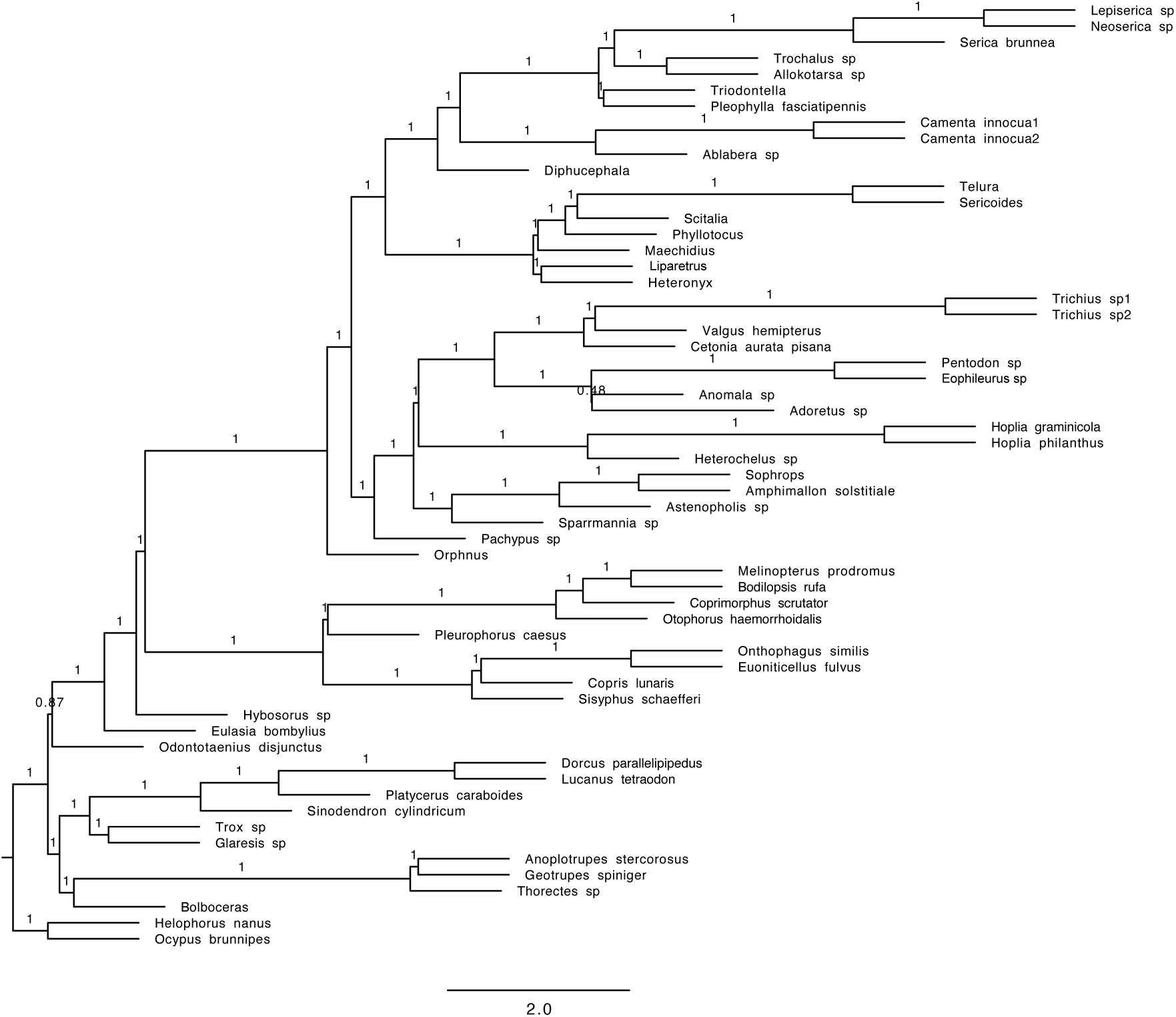
Phylogenetic tree obtained with the coalescent tree search with Astral and nucleotide data containing only the first and second base pair (nt12) (Mafft alignment; dataset with 70% complete data; Coleoptera orthologs).

**Figure S15.**
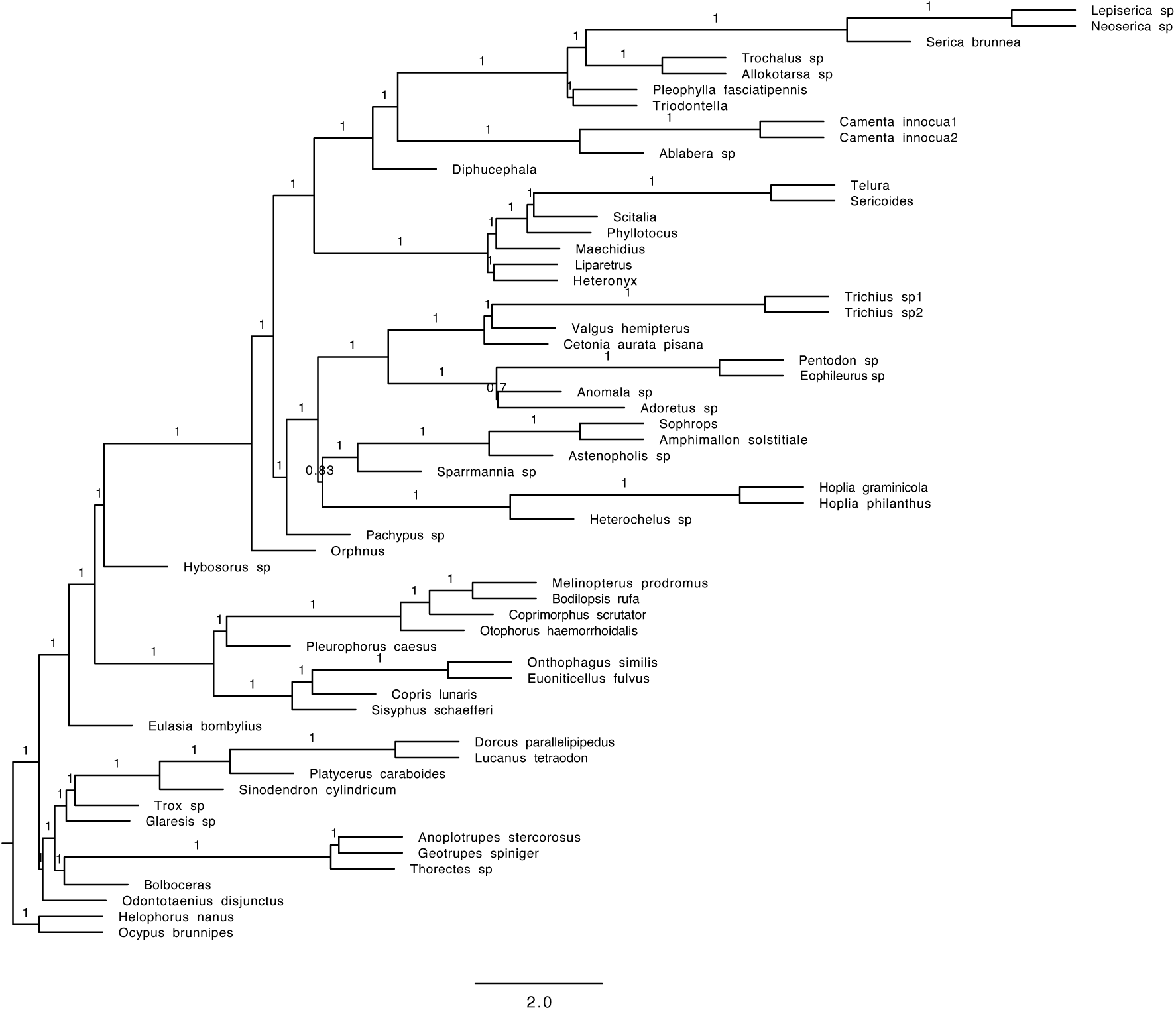
Phylogenetic tree obtained with the coalescent tree search with Astral and nucleotide data containing all three base pairs (nt123) (Mafft alignment; dataset with 70% complete data; Coleoptera orthologs).

**Figure S16.**
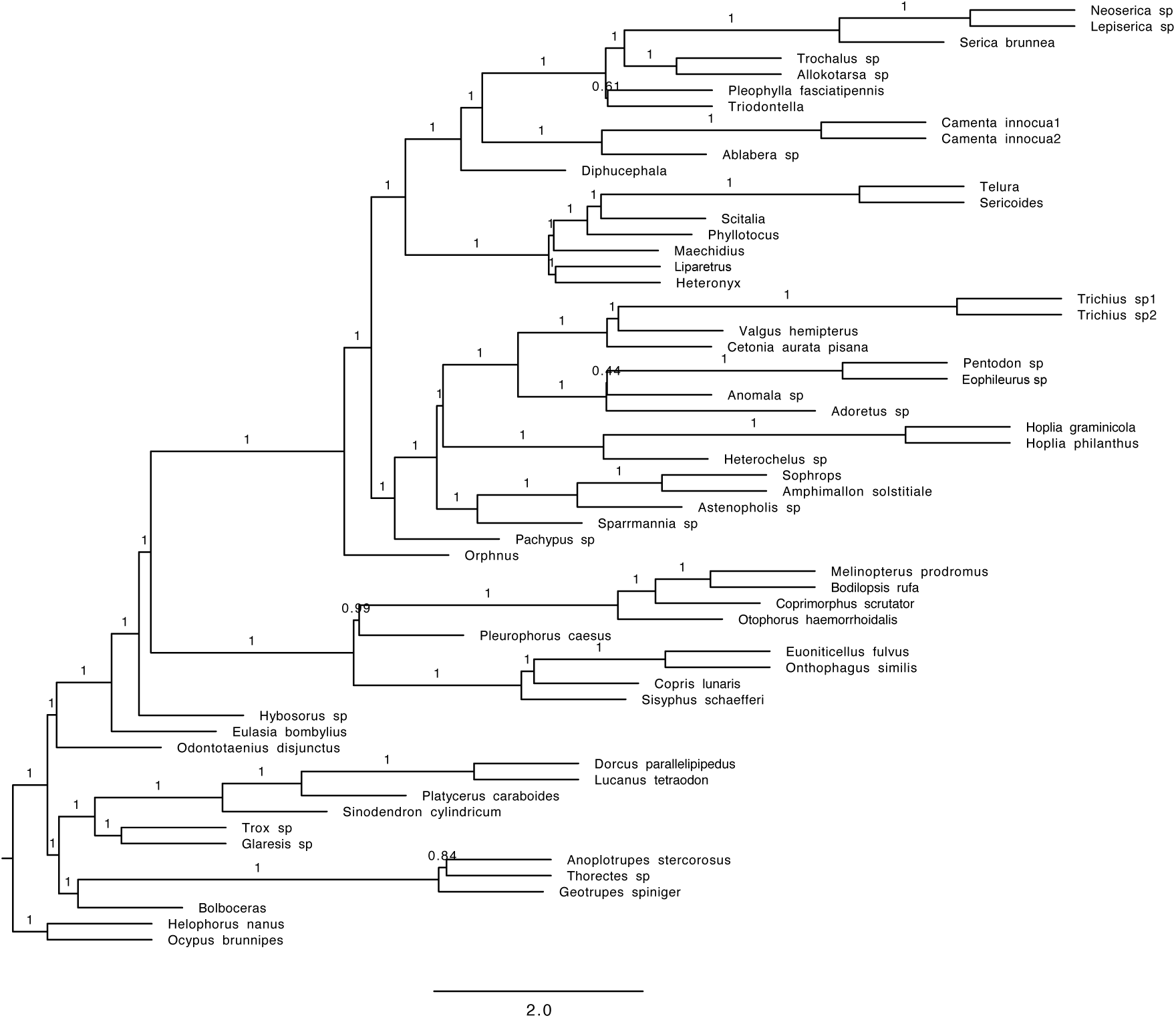
Phylogenetic tree obtained with the coalescent tree search with Astral and amino acid sequences (HmmAlign alignment; dataset with 70% complete data; Coleoptera orthologs).

**Figure S17.**
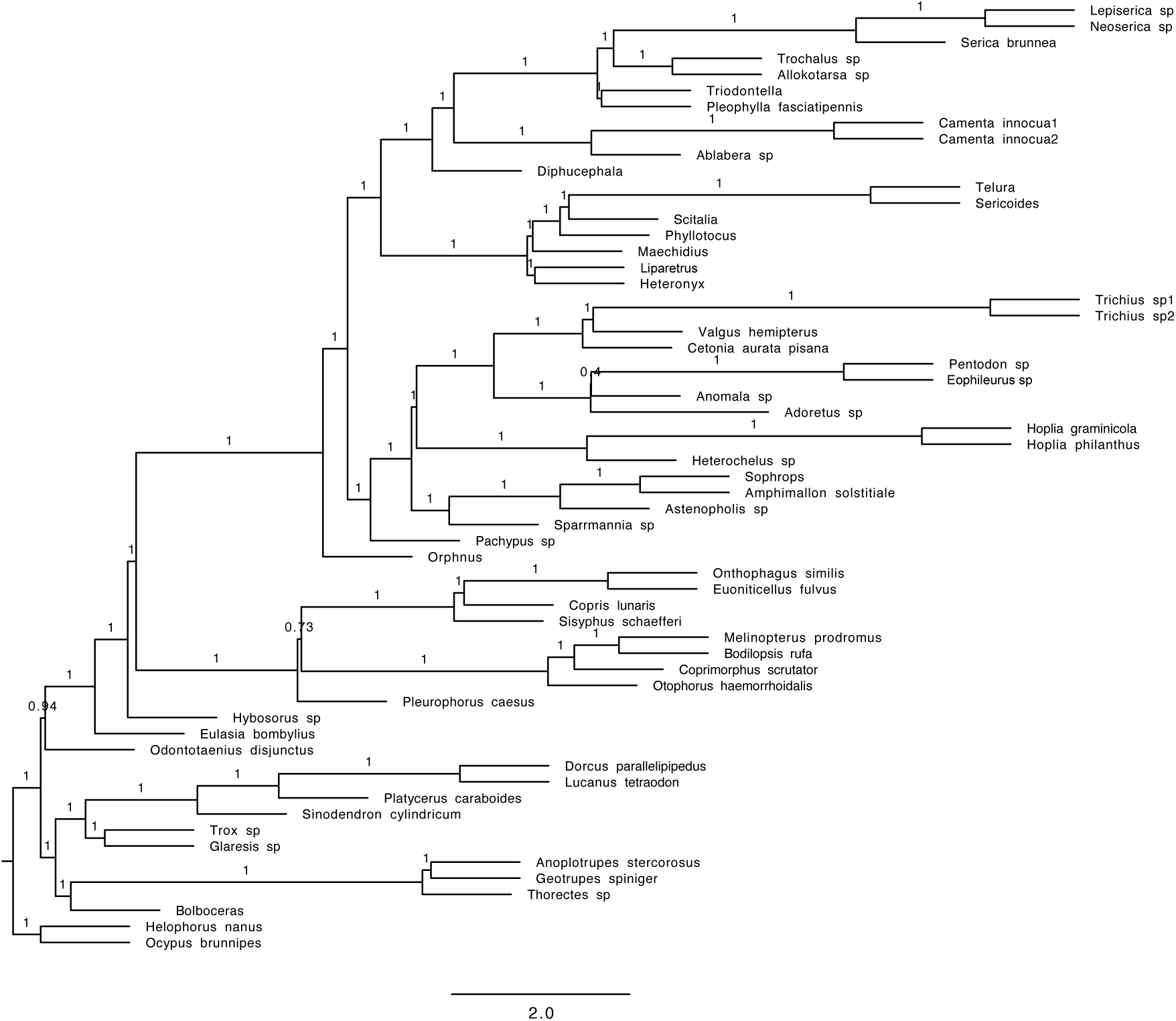
Phylogenetic tree obtained with the coalescent tree search with Astral and nucleotide data containing only the first and second base pair (nt12) (Mafft alignment; dataset with 70% complete data; Coleoptera orthologs).

**Figure S18.**
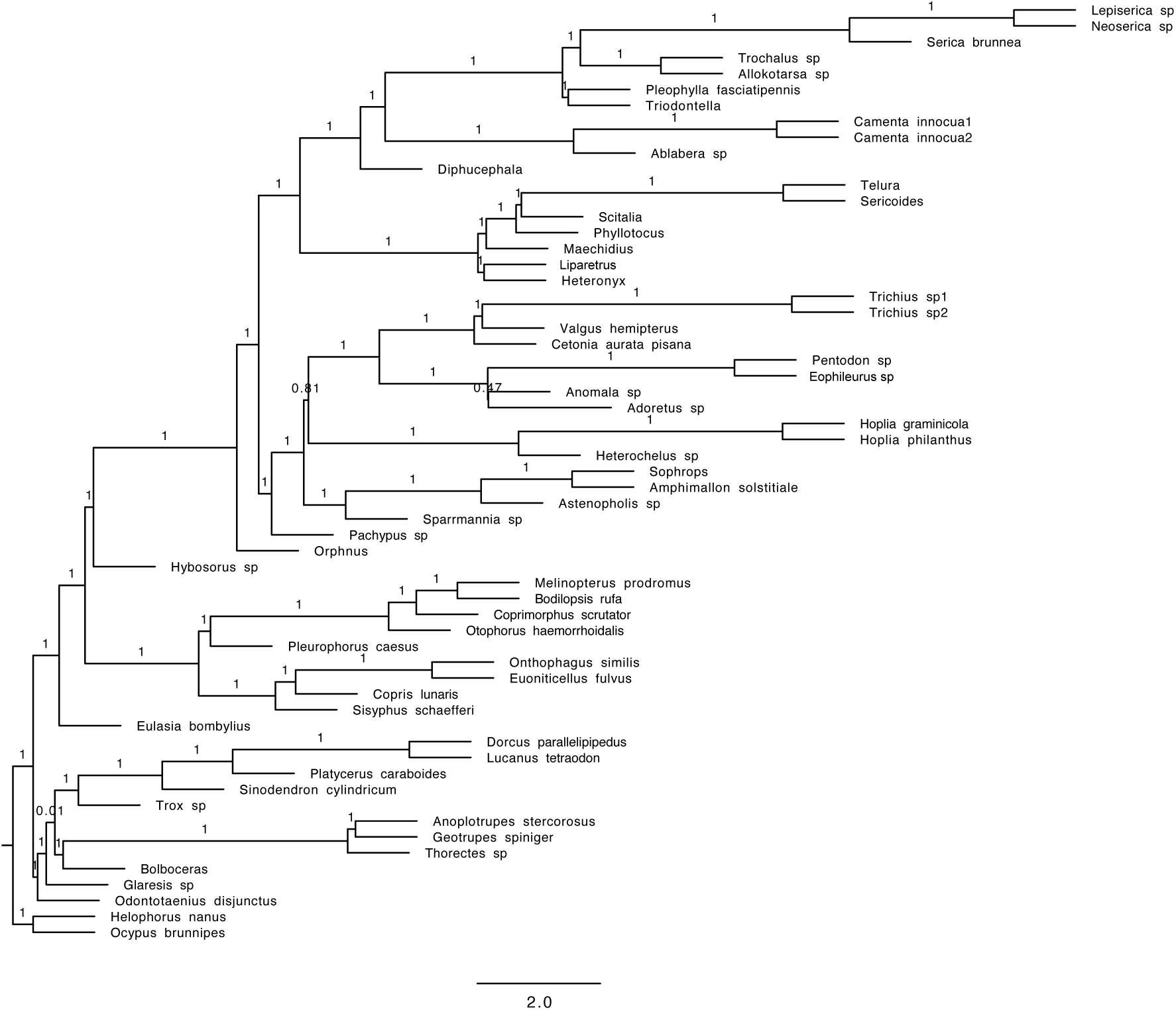
Phylogenetic tree obtained with the coalescent tree search with Astral and nucleotide data containing all three base pairs (nt123) (HmmAlign alignment; dataset with 70% complete data; Coleoptera orthologs).

**Figure S19.**
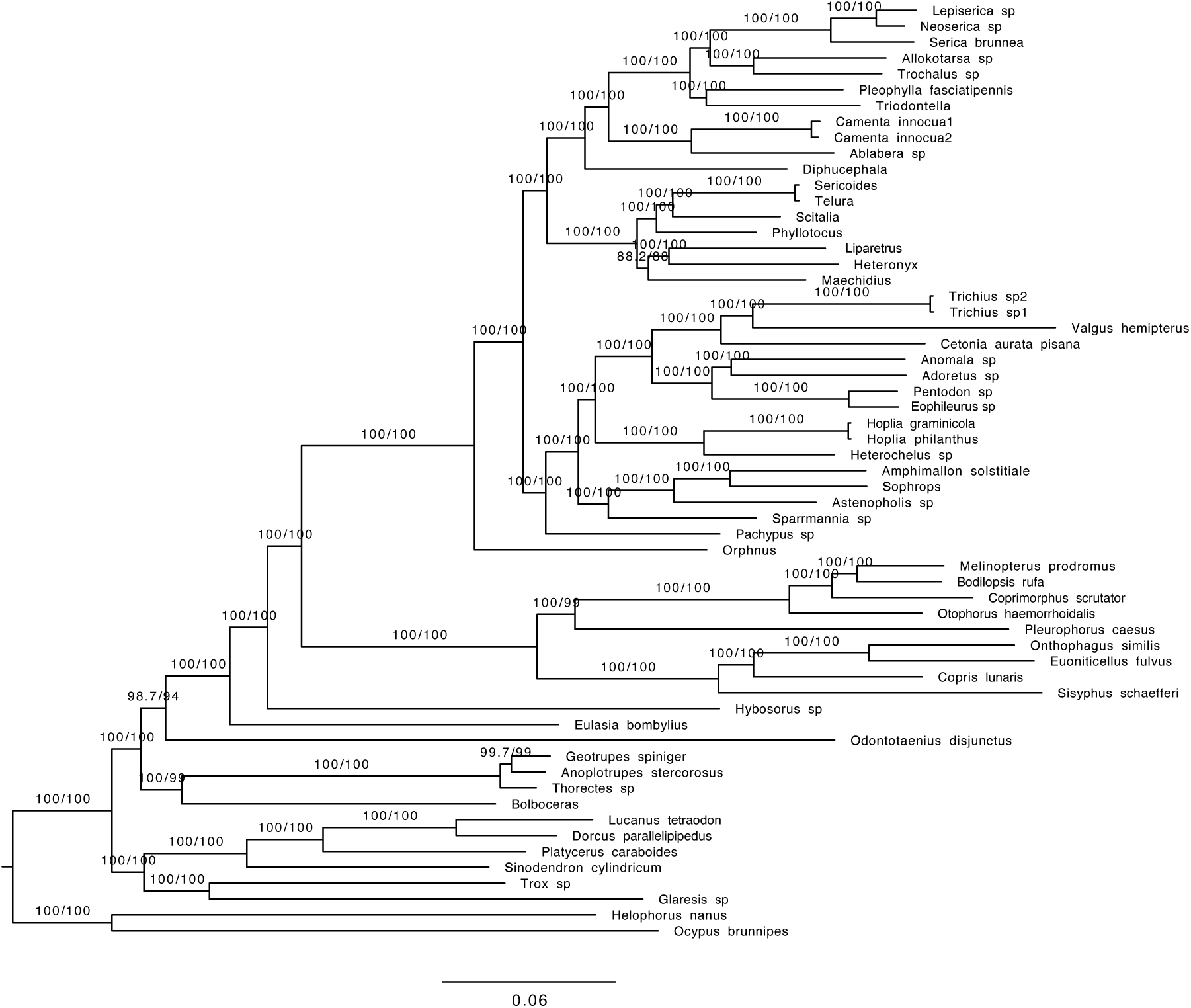
Phylogenetic tree obtained with the concatenated data and amino acid sequences (AA) (Mafft alignment; dataset with 70% complete data; Coleoptera orthologs).

**Figure S20.**
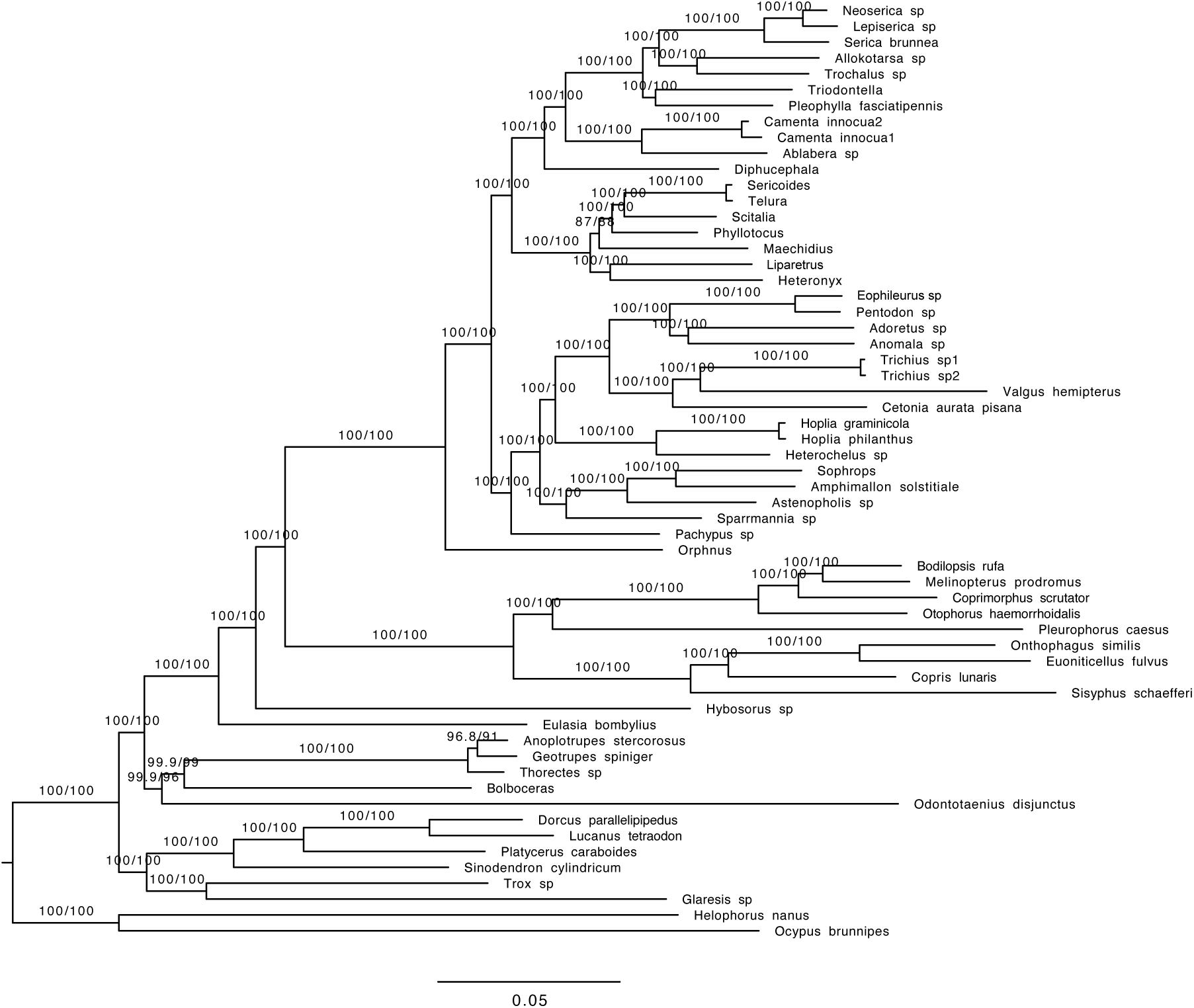
Phylogenetic tree obtained with the concatenated data and nucleotide data containing only the first and second base pair (nt12) (Mafft alignment; dataset with 70% complete data; Coleoptera orthologs).

**Figure S21.**
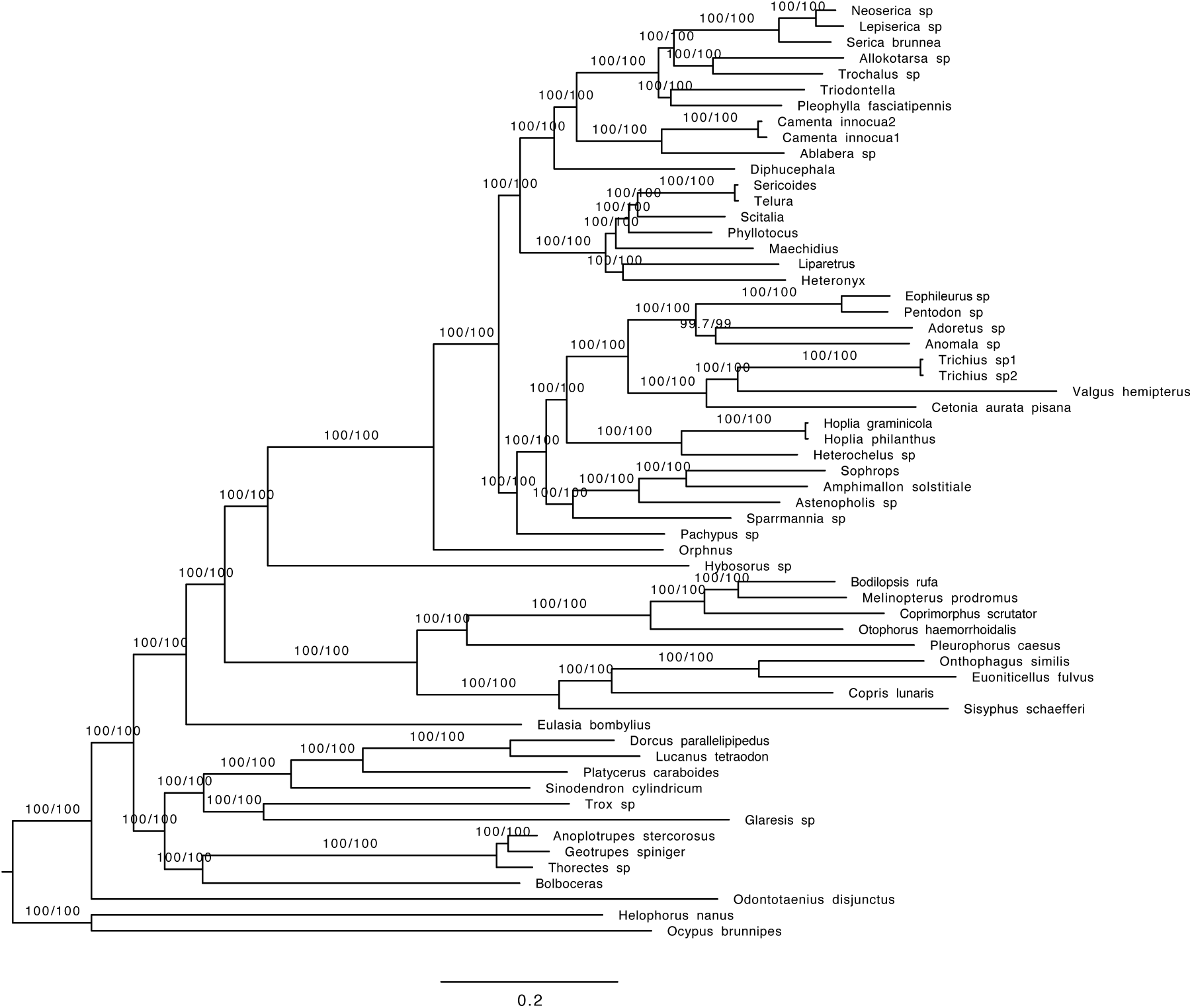
Phylogenetic tree obtained with the concatenated data and nucleotide data containing all three base pairs (nt123) (Mafft alignment; dataset with 70% complete data; Coleoptera orthologs).

**Figure S22.**
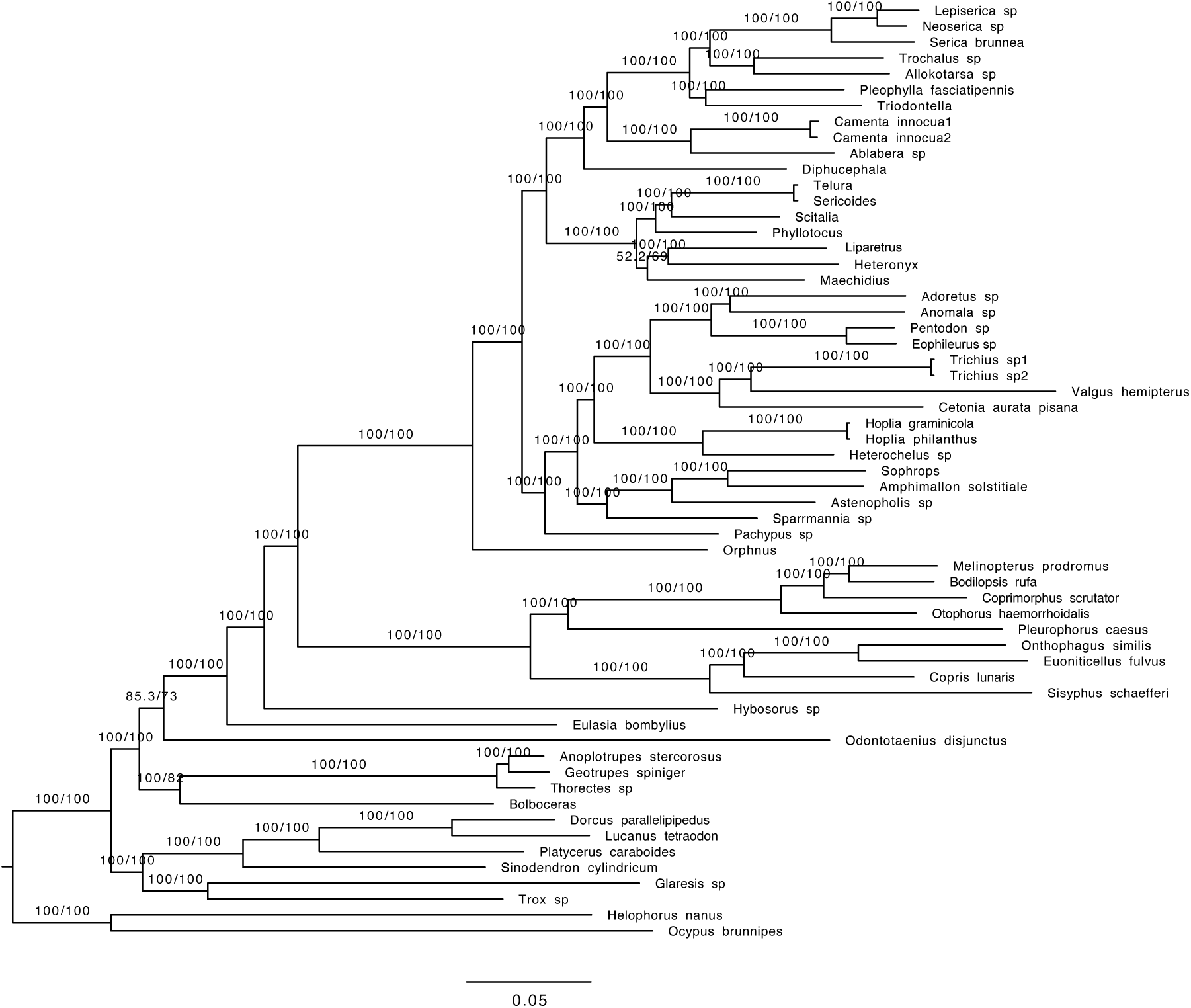
Phylogenetic tree obtained with the concatenated data and amino acid sequences (AA) (HmmAlign alignment; dataset with 70% complete data; Coleoptera orthologs).

**Figure S23.**
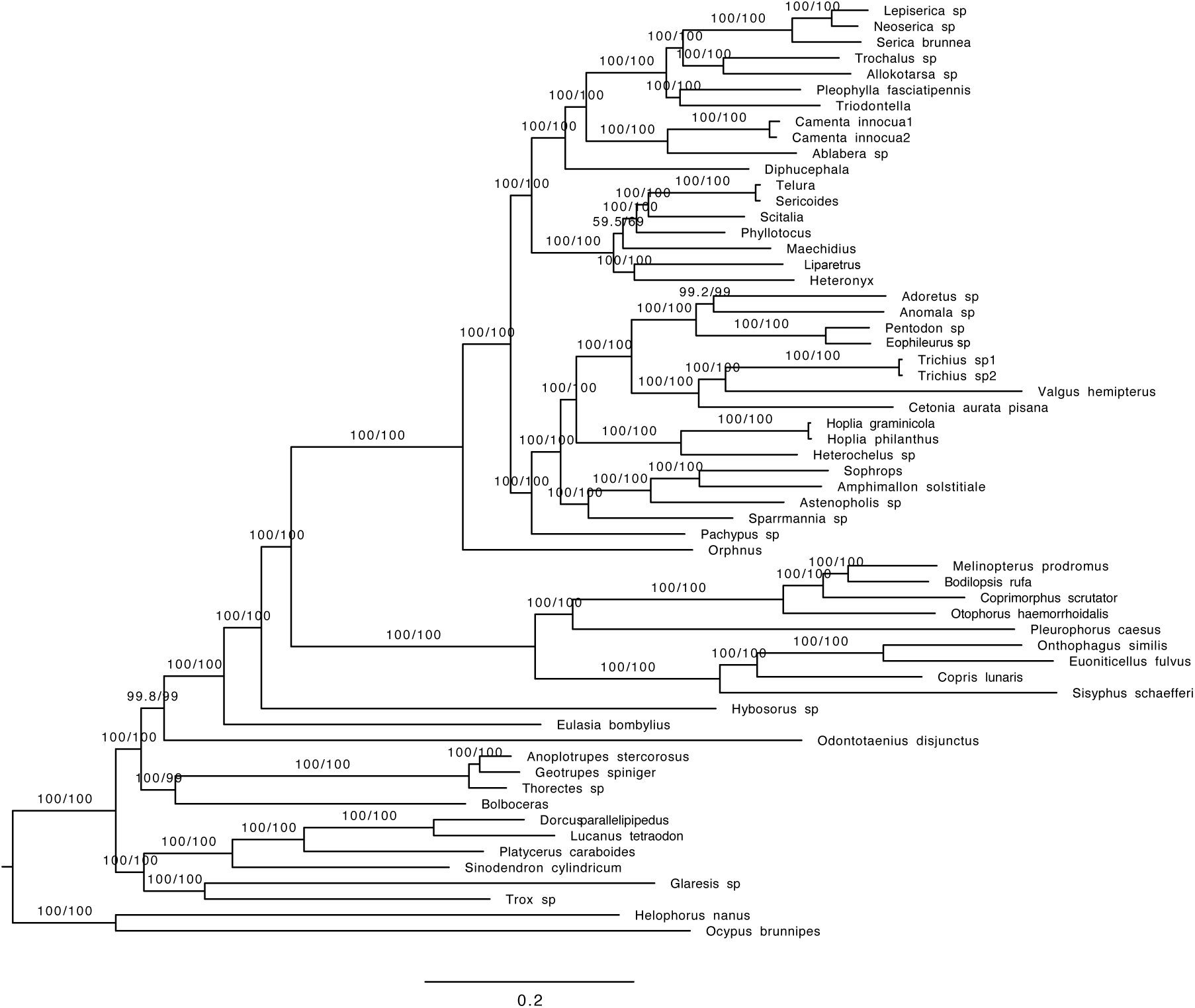
Phylogenetic tree obtained with the concatenated data and nucleotide data containing only the first and second base pair (nt12) (HmmAlign alignment; dataset with 70% complete data; Coleoptera orthologs).

**Figure S24.**
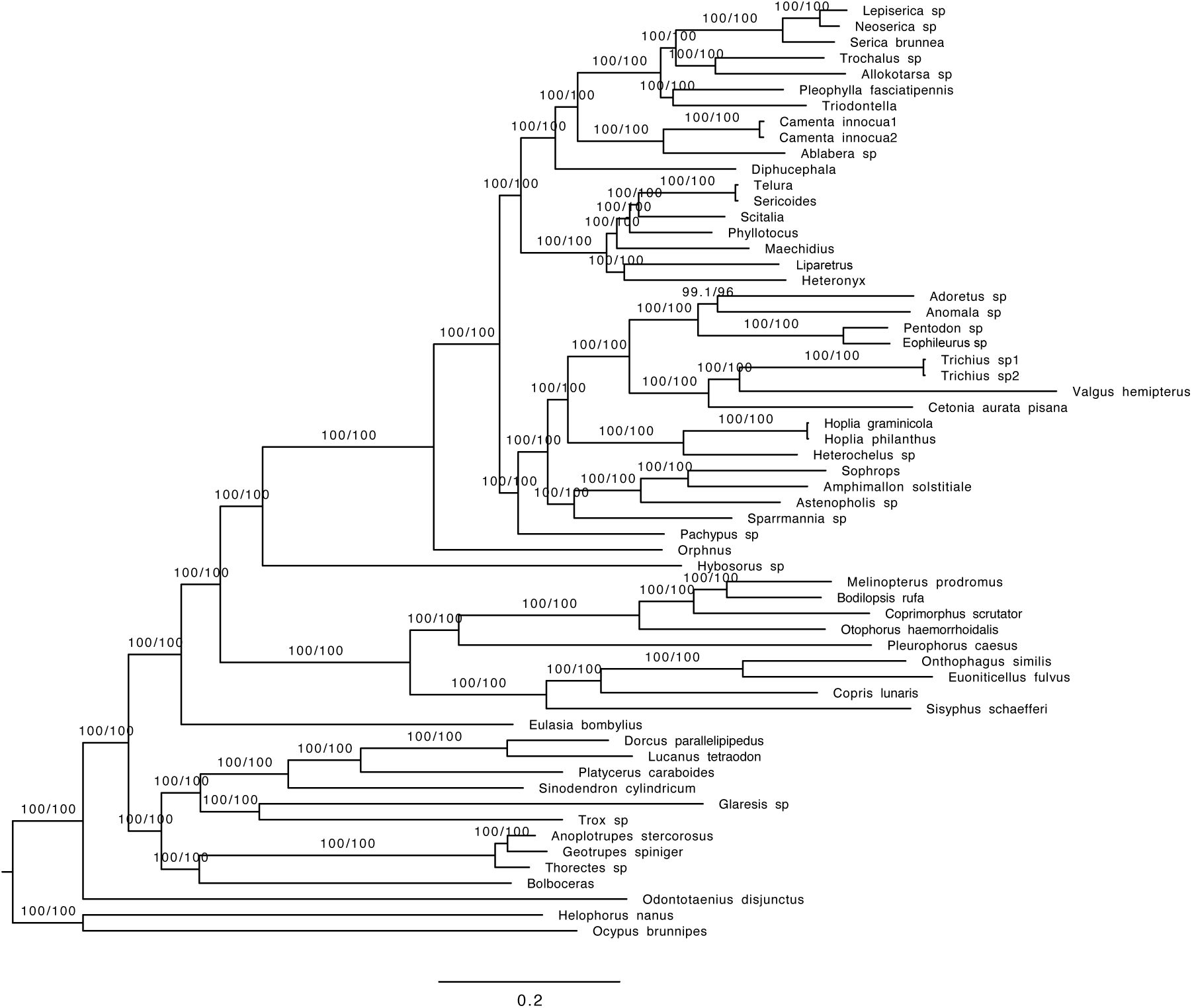
Phylogenetic tree obtained with the concatenated data and nucleotide data containing all three base pairs (nt123) (HmmAlign alignment; dataset with 70% complete data; Coleoptera orthologs).

**Figure S25.**
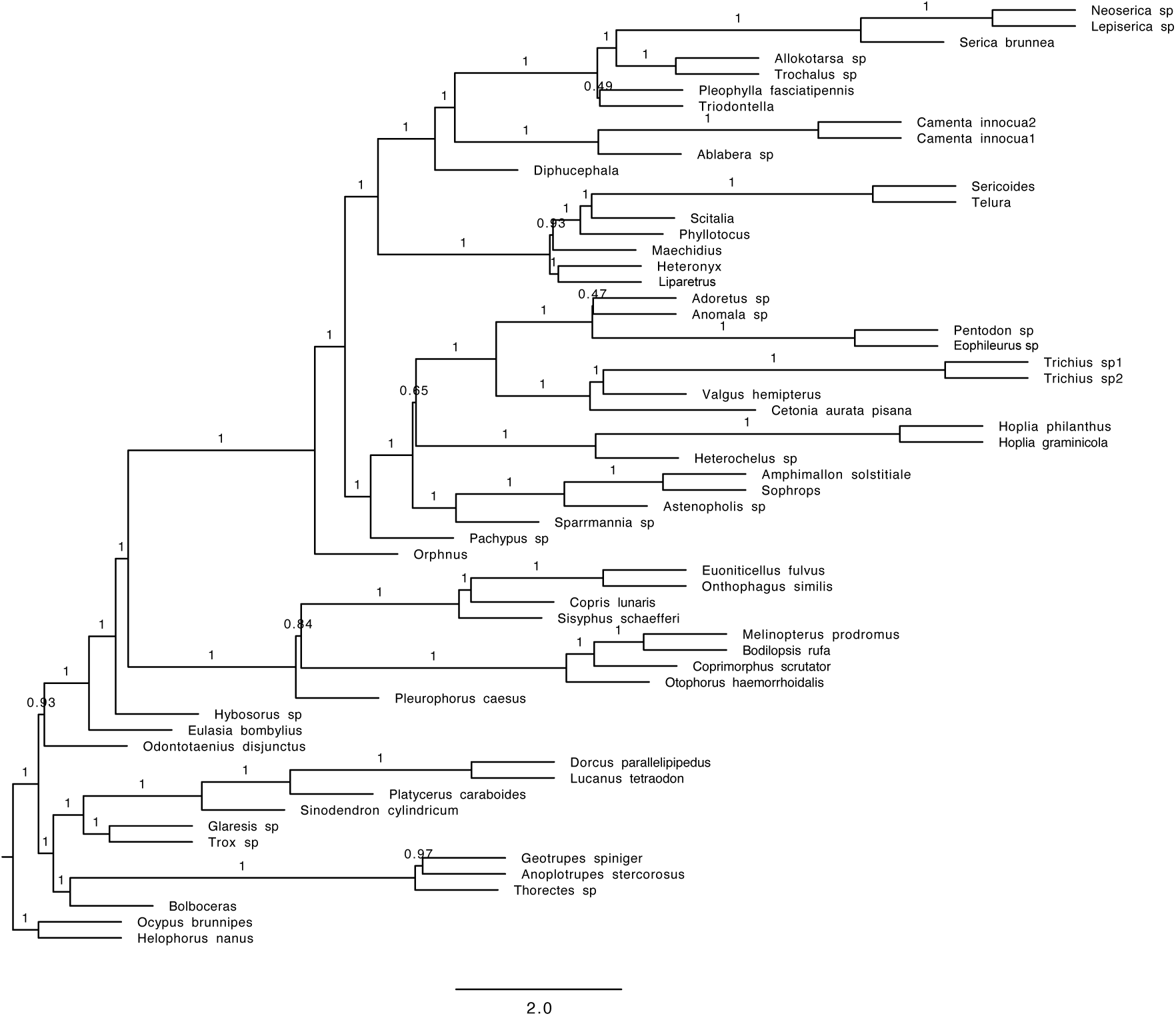
Phylogenetic tree obtained with the coalescent tree search with Astral and amino acid sequences (Mafft alignment; fast evolving genes only; Coleoptera orthologs).

**Figure S26.**
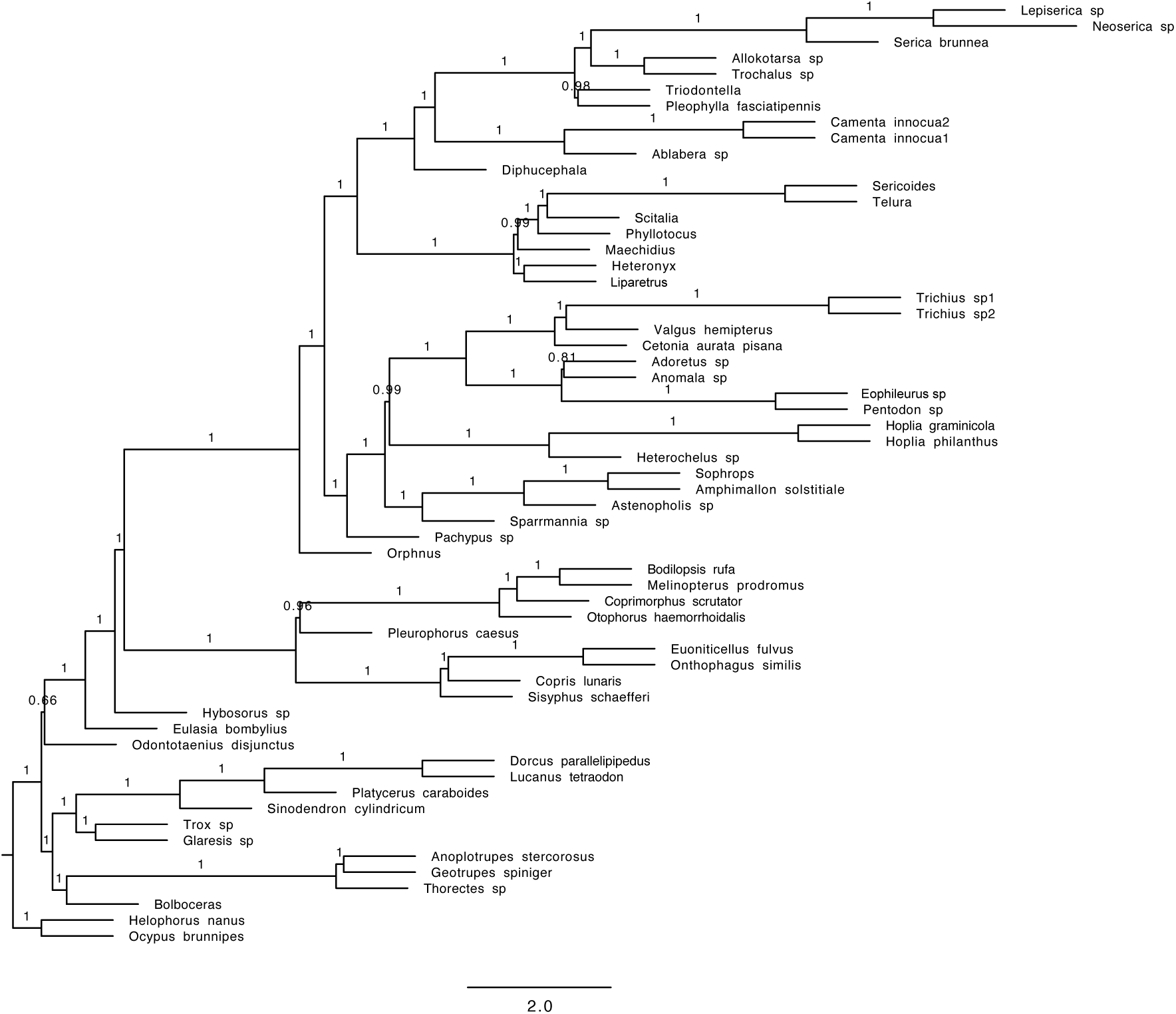
Phylogenetic tree obtained with the coalescent tree search with Astral and nucleotide data containing only the first and second base pair (nt12) (Mafft alignment; fast evolving genes only; Coleoptera orthologs).

**Figure S27.**
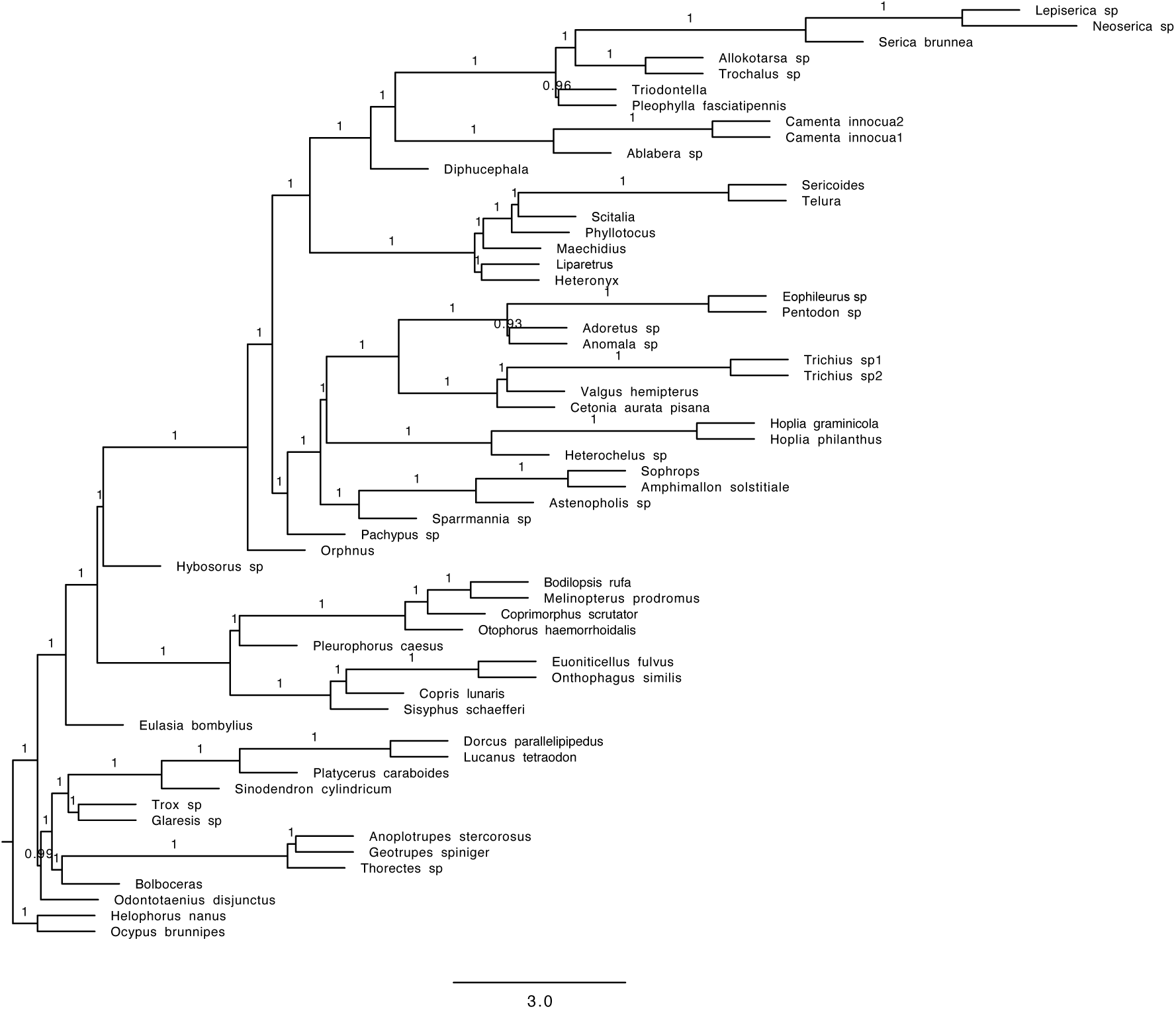
Phylogenetic tree obtained with the coalescent tree search with Astral and nucleotide data containing all three base pairs (nt123) (Mafft alignment; fast evolving genes only; Coleoptera orthologs).

**Figure S28.**
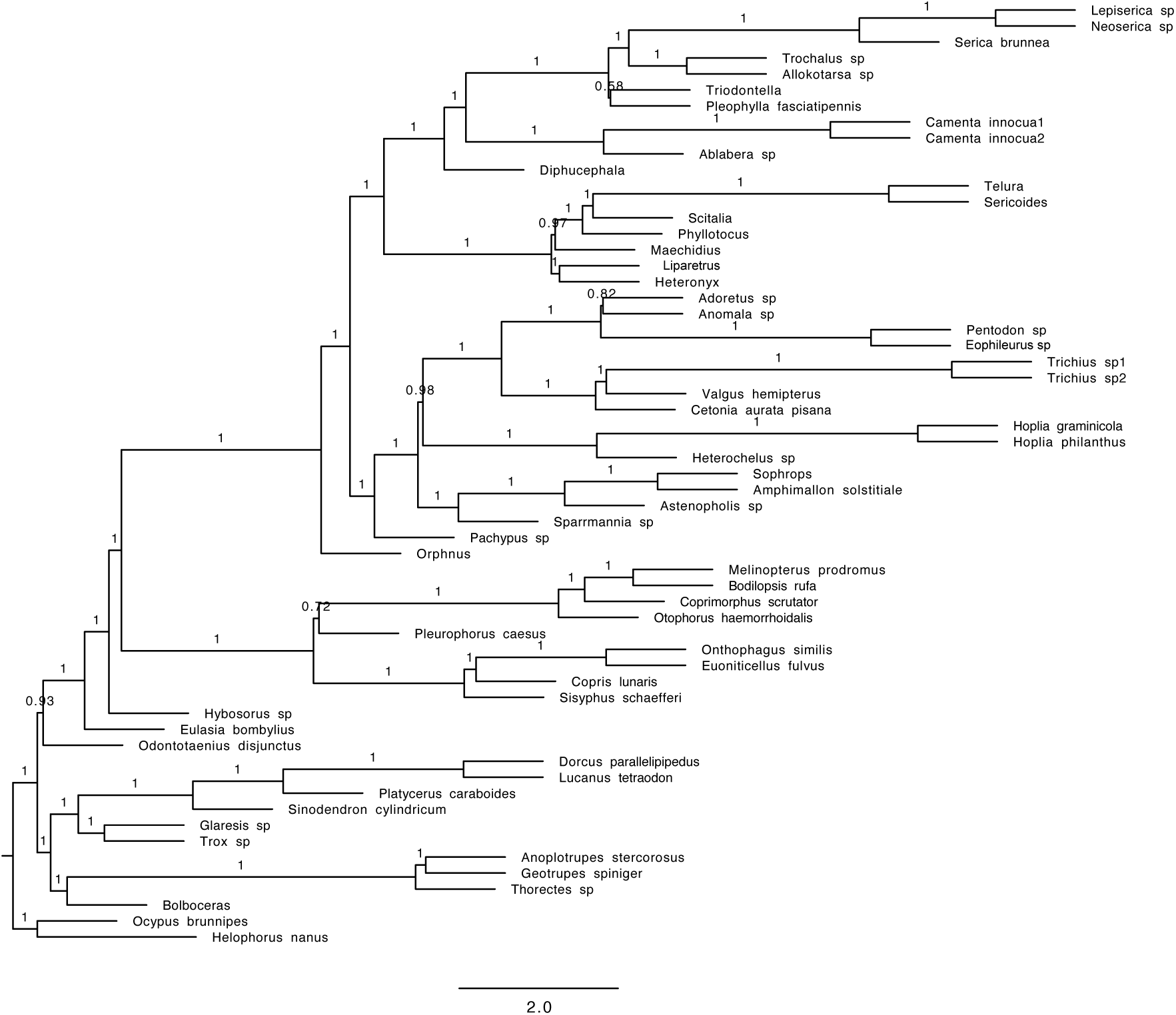
Phylogenetic tree obtained with the coalescent tree search with Astral and amino acid sequences (HmmAlign alignment; fast evolving genes only; Coleoptera orthologs).

**Figure S29.**
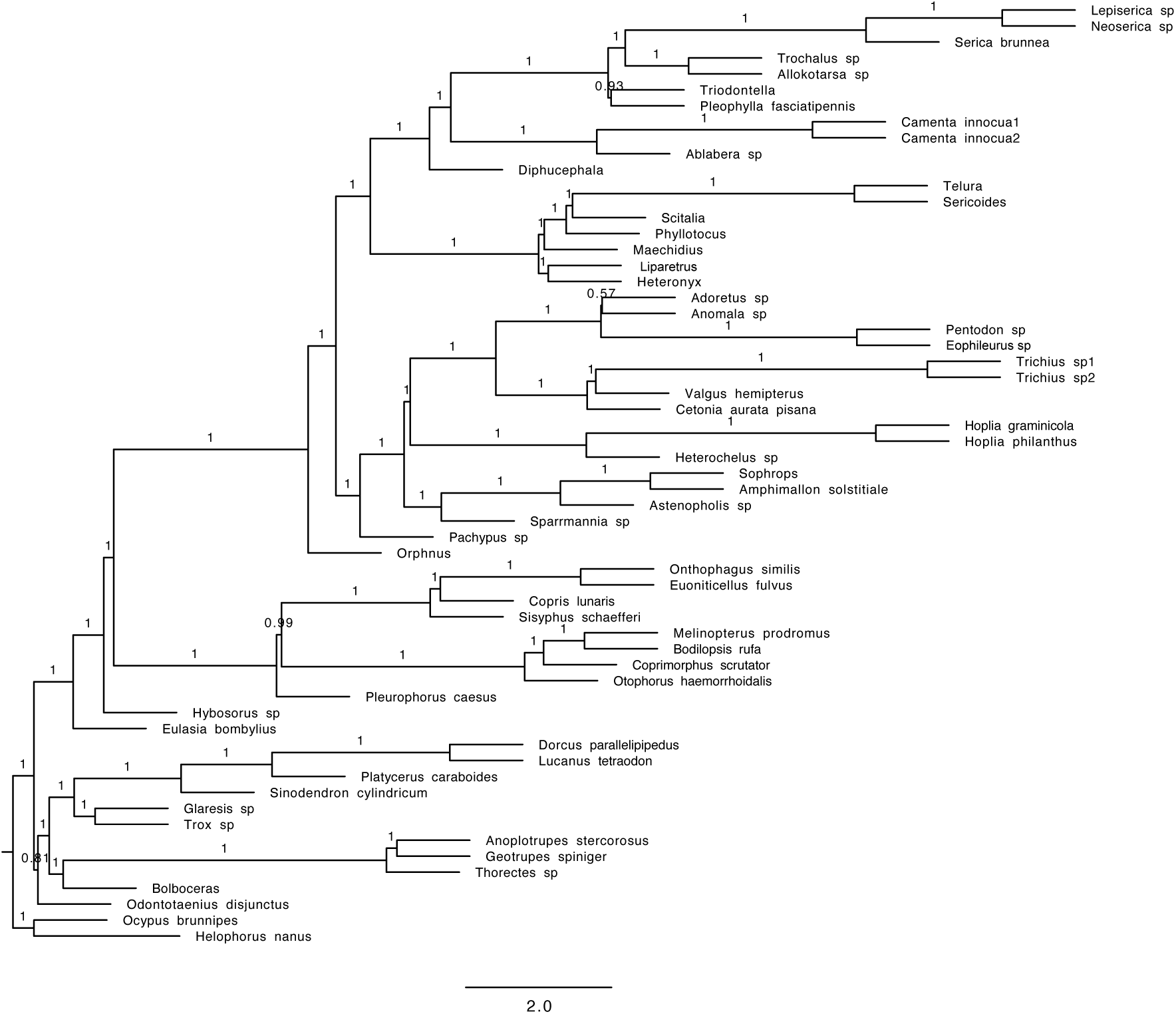
Phylogenetic tree obtained with the coalescent tree search with Astral and nucleotide data containing only the first and second base pair (nt12) (HmmAlign alignment; fast evolving genes only; Coleoptera orthologs).

**Figure S30.**
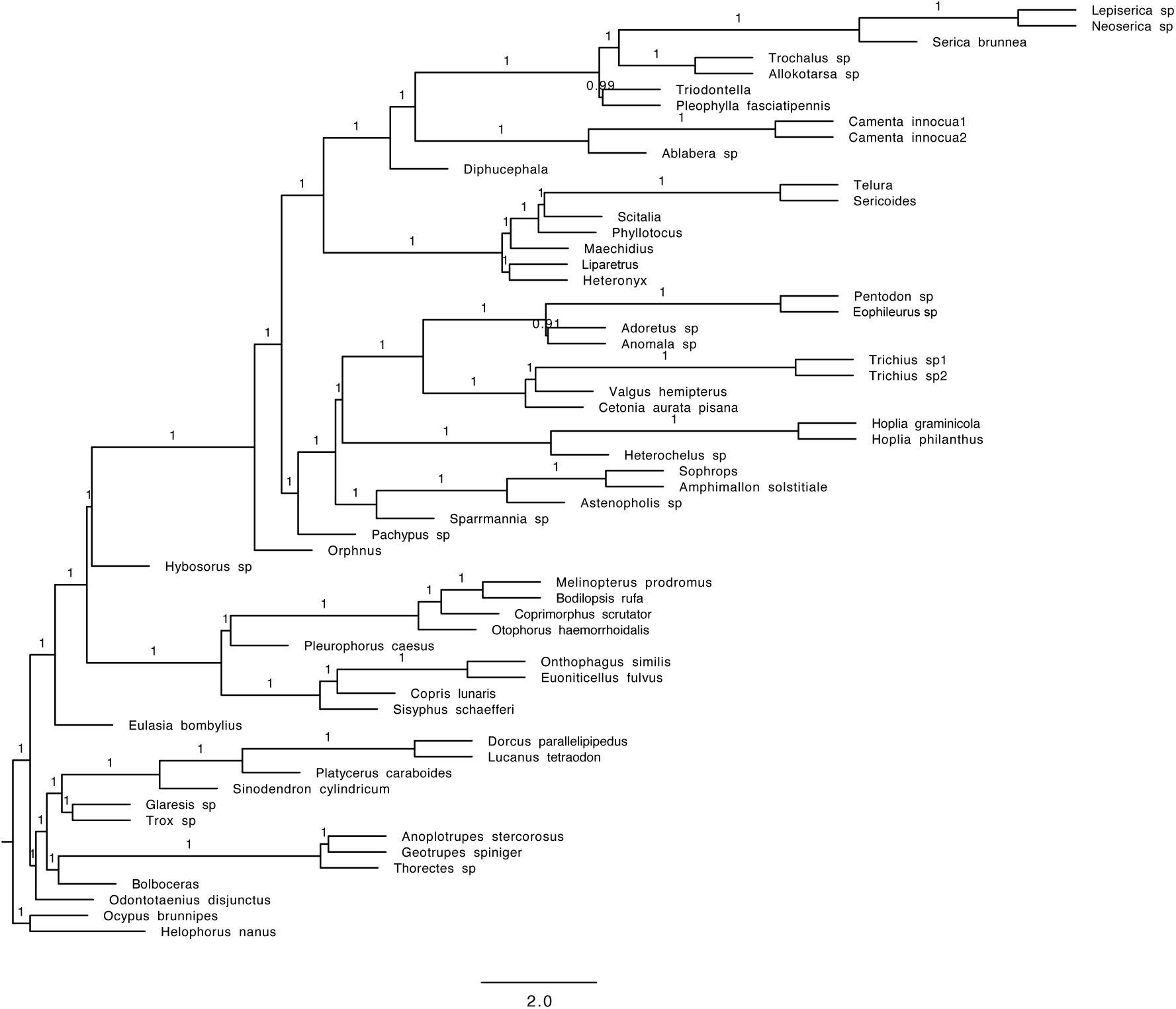
Phylogenetic tree obtained with the coalescent tree search with Astral and nucleotide data containing all three base pairs (nt123) (HmmAlign alignment; fast evolving genes only; Coleoptera orthologs).

**Figure S31.**
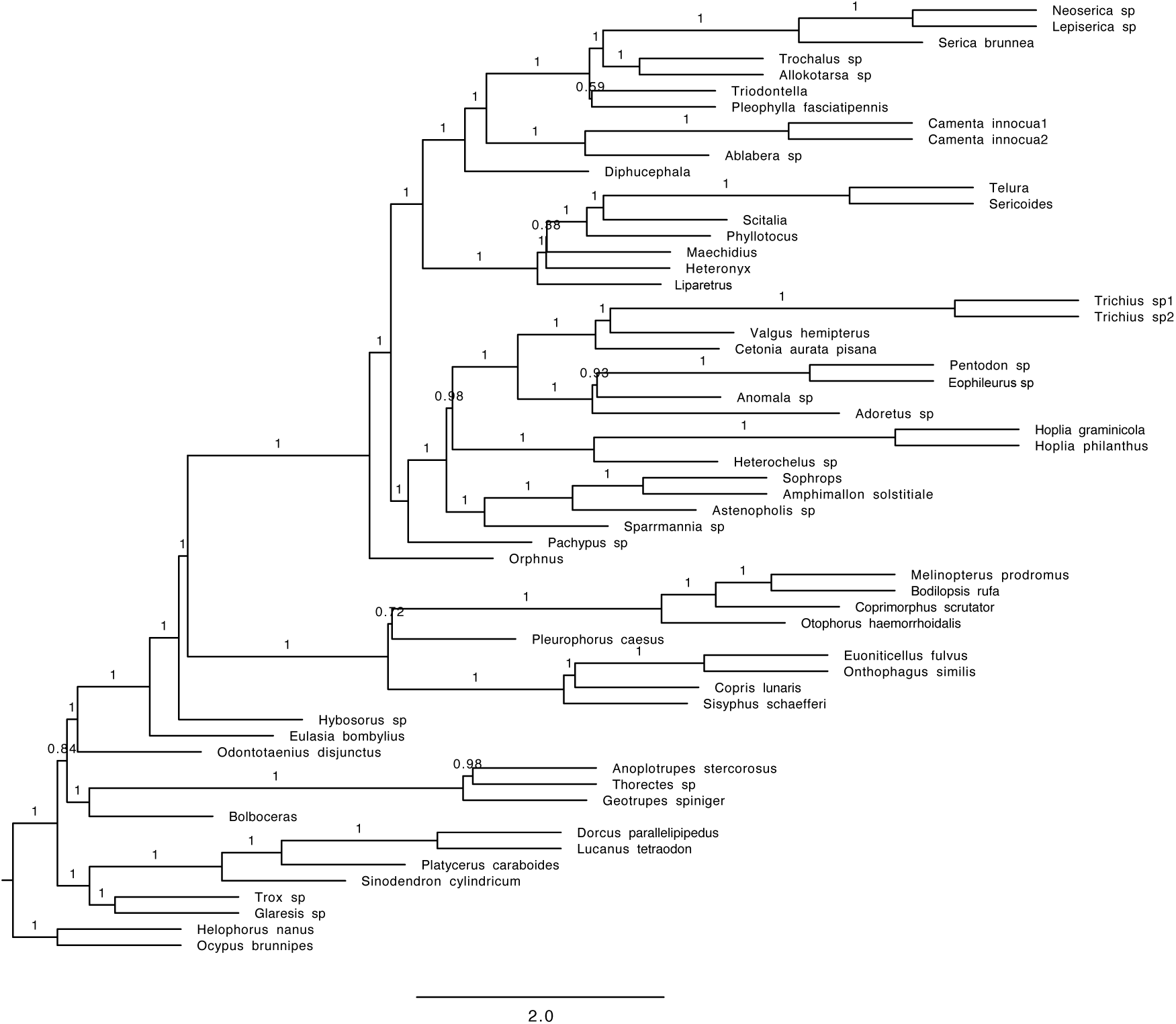
Phylogenetic tree obtained with the coalescent tree search with Astral and amino acid sequences (Mafft alignment; slowly evolving genes only; Coleoptera orthologs).

**Figure S32.**
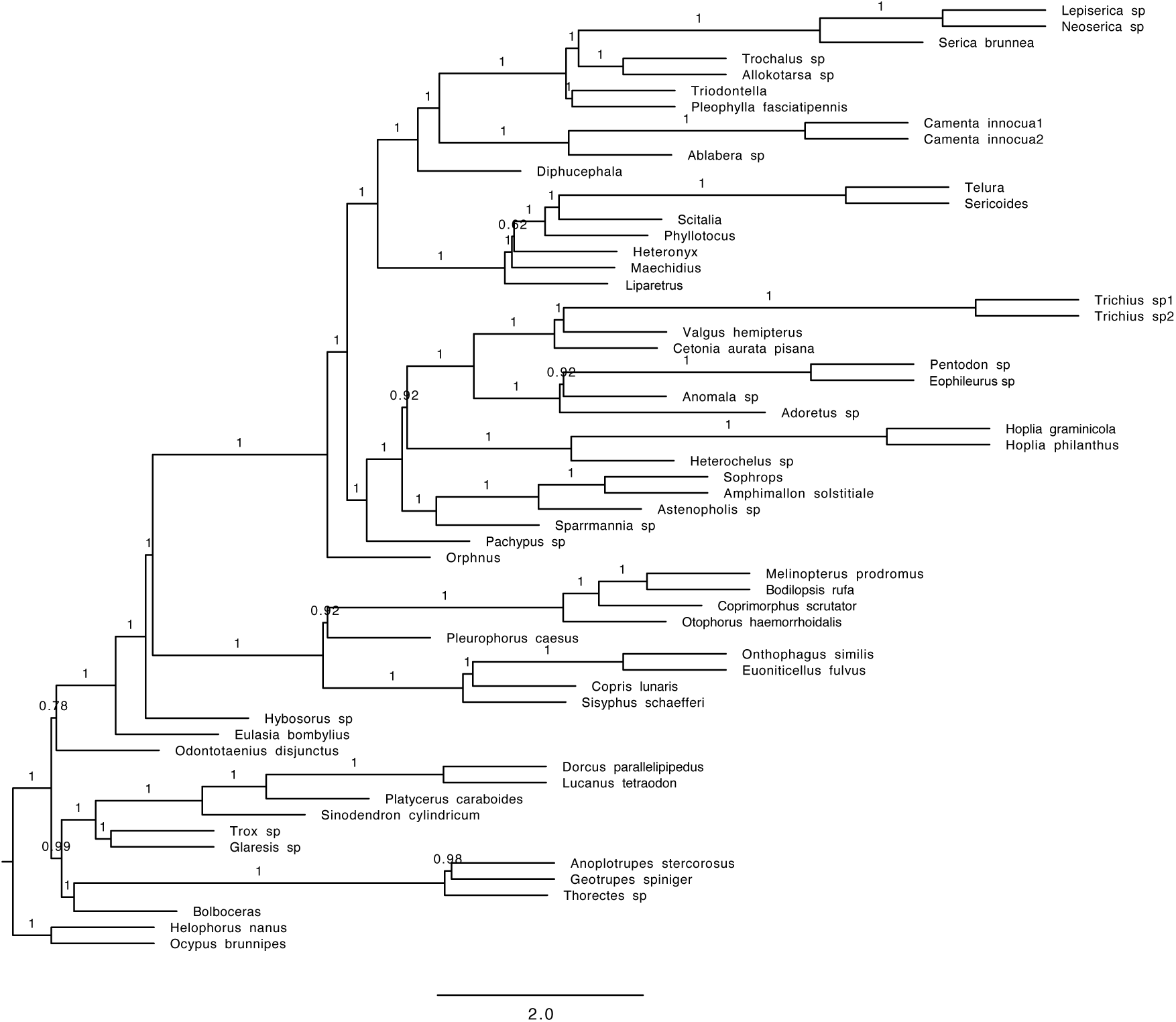
Phylogenetic tree obtained with the coalescent tree search with Astral and nucleotide data containing only the first and second base pair (nt12) (Mafft alignment; slowly evolving genes only; Coleoptera orthologs).

**Figure S33.**
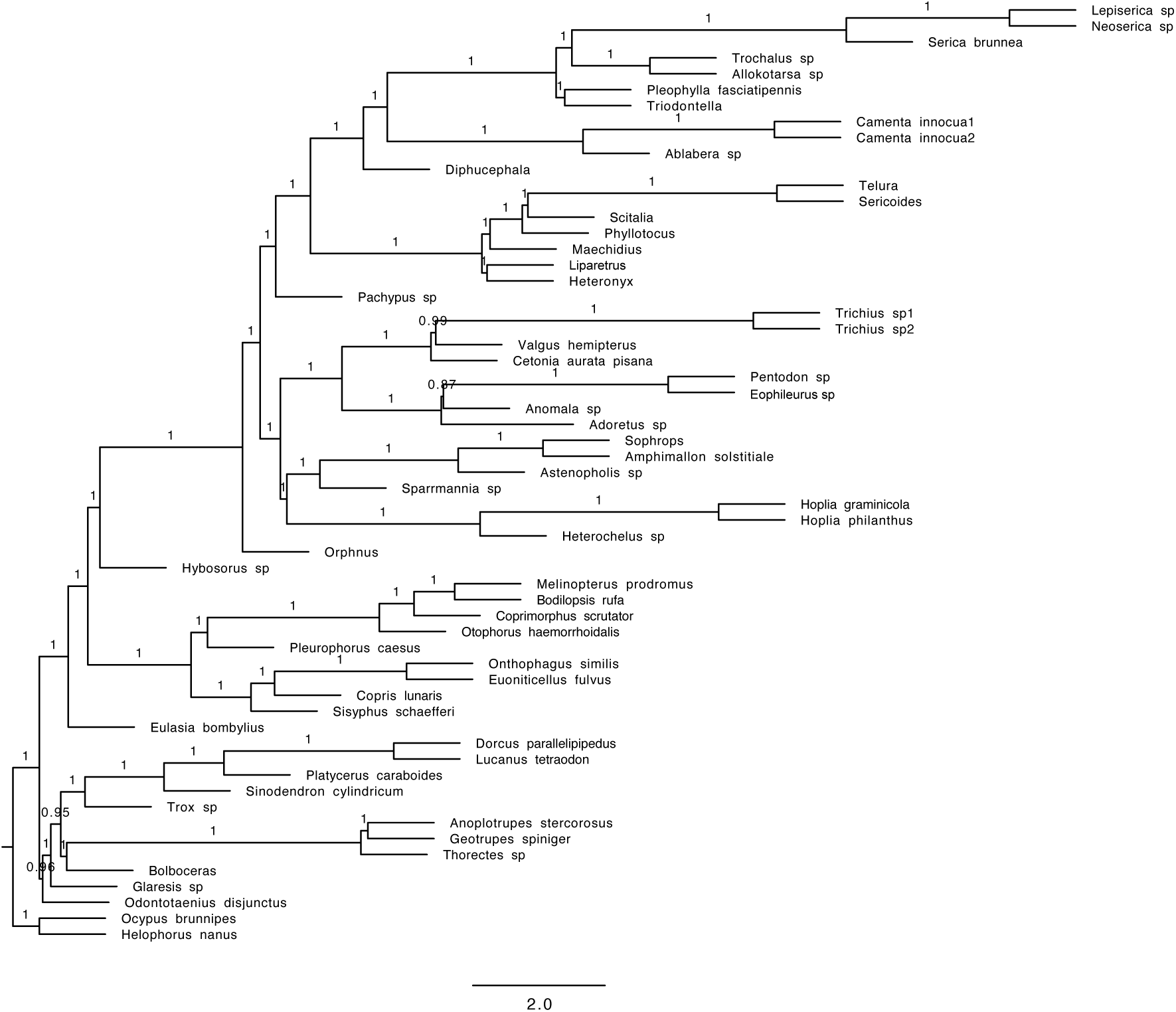
Phylogenetic tree obtained with the coalescent tree search with Astral and nucleotide data containing all three base pairs (nt123) (Mafft alignment; slowly evolving genes only; Coleoptera orthologs).

**Figure S34.**
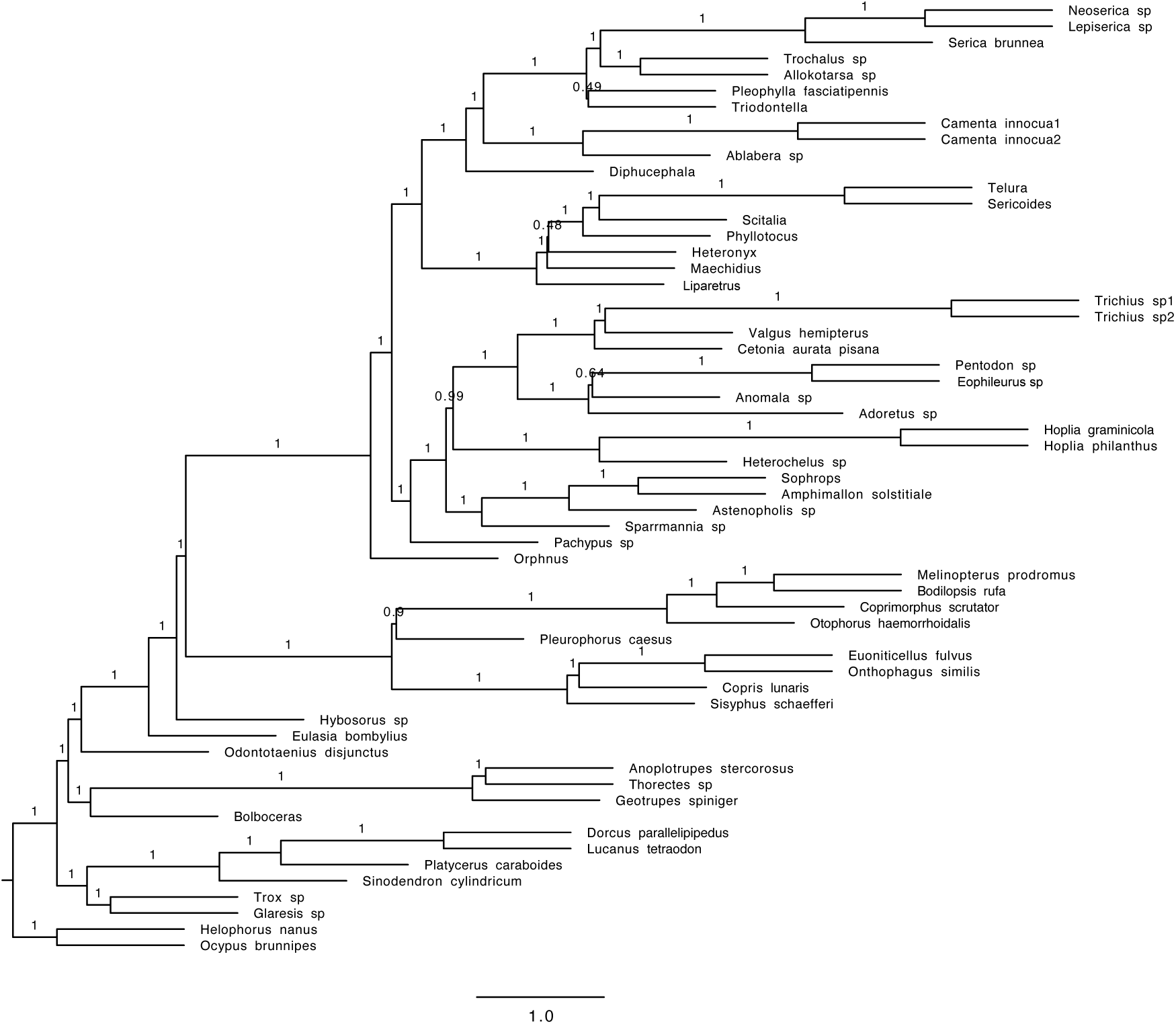
Phylogenetic tree obtained with the coalescent tree search with Astral and amino acid sequences (HmmAlign alignment; slowly evolving genes only; Coleoptera orthologs).

**Figure S35.**
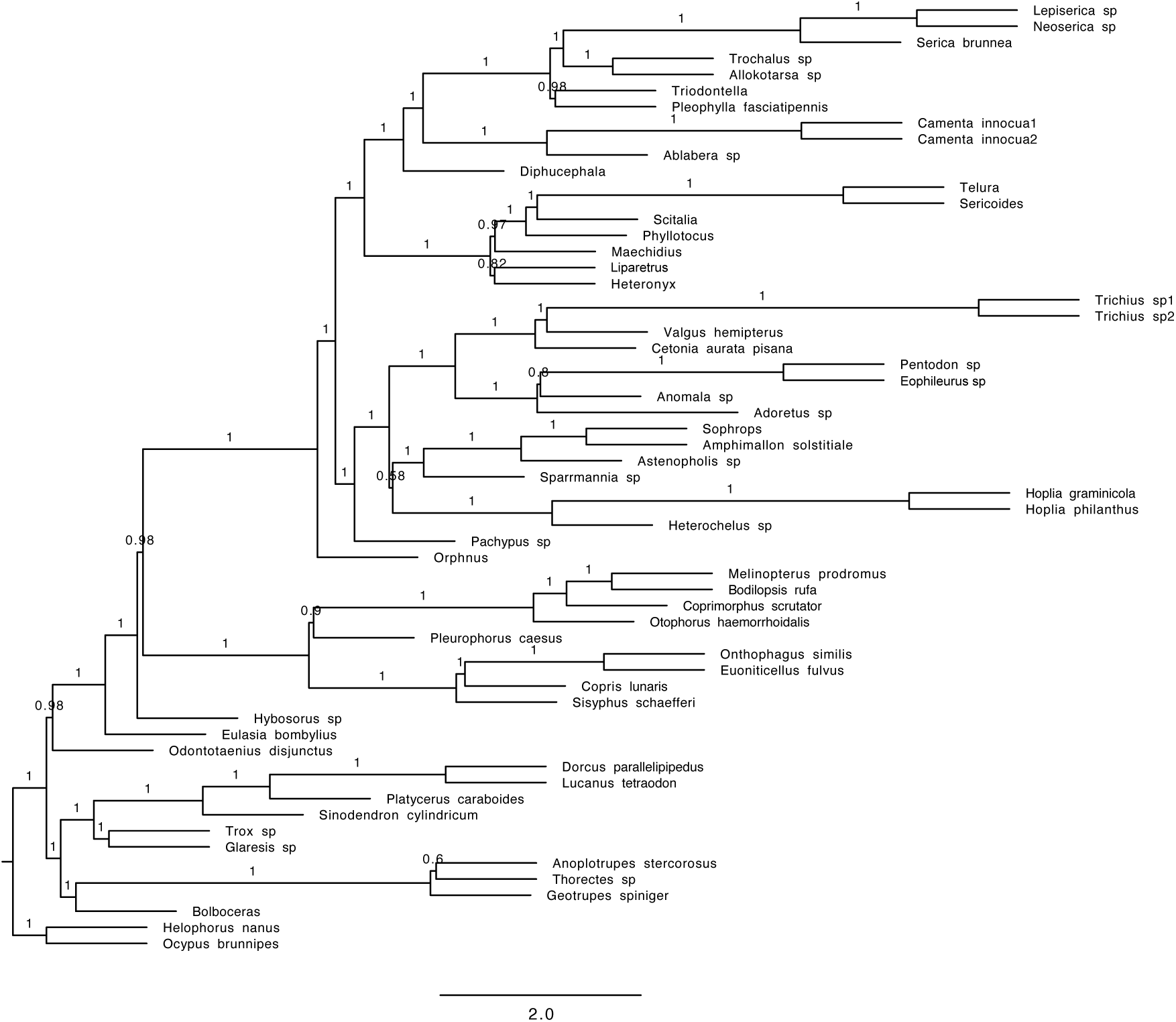
Phylogenetic tree obtained with the coalescent tree search with Astral and nucleotide data containing only the first and second base pair (nt12) (HmmAlign alignment; slowly evolving genes only; Coleoptera orthologs).

**Figure S36.**
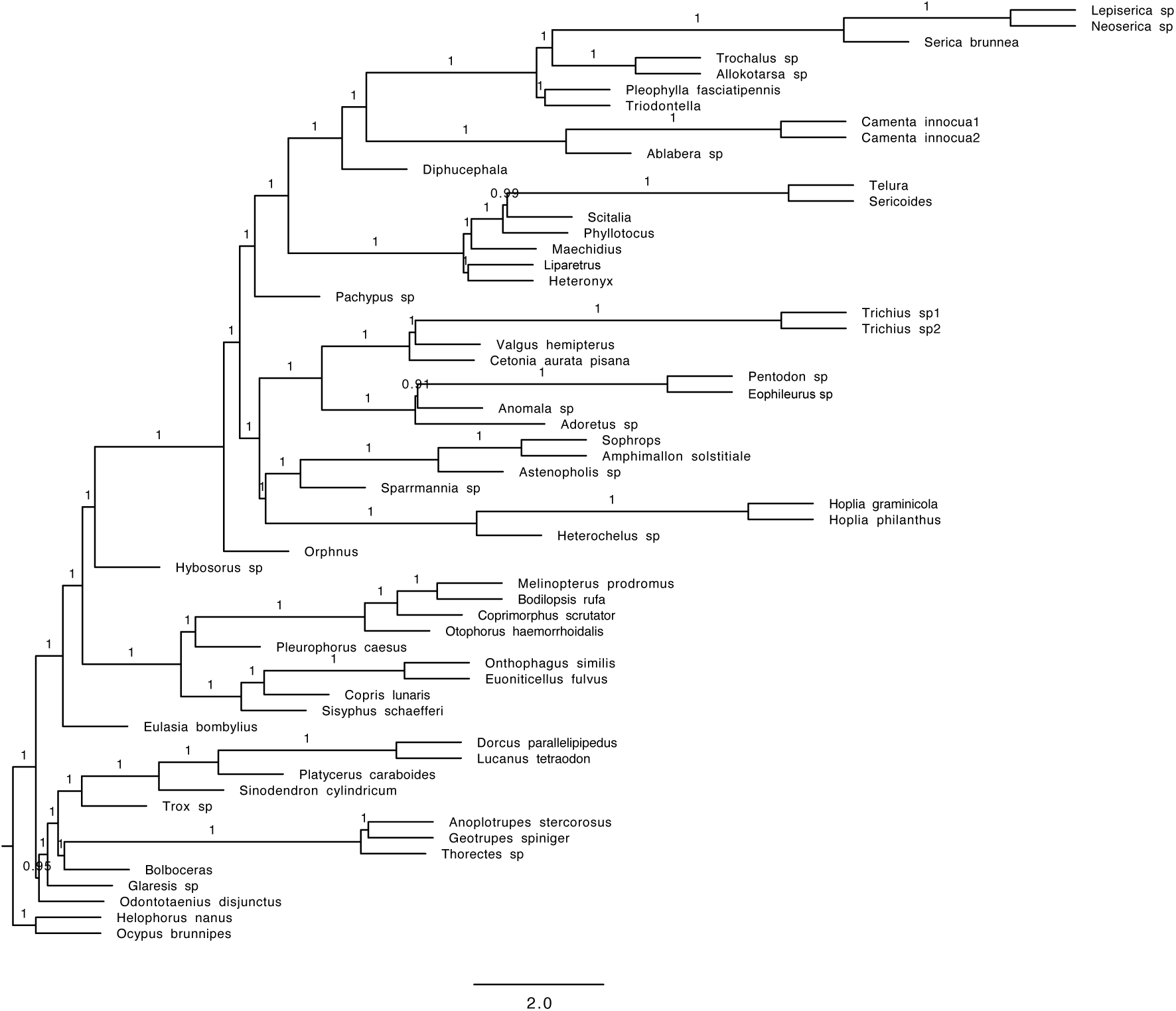
Phylogenetic tree obtained with the coalescent tree search with Astral and nucleotide data containing all three base pairs (nt123) (HmmAlign alignment; slowly evolving genes only; Coleoptera orthologs).

**Figure S37.**
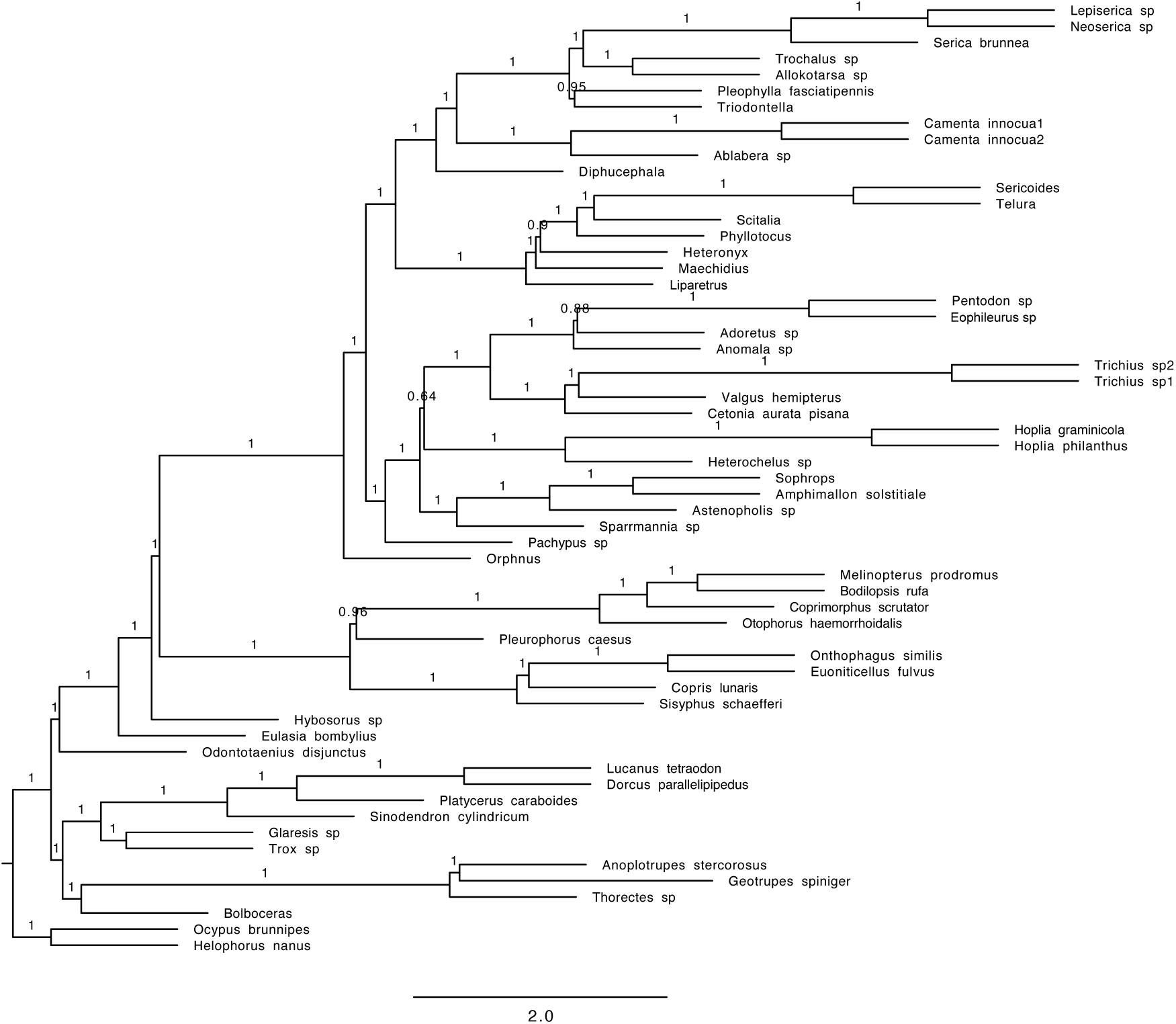
Phylogenetic tree obtained with the coalescent tree search with Astral and amino acid sequences (Mafft alignment; full dataset; Endopterygota orthologs).

**Figure S38.**
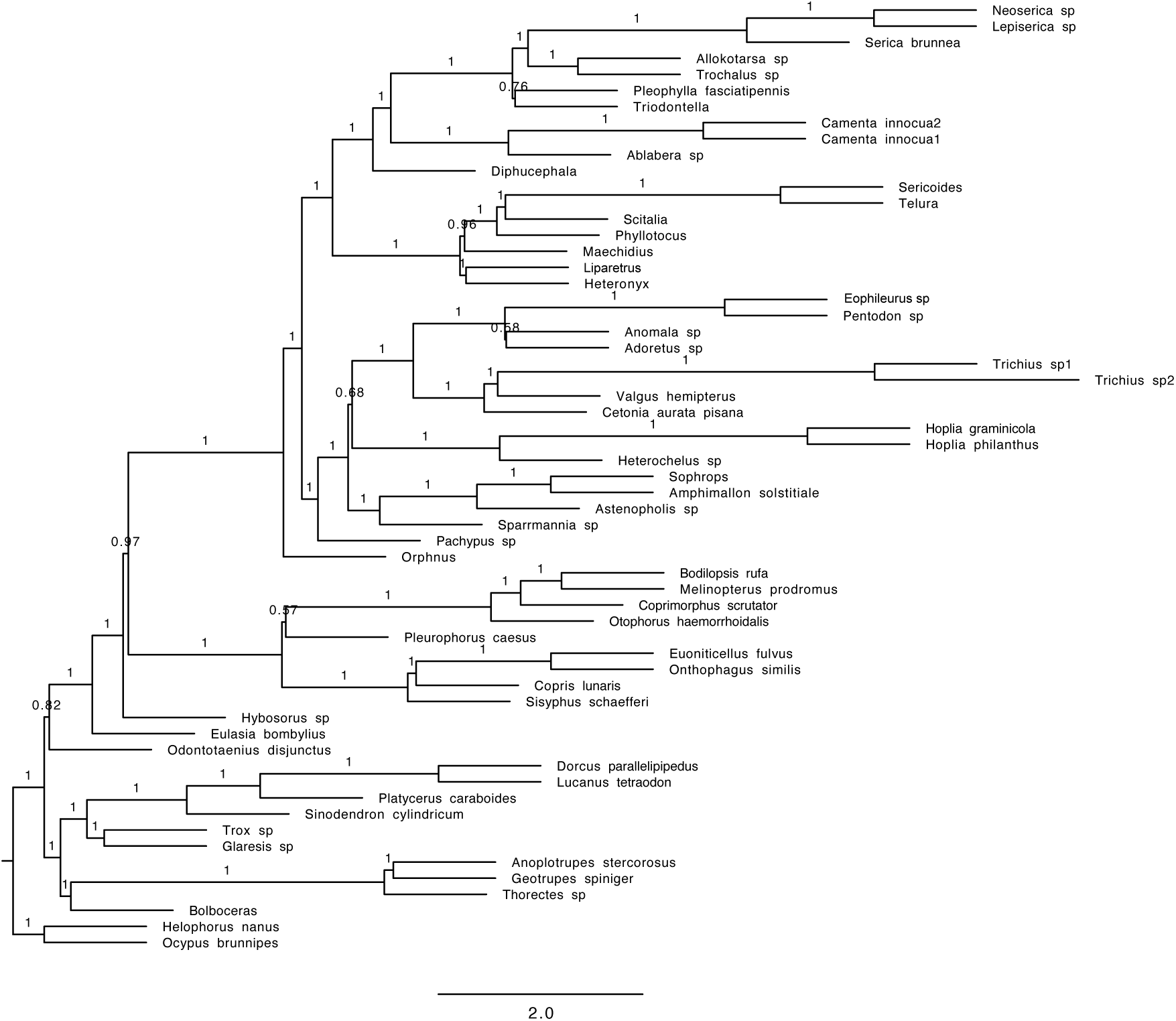
Phylogenetic tree obtained with the coalescent tree search with Astral and nucleotide data containing only the first and second base pair (nt12) (Mafft alignment; full dataset; Endopterygota orthologs).

**Figure S39.**
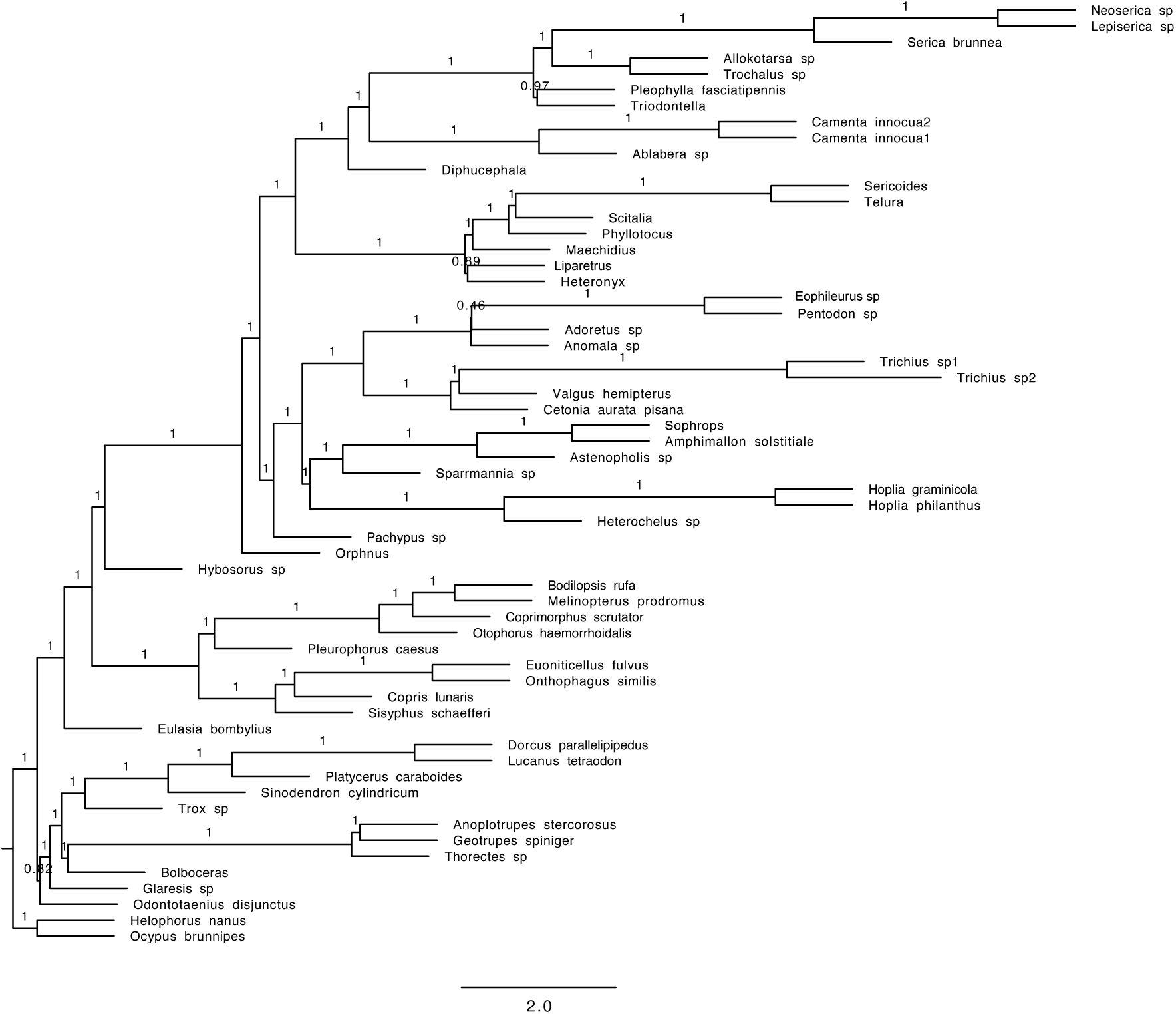
Phylogenetic tree obtained with the coalescent tree search with Astral and nucleotide data containing all three base pairs (nt123) (Mafft alignment; full dataset; Endopterygota orthologs).

**Figure S40.**
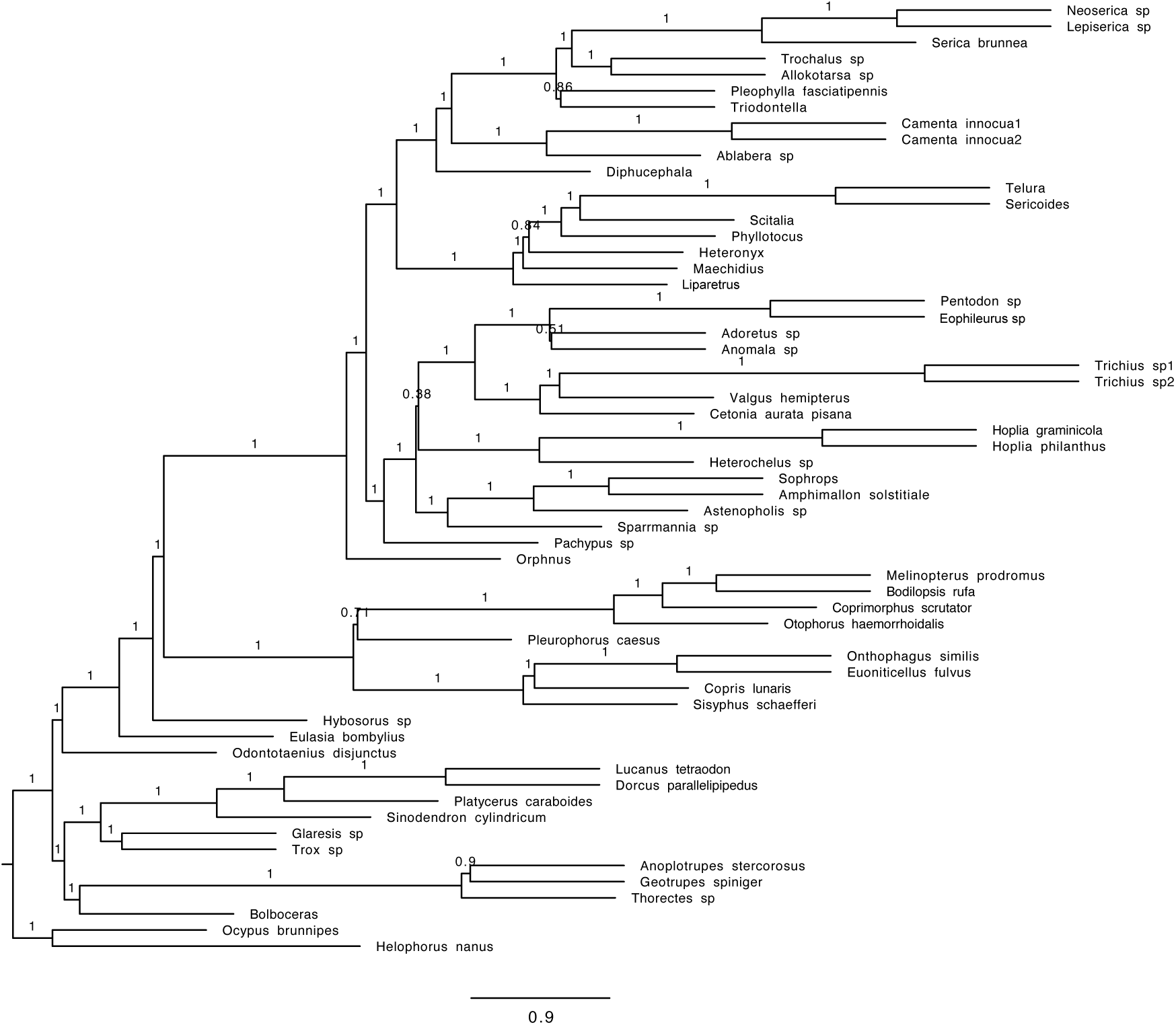
Phylogenetic tree obtained with the coalescent tree search with Astral and amino acid sequences (HmmAlign alignment; full dataset; Endopterygota orthologs).

**Figure S41.**
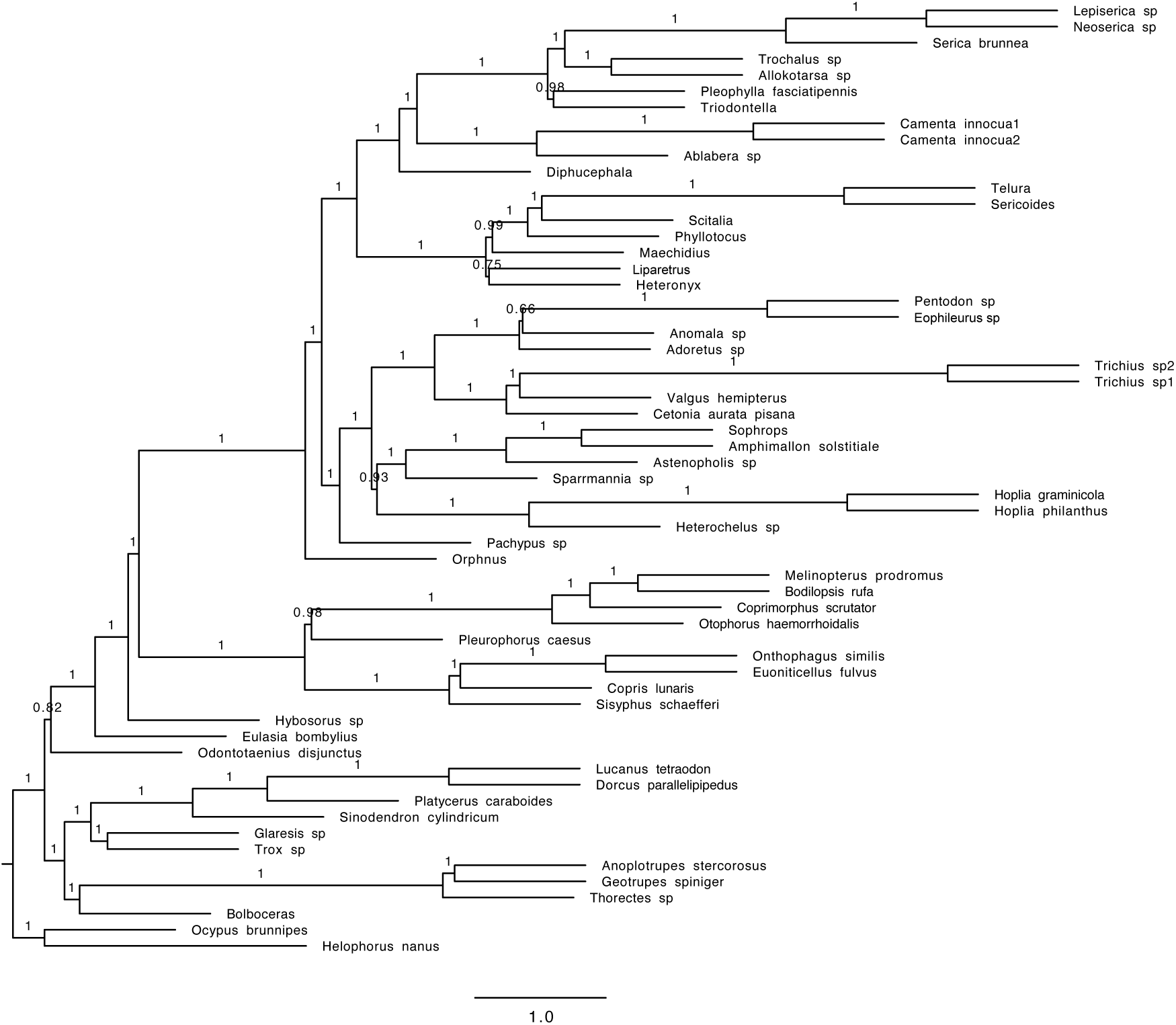
Phylogenetic tree obtained with the coalescent tree search with Astral and nucleotide data containing only the first and second base pair (nt12) (HmmAlign alignment; full dataset; Endopterygota orthologs).

**Figure S42.**
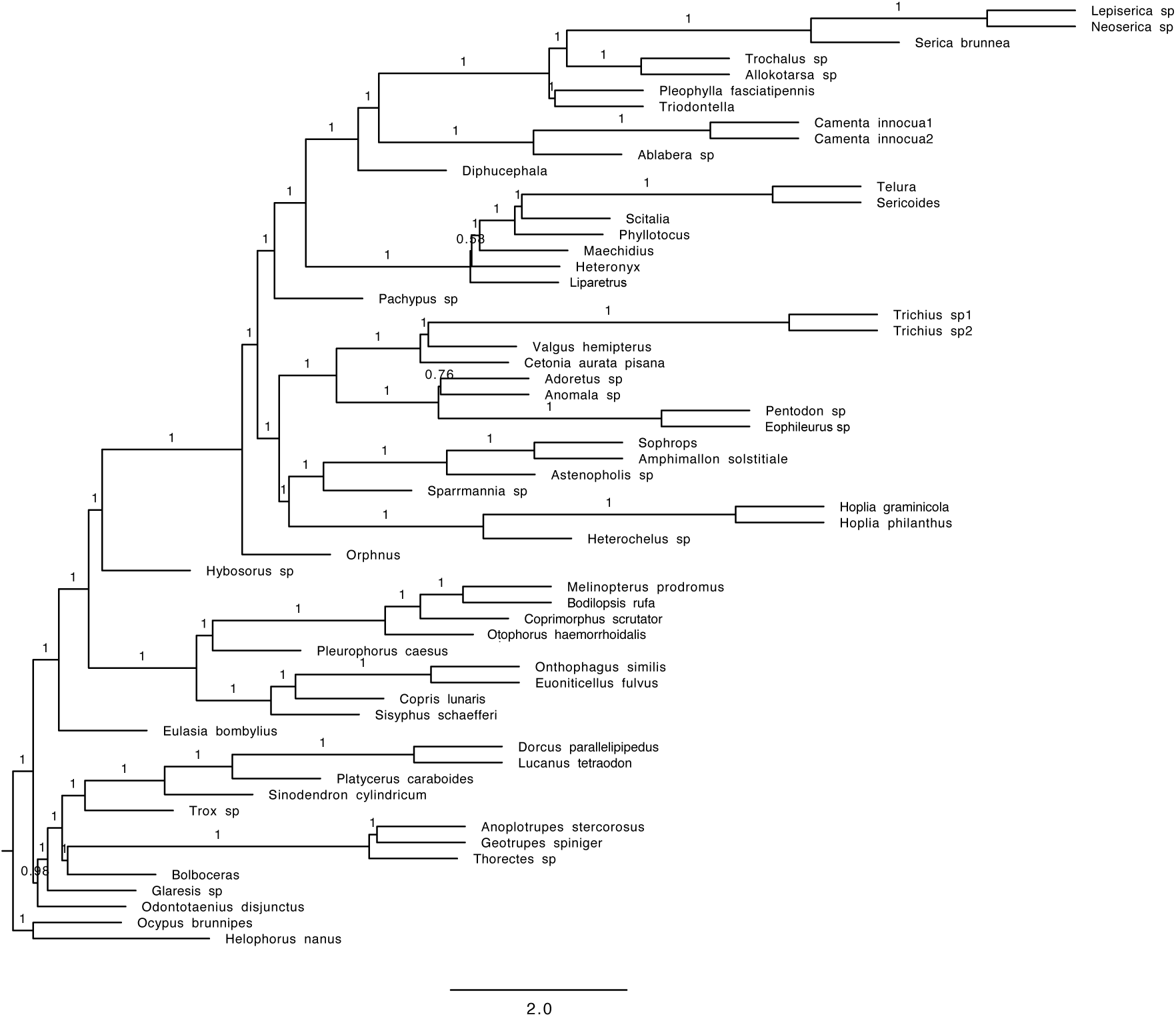
Phylogenetic tree obtained with the coalescent tree search with Astral and nucleotide data containing all three base pairs (nt123) (HmmAlign alignment; full dataset; Endopterygota orthologs).

**Figure S43.**
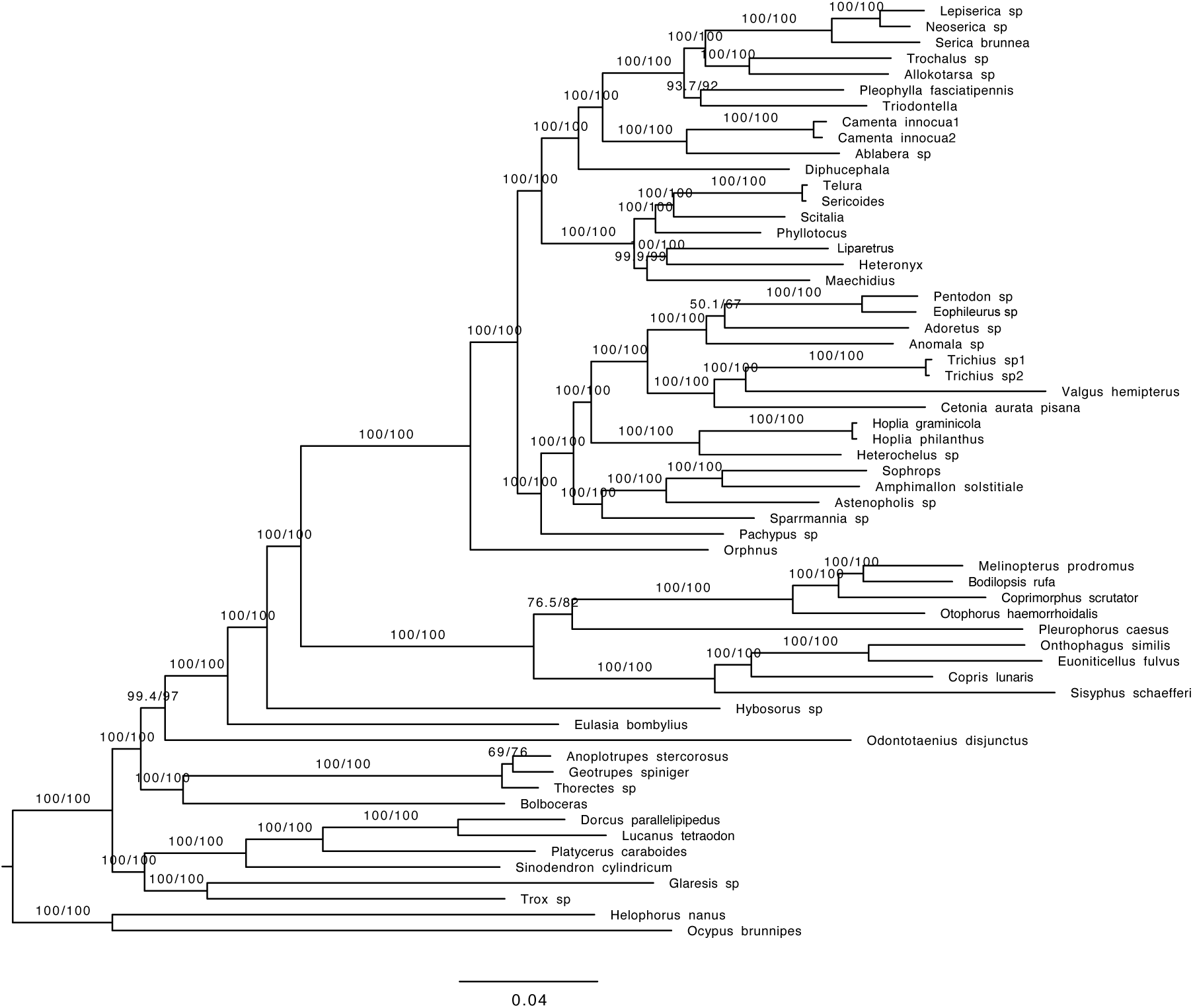
Phylogenetic tree obtained with the concatenated data and amino acid sequences (AA) (Mafft alignment; full dataset; Endopterygota orthologs).

**Figure S44.**
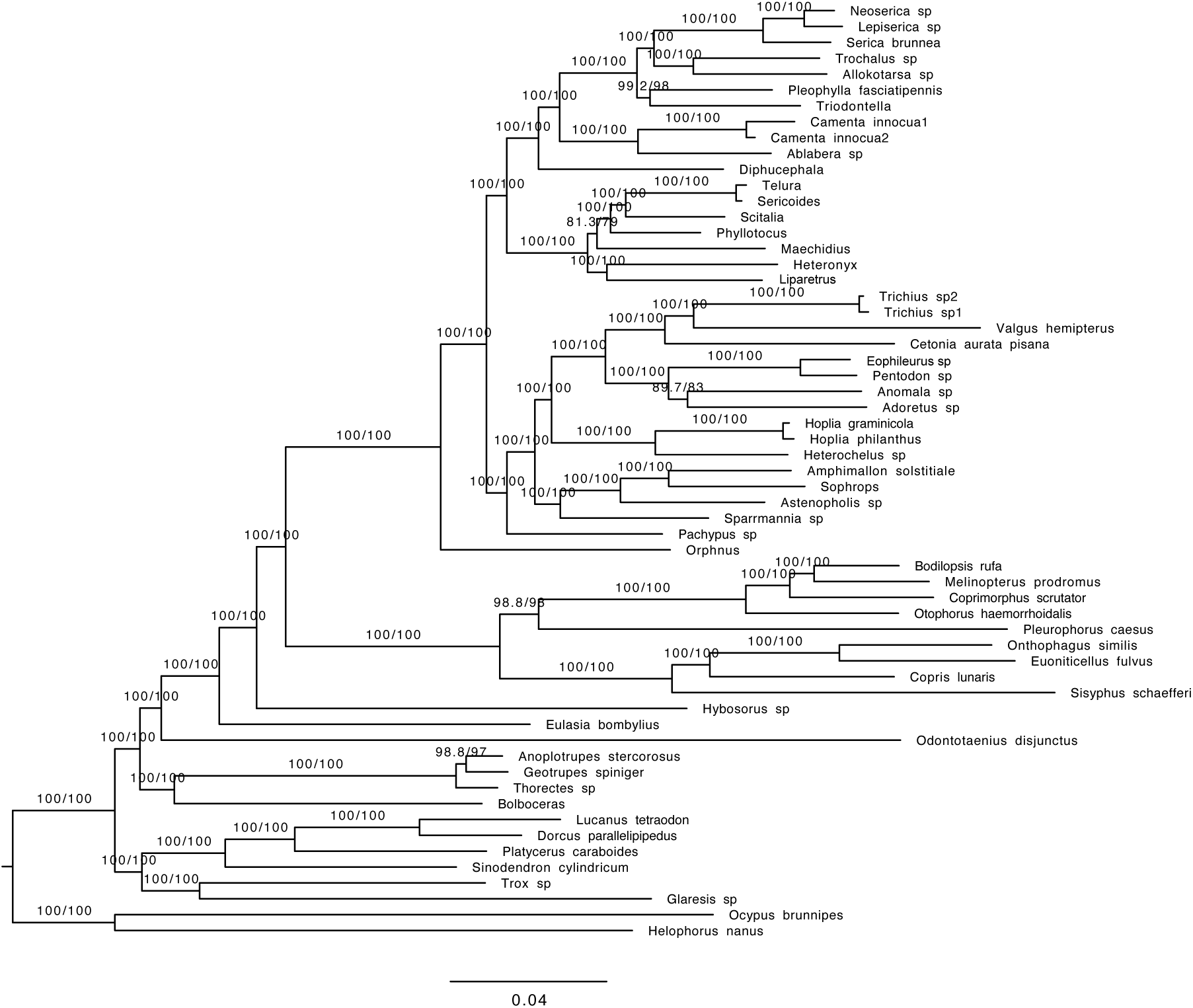
Phylogenetic tree obtained with the concatenated data and nucleotide data containing only the first and second base pair (nt12) (Mafft alignment; full dataset; Endopterygota orthologs).

**Figure S45.**
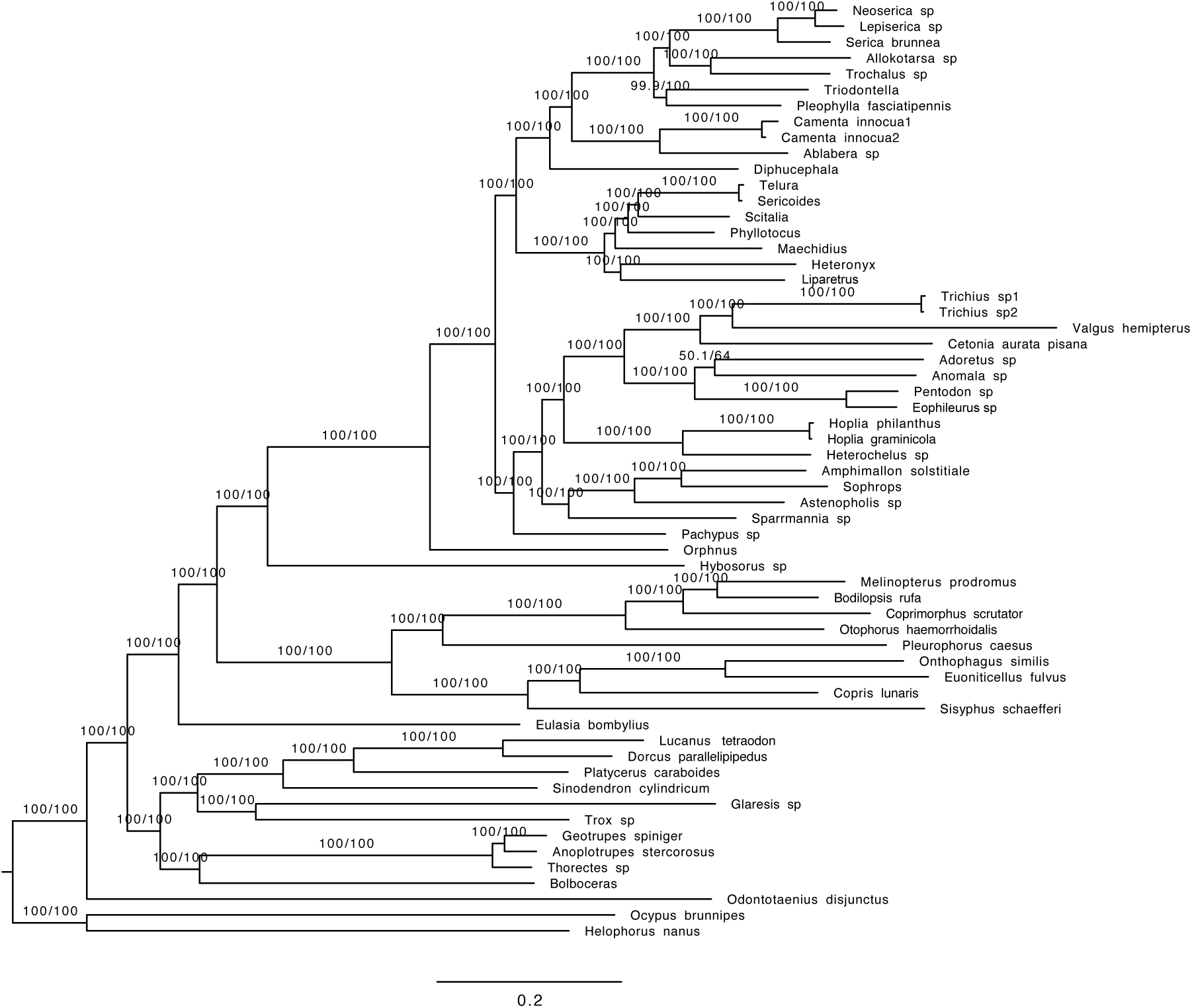
Phylogenetic tree obtained with the concatenated data and nucleotide data containing all three base pairs (nt123) (Mafft alignment; full dataset; Endopterygota orthologs).

**Figure S46.**
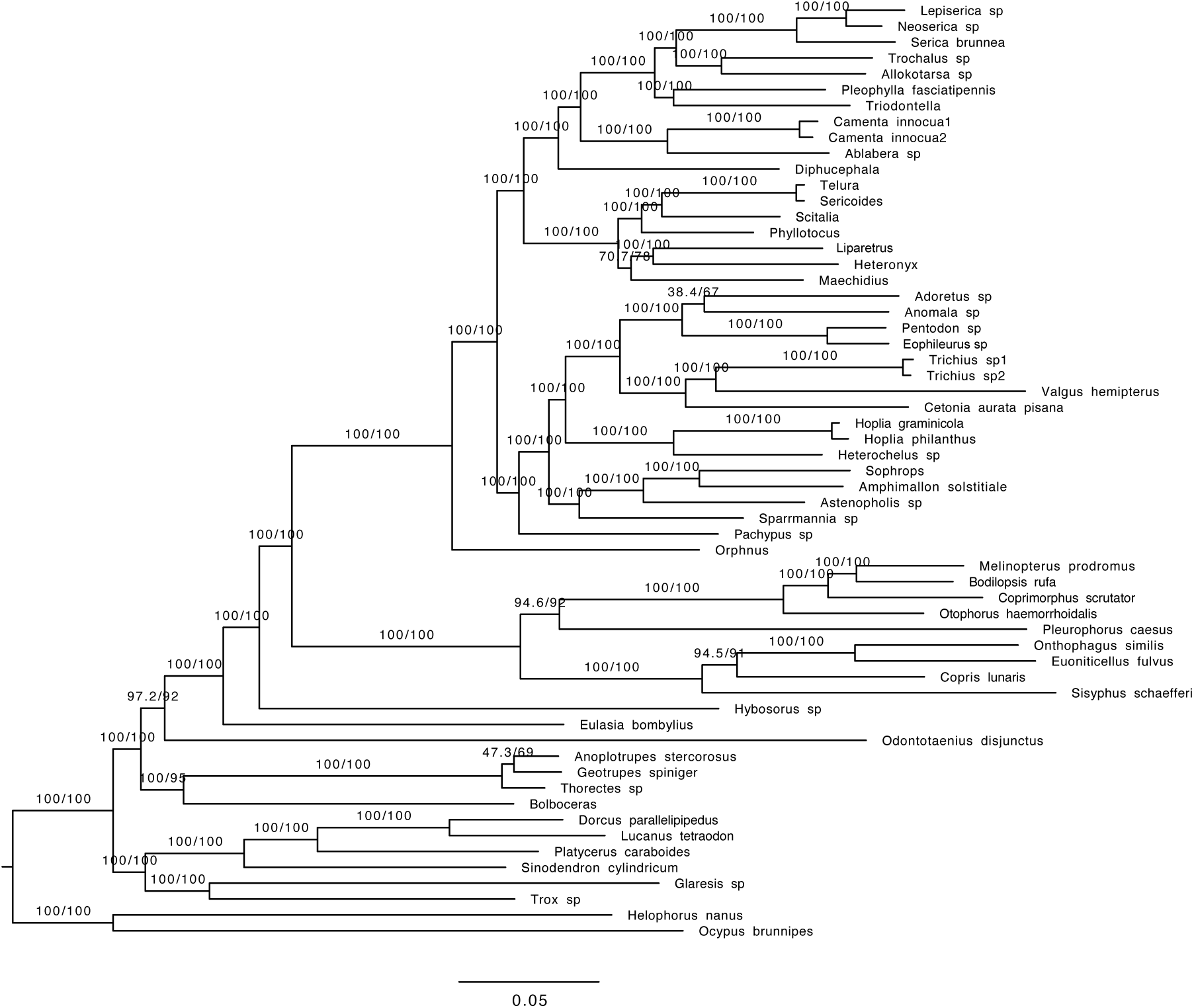
Phylogenetic tree obtained with the concatenated data and amino acid sequences (AA) (HmmAlign alignment; full dataset; Endopterygota orthologs).

**Figure S47.**
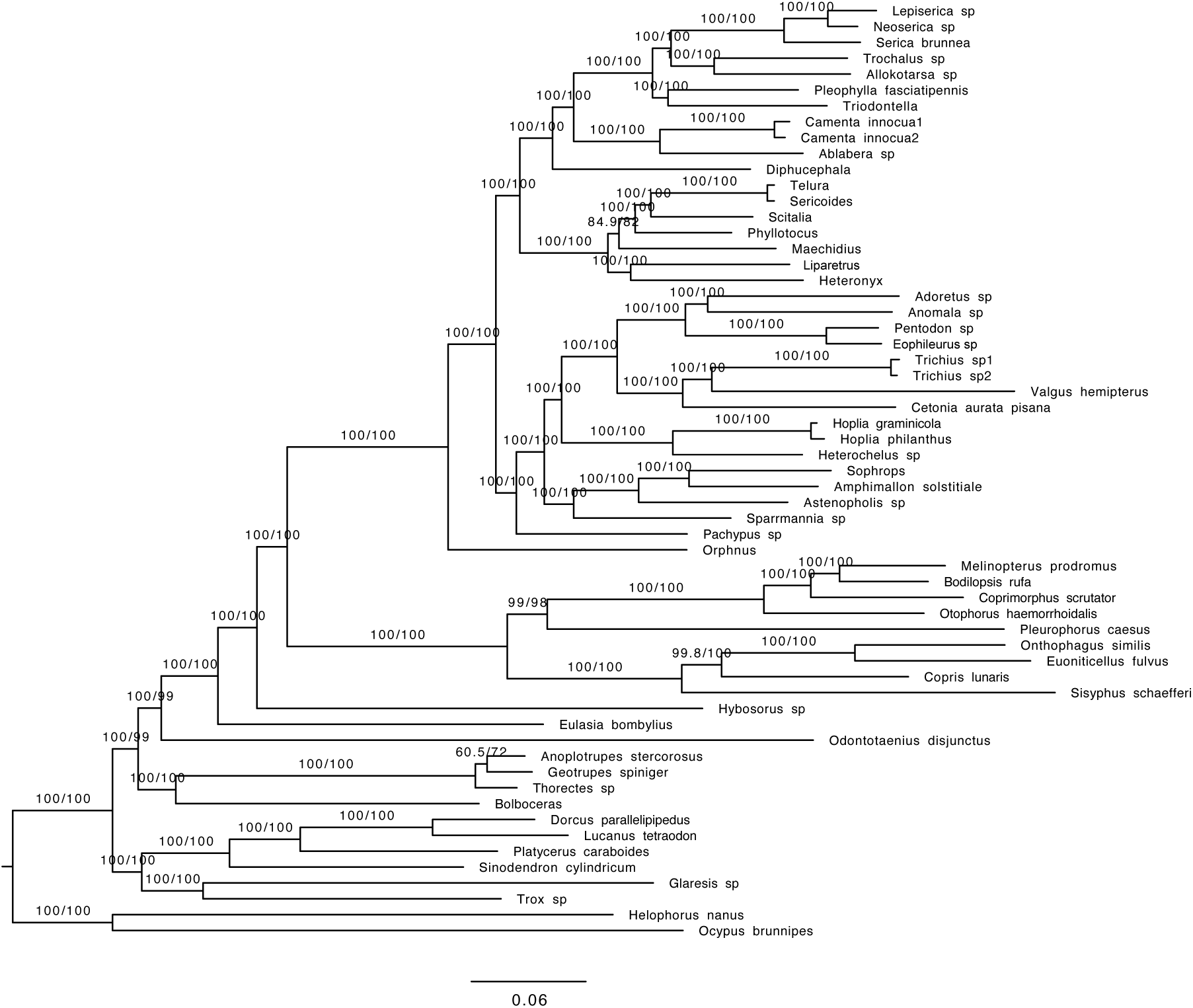
Phylogenetic tree obtained with the concatenated data and nucleotide data containing only the first and second base pair (nt12) (HmmAlign alignment; full dataset; Endopterygota orthologs).

**Figure S48.**
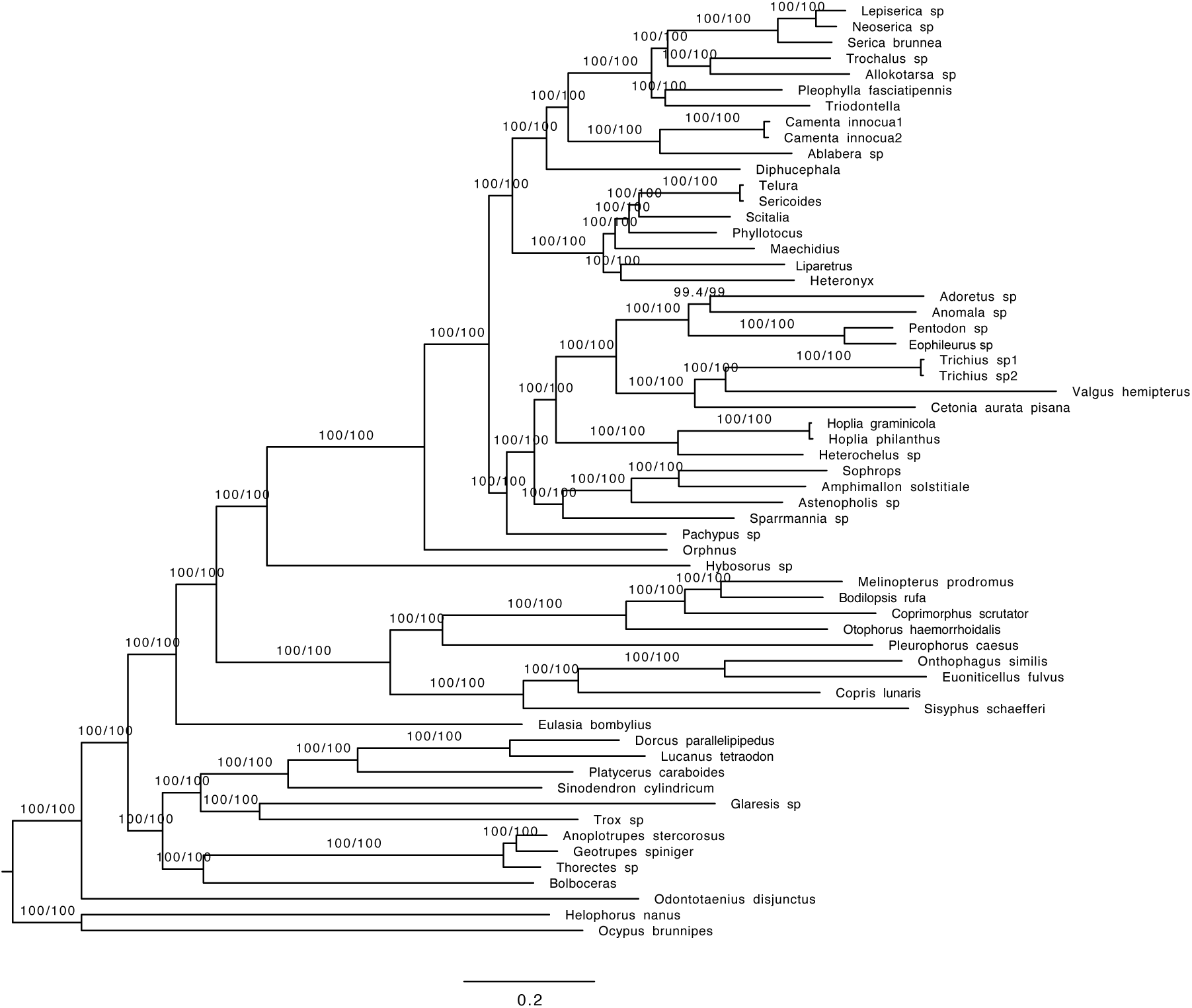
Phylogenetic tree obtained with the concatenated data and nucleotide data containing all three base pairs (nt123) (HmmAlign alignment; full dataset; Endopterygota orthologs).

**Figure S49.**
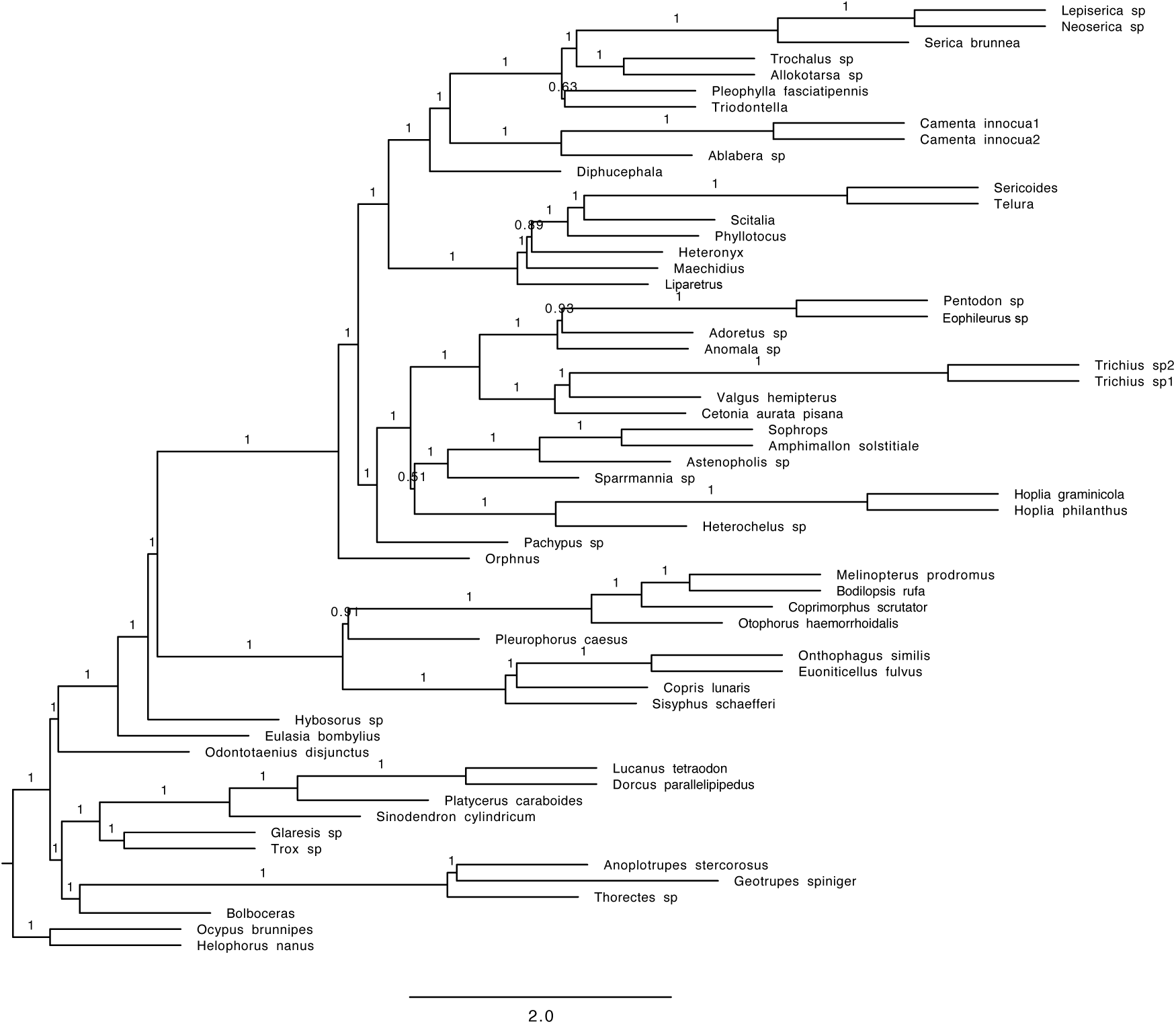
Phylogenetic tree obtained with the coalescent tree search with Astral and amino acid sequences (Mafft alignment; dataset with 70% complete data; Endopterygota orthologs).

**Figure S50.**
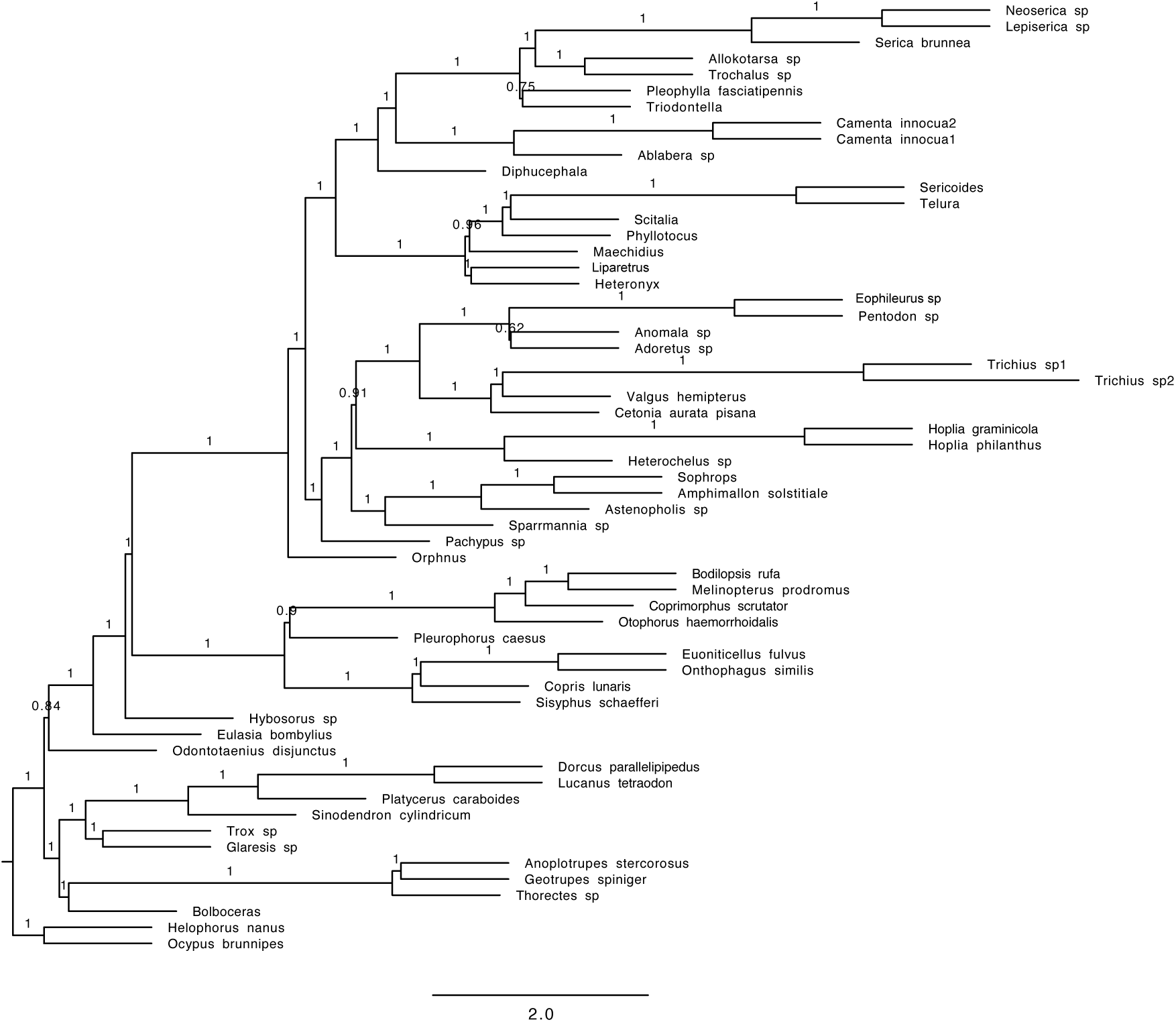
Phylogenetic tree obtained with the coalescent tree search with Astral and nucleotide data containing only the first and second base pair (nt12) (Mafft alignment; dataset with 70% complete data; Endopterygota orthologs).

**Figure S51.**
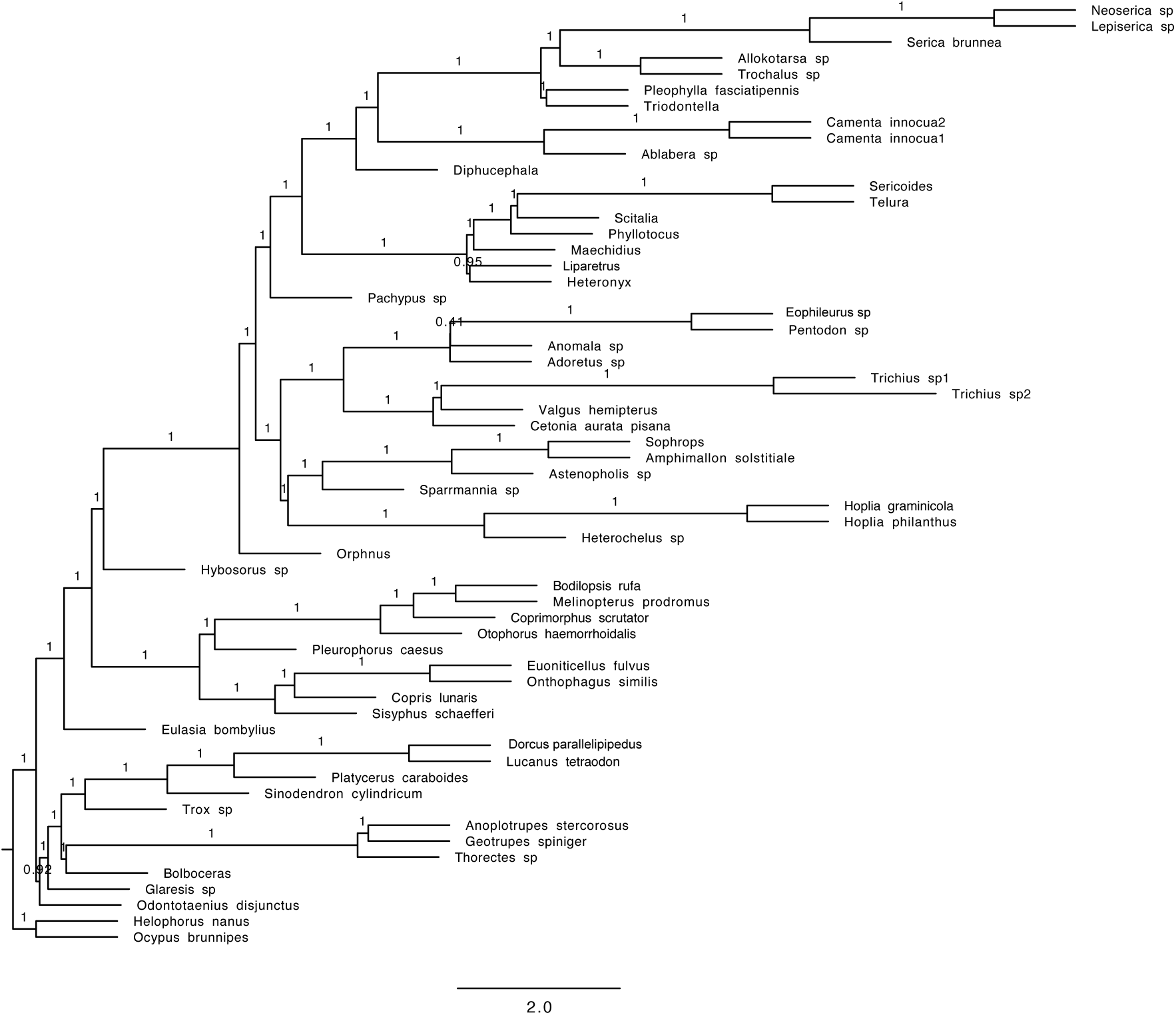
Phylogenetic tree obtained with the coalescent tree search with Astral and nucleotide data containing all three base pairs (nt123) (Mafft alignment; dataset with 70% complete data; Endopterygota orthologs).

**Figure S52.**
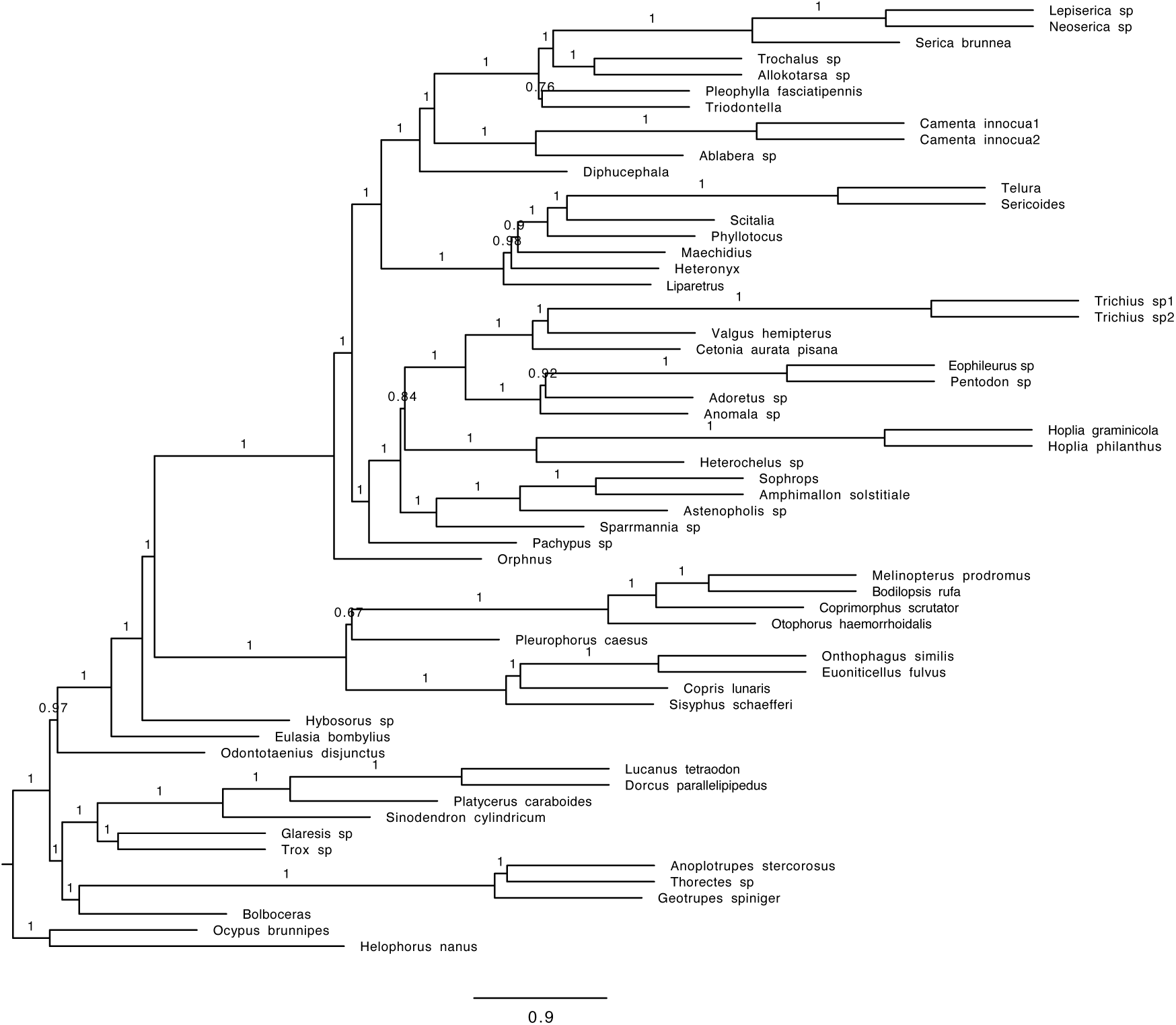
Phylogenetic tree obtained with the coalescent tree search with Astral and amino acid sequences (HmmAlign alignment; dataset with 70% complete data; Endopterygota orthologs).

**Figure S53.**
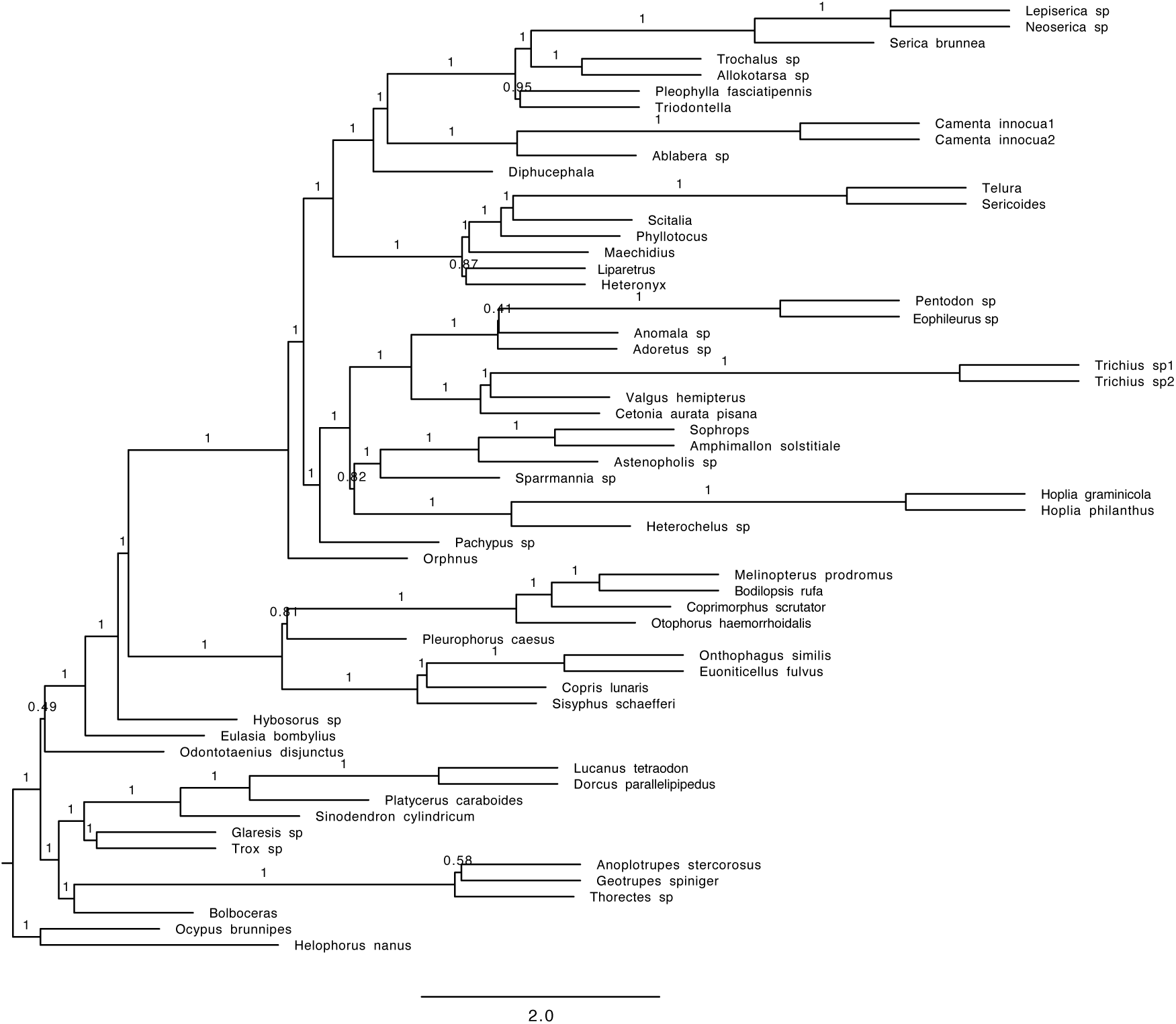
Phylogenetic tree obtained with the coalescent tree search with Astral and nucleotide data containing only the first and second base pair (nt12) (HmmAlign alignment; dataset with 70% complete data; Endopterygota orthologs).

**Figure S54.**
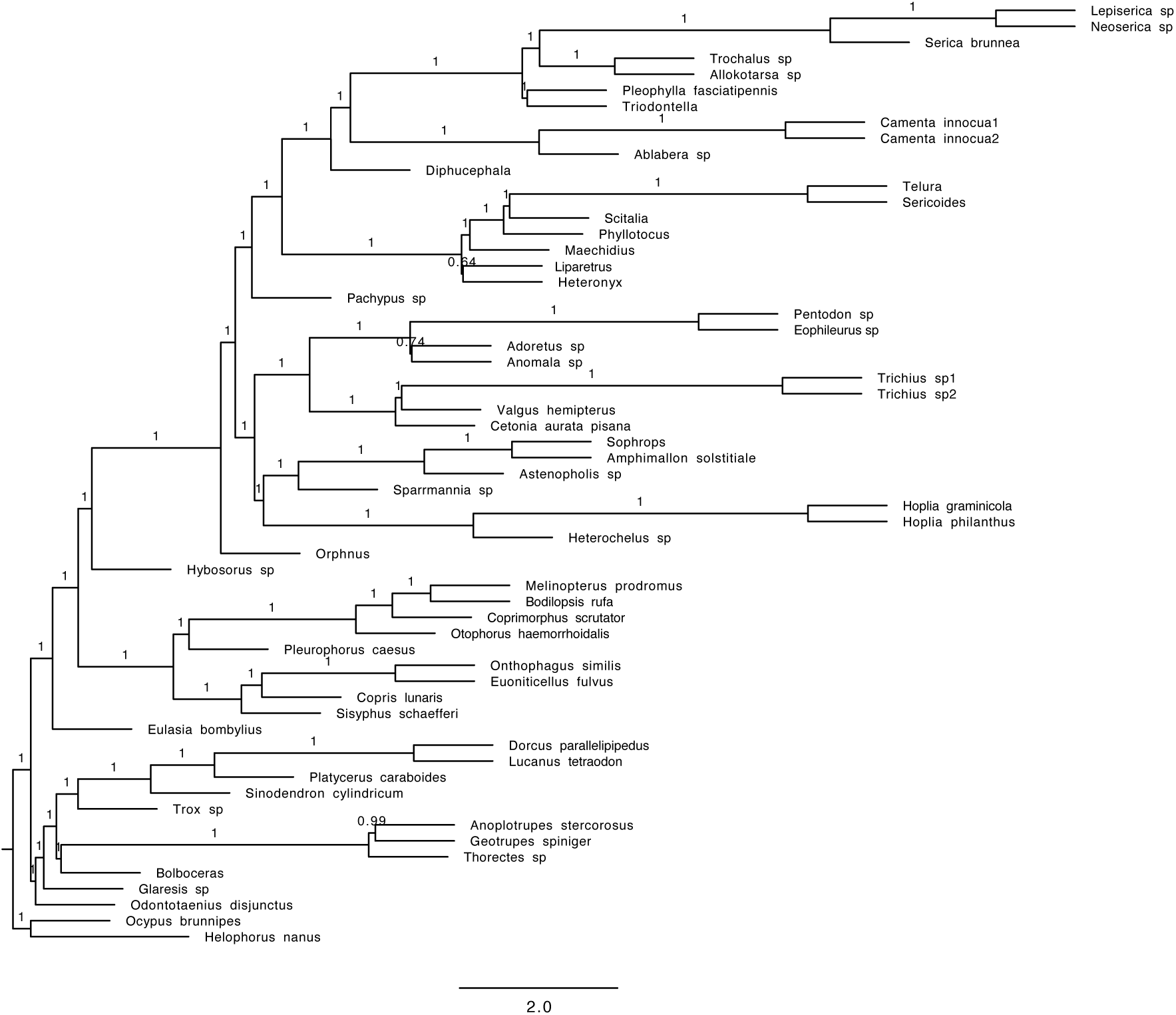
Phylogenetic tree obtained with the coalescent tree search with Astral and nucleotide data containing all three base pairs (nt123) (HmmAlign alignment; dataset with 70% complete data; Endopterygota orthologs).

**Figure S55.**
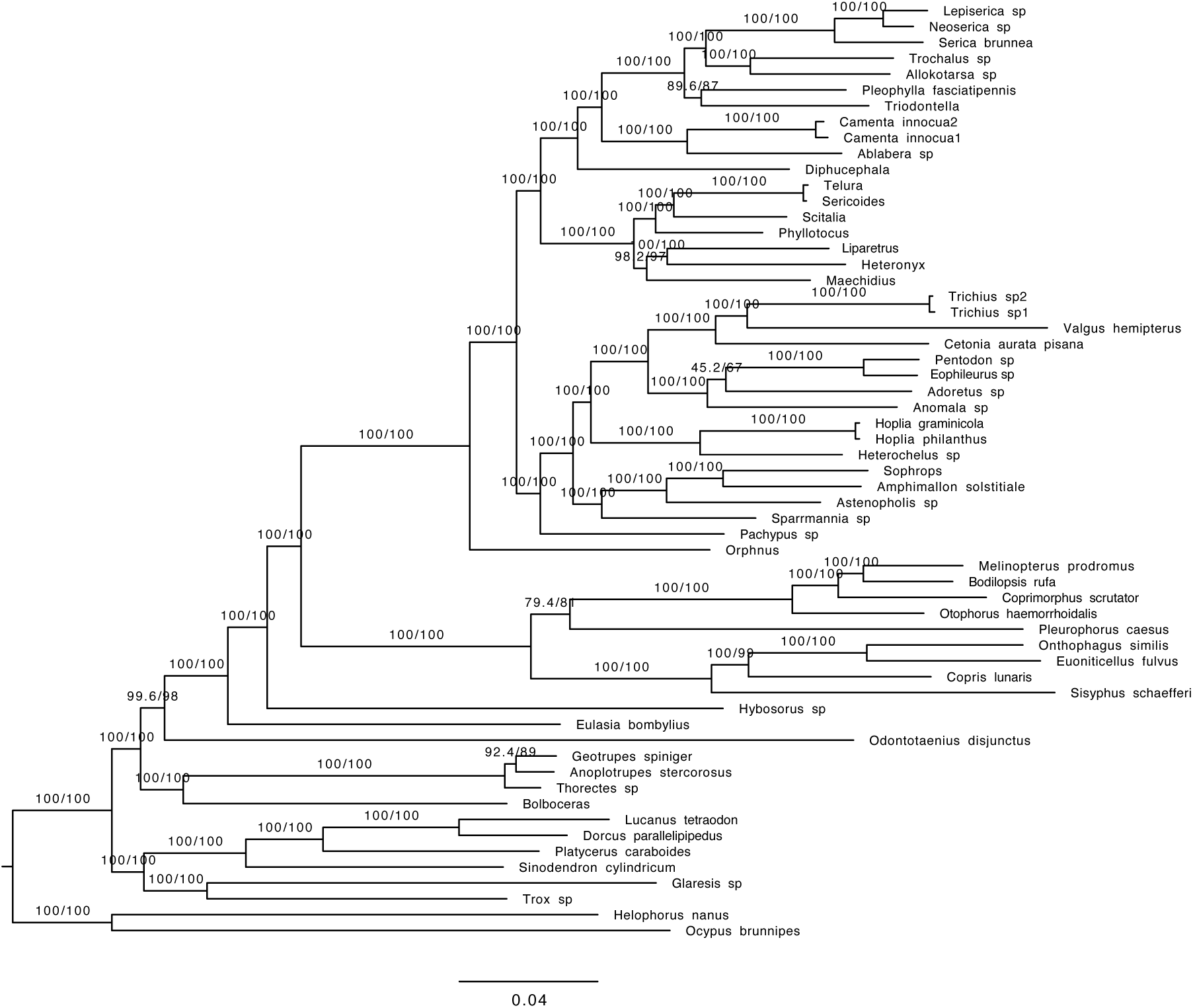
Phylogenetic tree obtained with the concatenated data and amino acid sequences (AA) (Mafft alignment; dataset with 70% complete data; Endopterygota orthologs).

**Figure S56.**
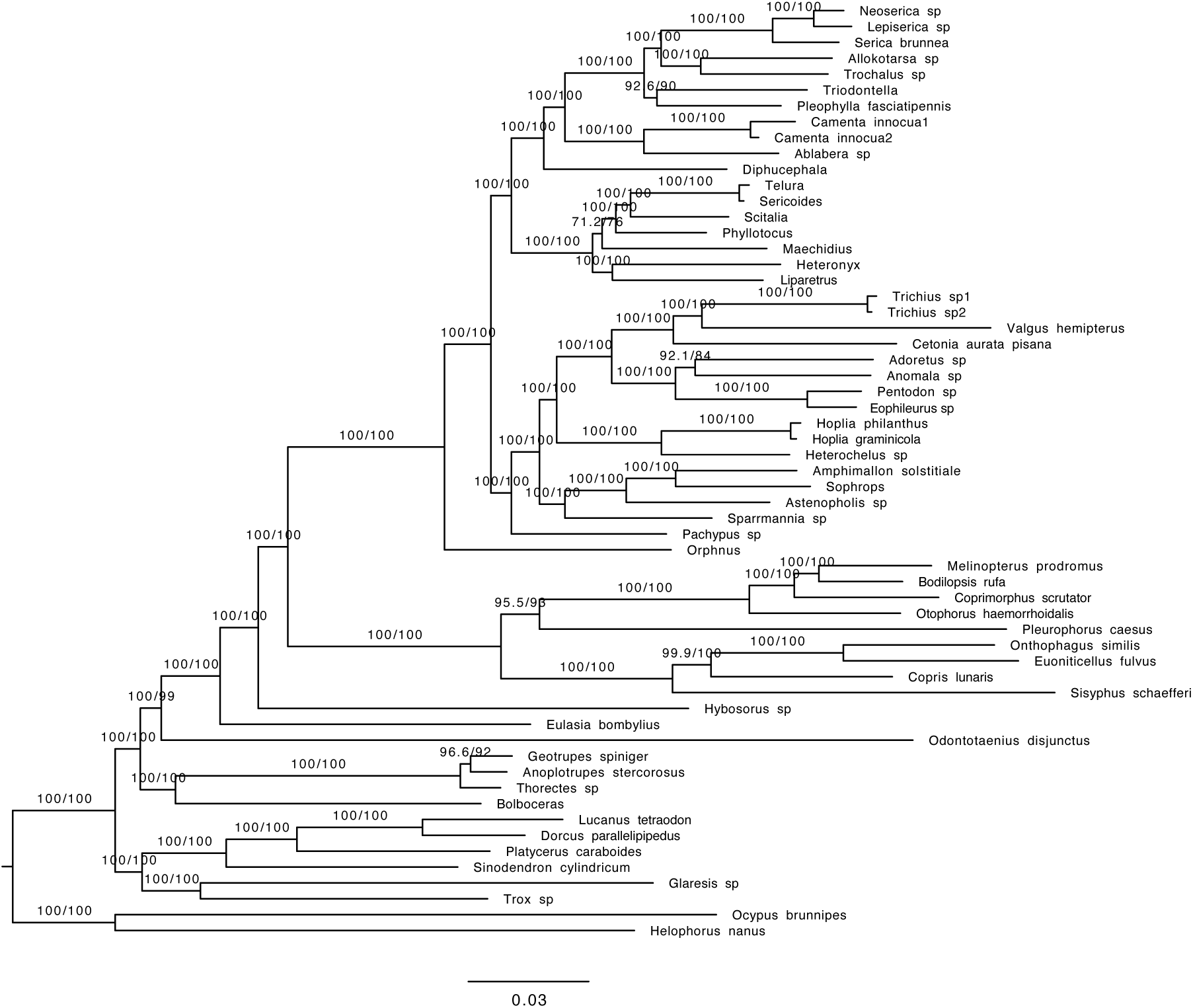
Phylogenetic tree obtained with the concatenated data and nucleotide data containing only the first and second base pair (nt12) (Mafft alignment; dataset with 70% complete data; Endopterygota orthologs).

**Figure S57.**
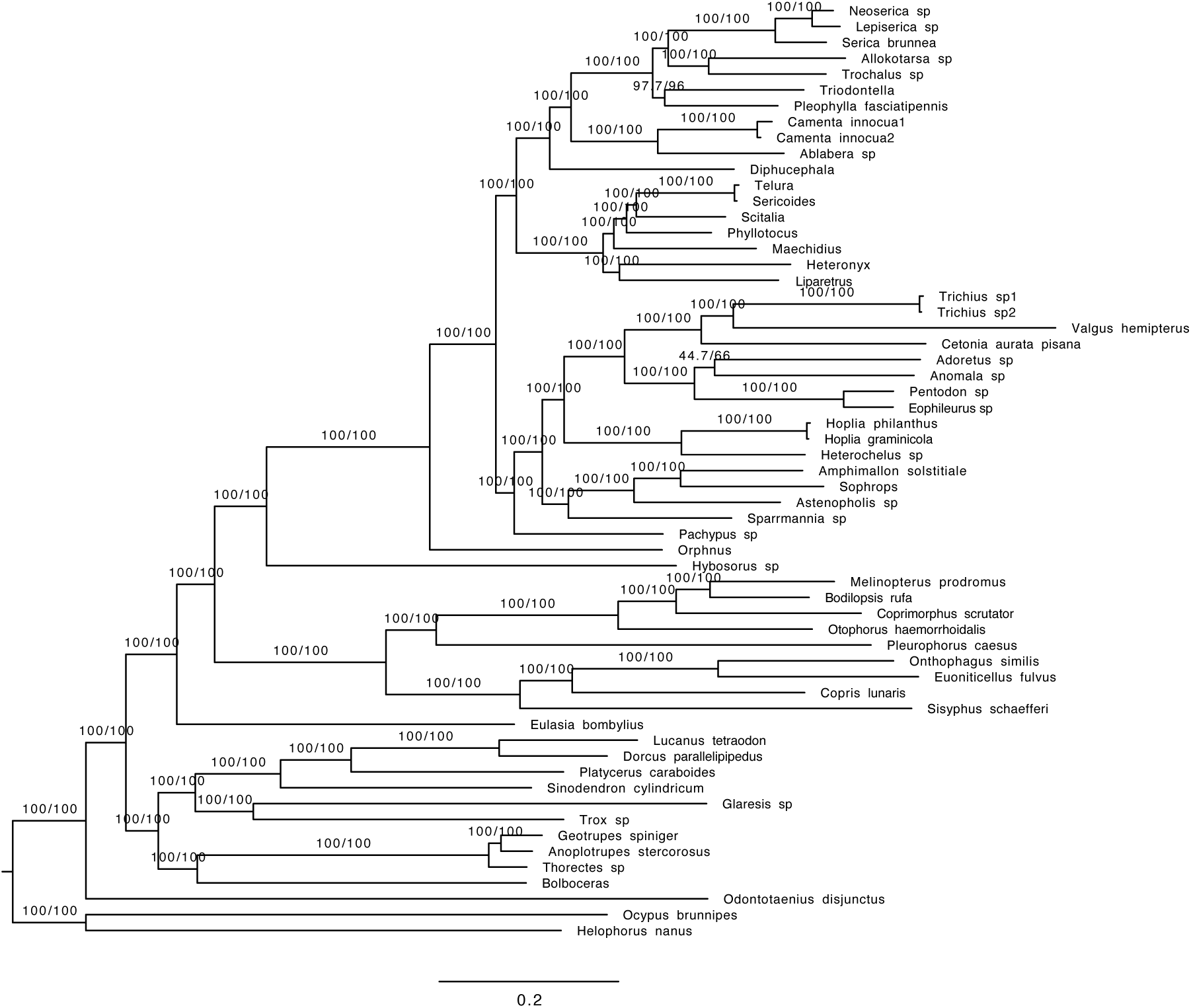
Phylogenetic tree obtained with the concatenated data and nucleotide data containing all three base pairs (nt123) (Mafft alignment; dataset with 70% complete data; Endopterygota orthologs).

**Figure S58.**
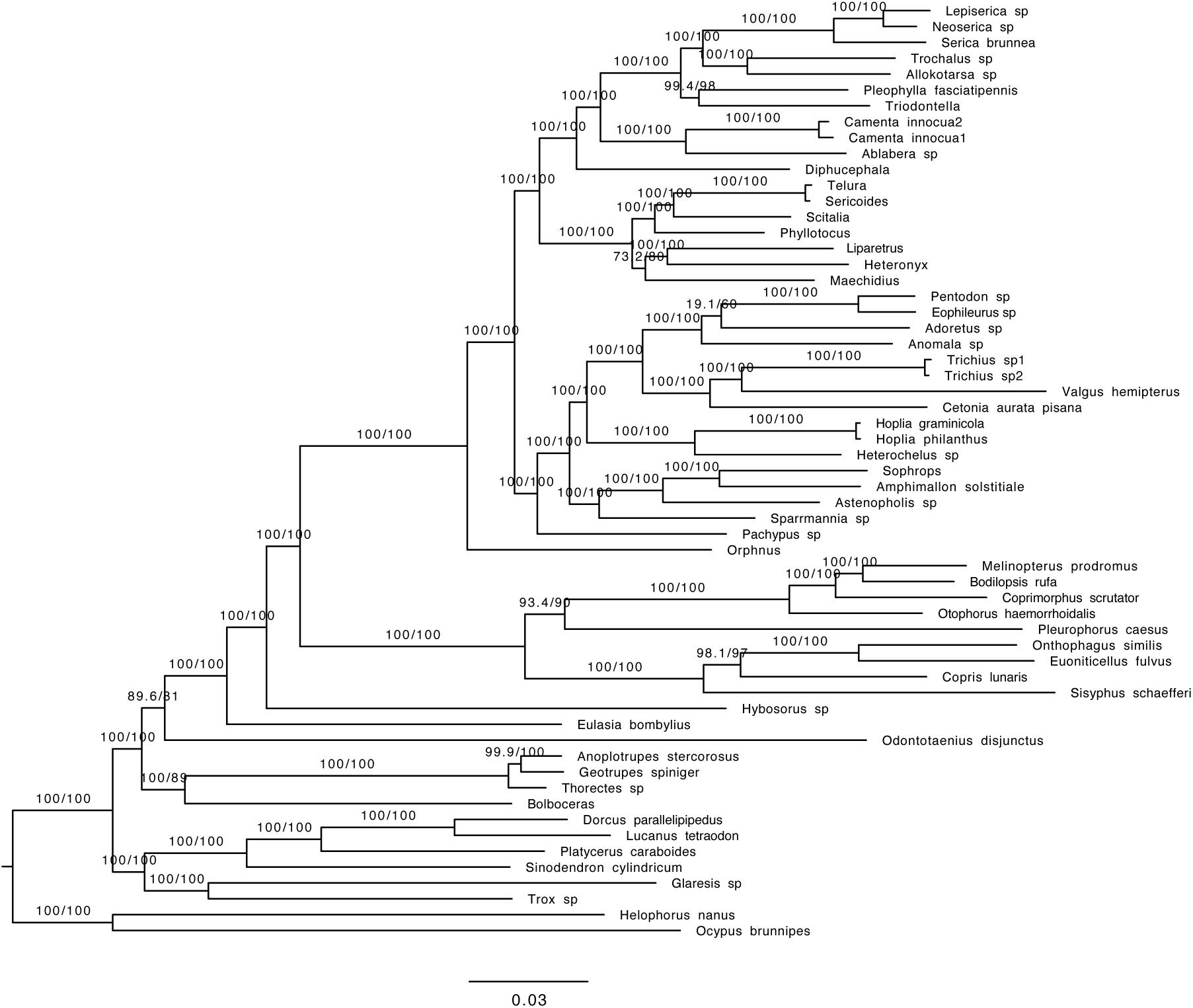
Phylogenetic tree obtained with the concatenated data and amino acid sequences (AA) HmmAlign alignment; dataset with 70% complete data; Endopterygota orthologs).

**Figure S59.**
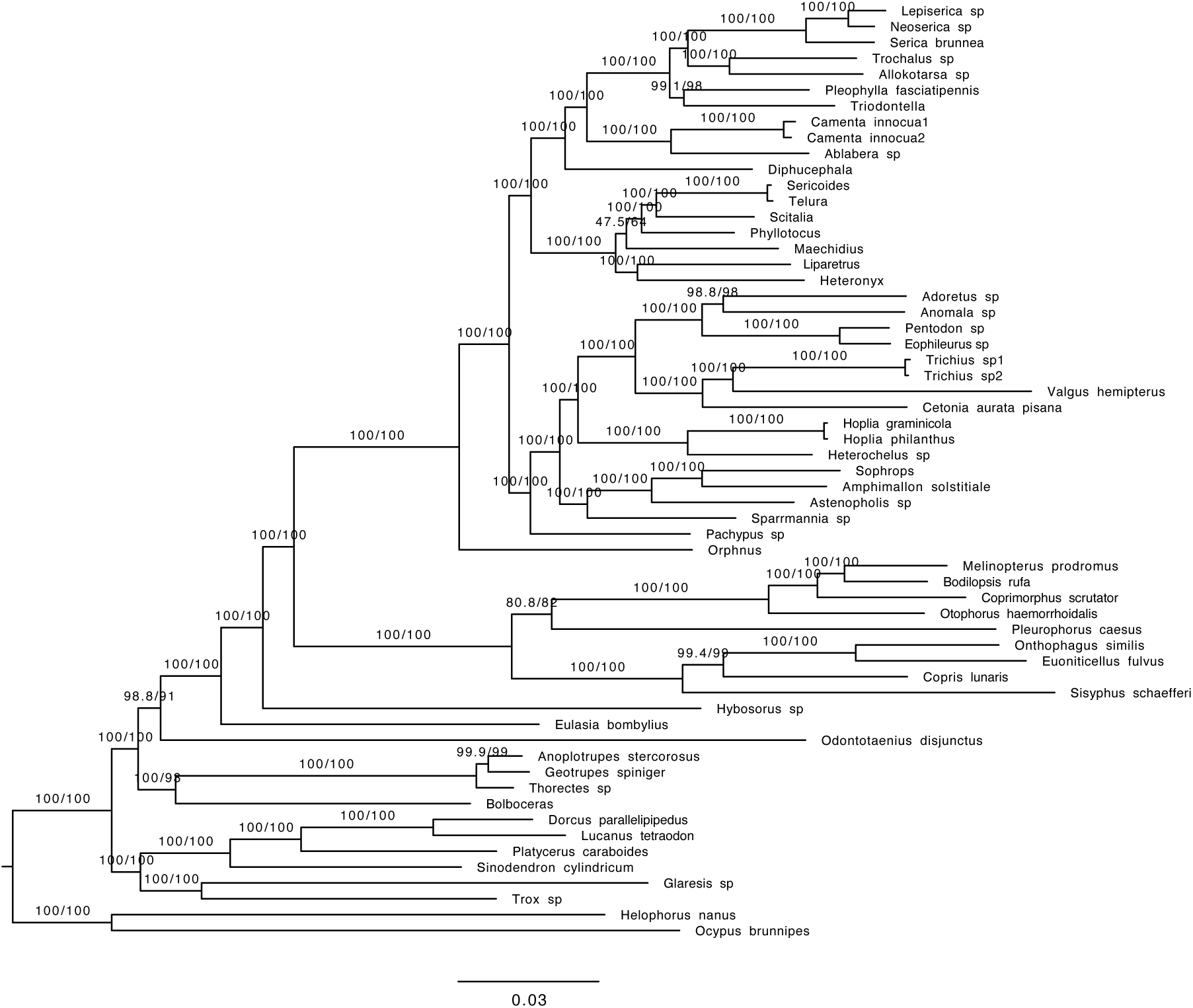
Phylogenetic tree obtained with the concatenated data and nucleotide data containing only the first and second base pair (nt12) (HmmAlign alignment; dataset with 70% complete data; Endopterygota orthologs).

**Figure S60.**
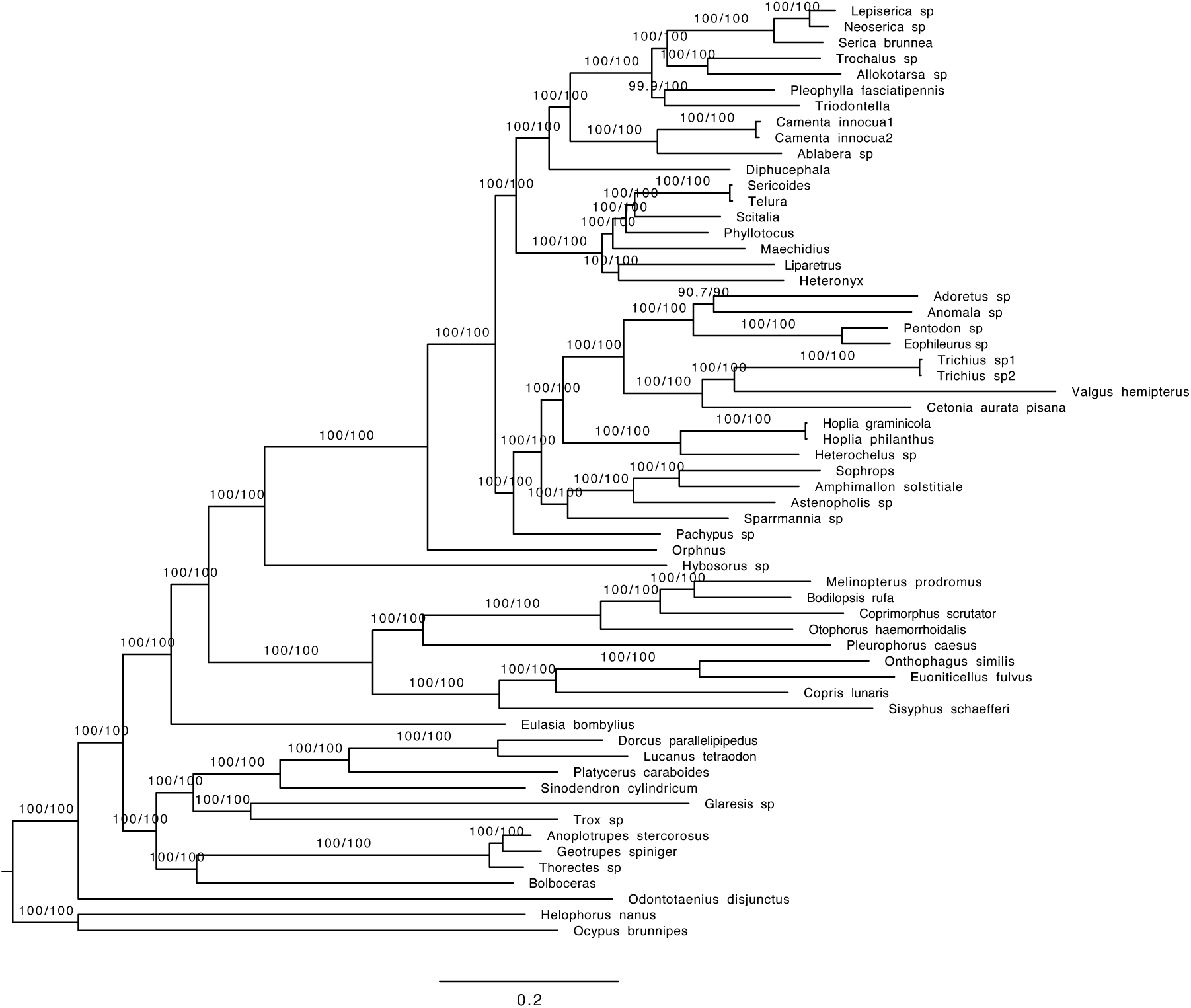
Phylogenetic tree obtained with the concatenated data and nucleotide data containing all three base pairs (nt123) (HmmAlign alignment; dataset with 70% complete data; Endopterygota orthologs).

**Suppl. Figure 61.**
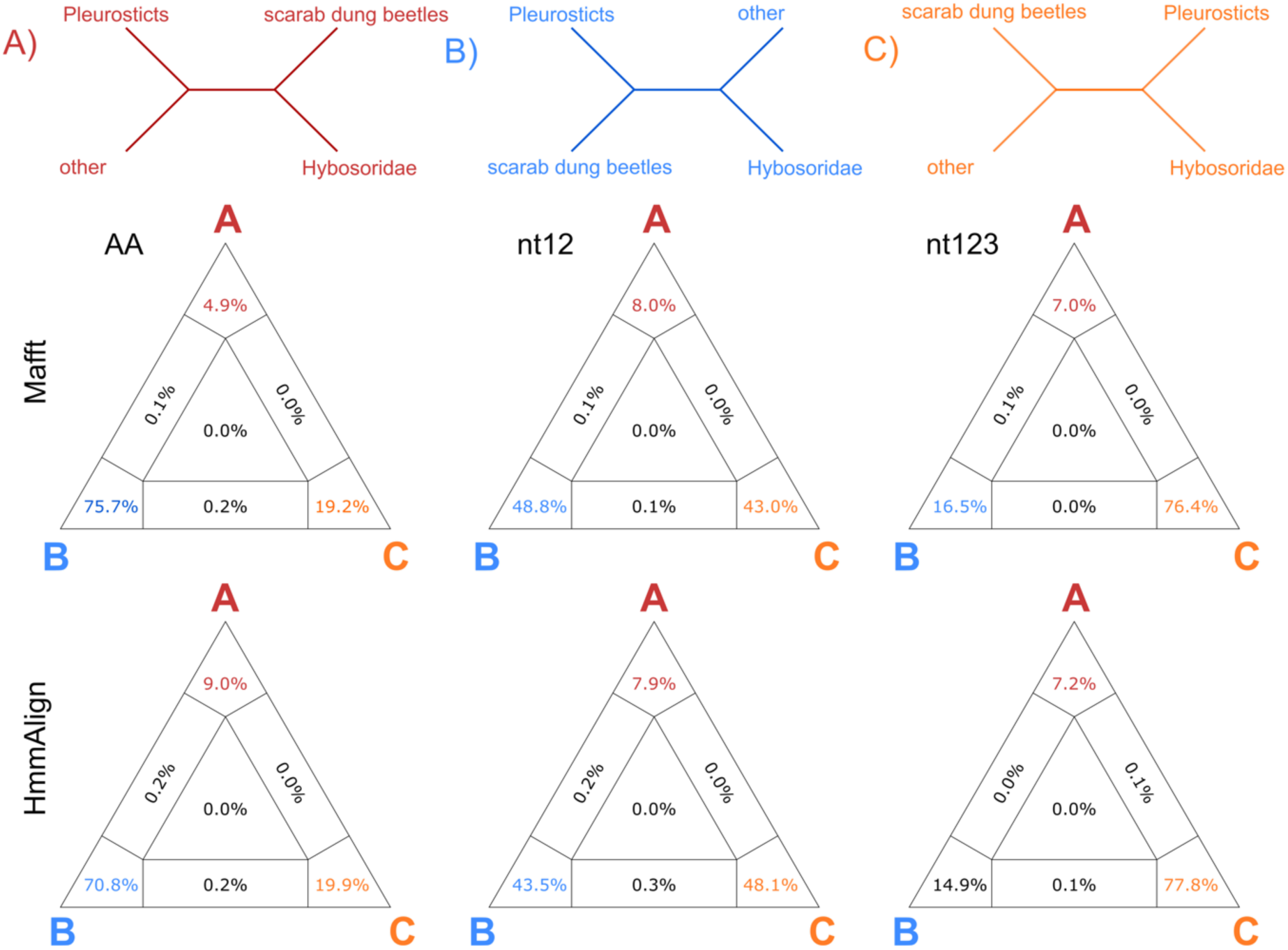
Quartet comparison of Endopterygota - single copy orthologs based on alignments with Hmmalign and Mafft, both based on the AA, nt12, and nt123 data set.

**Suppl. Figure 62.**
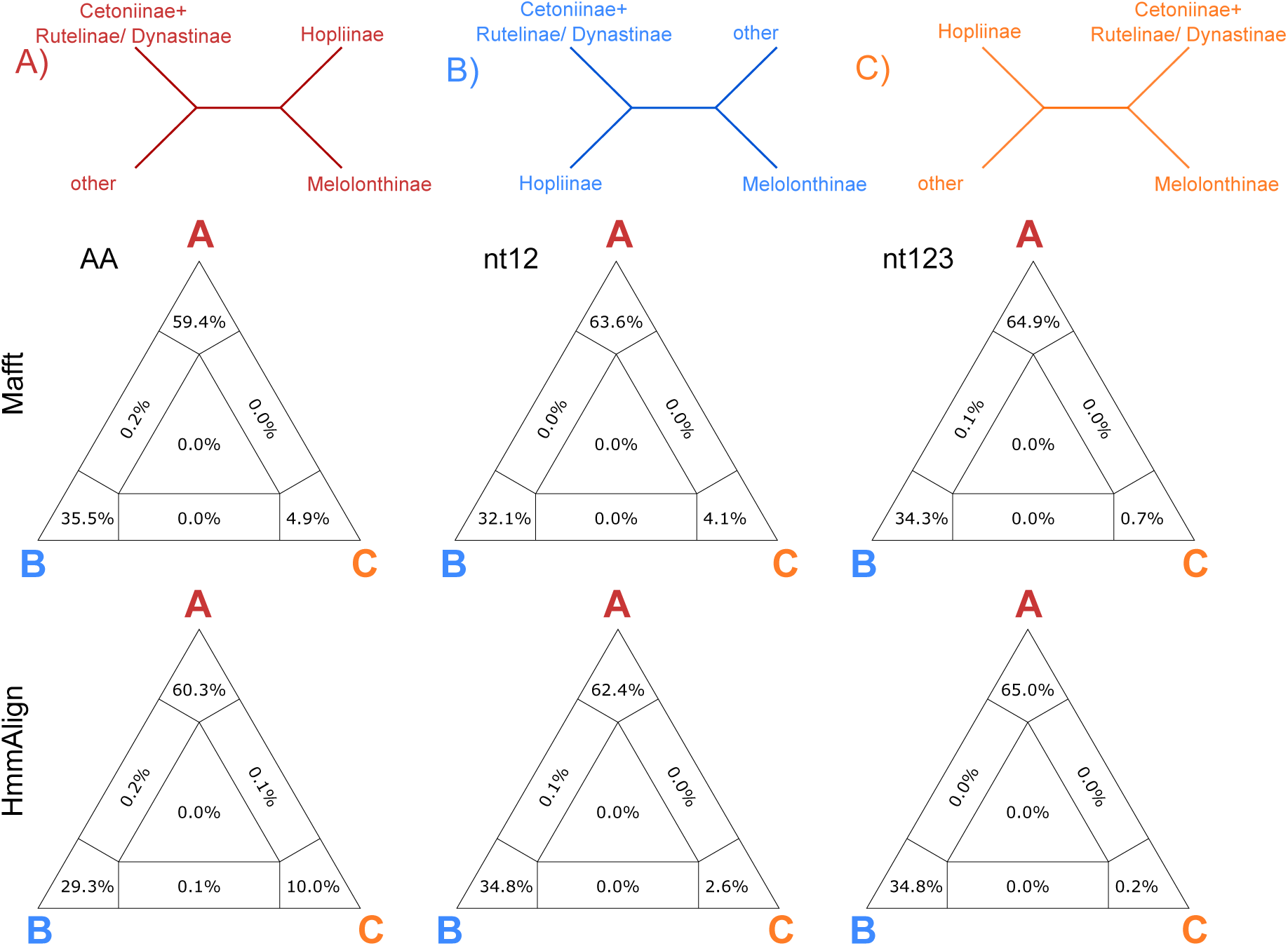
Quartet comparison of Endopterygota - single copy orthologs based on alignments with Hmmalign and Mafft, both based on the AA, nt12, and nt123 data set, regarding the monophyly of Melolonthinae + Hopliinae.

**Table S1.**
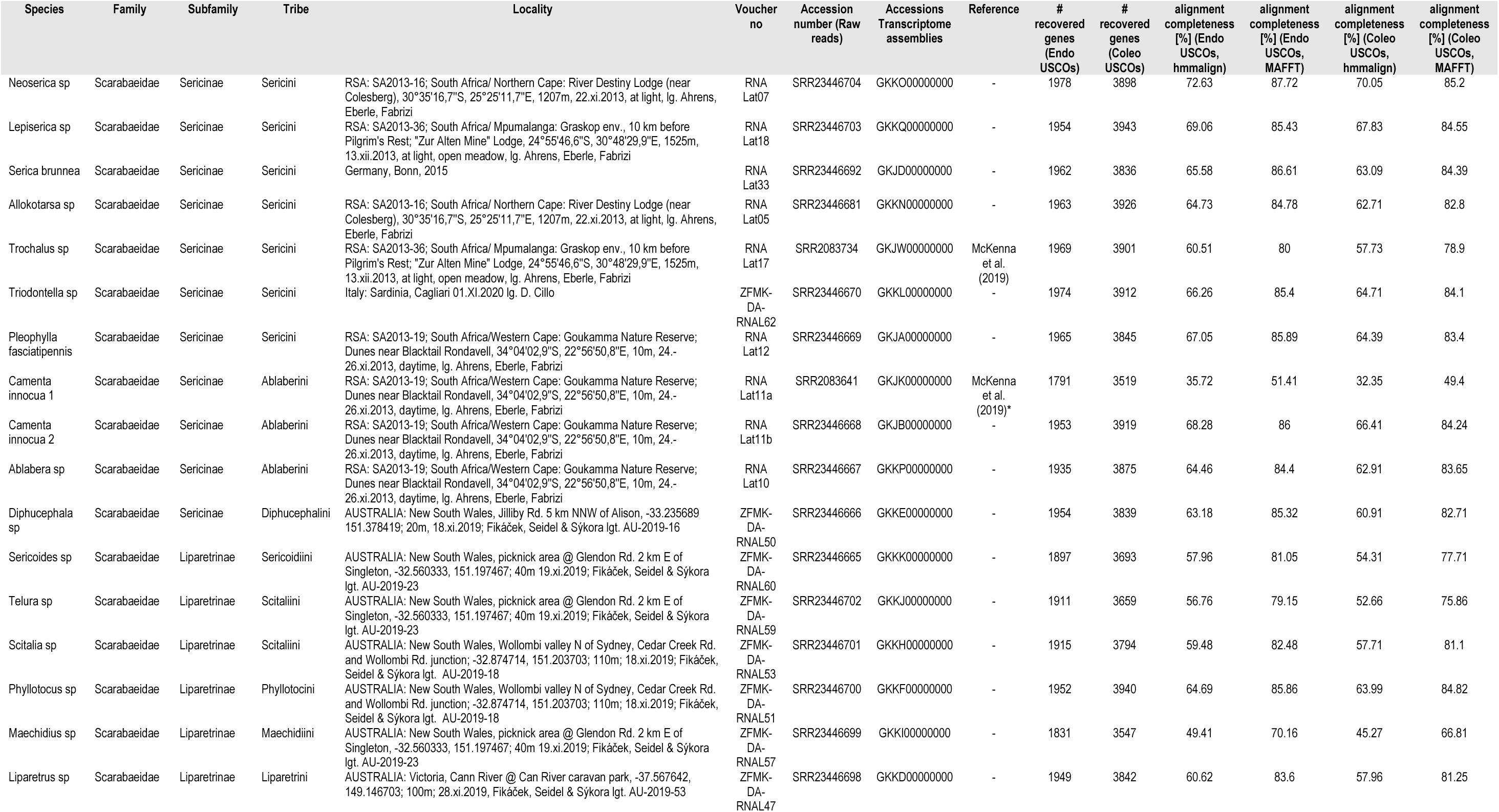
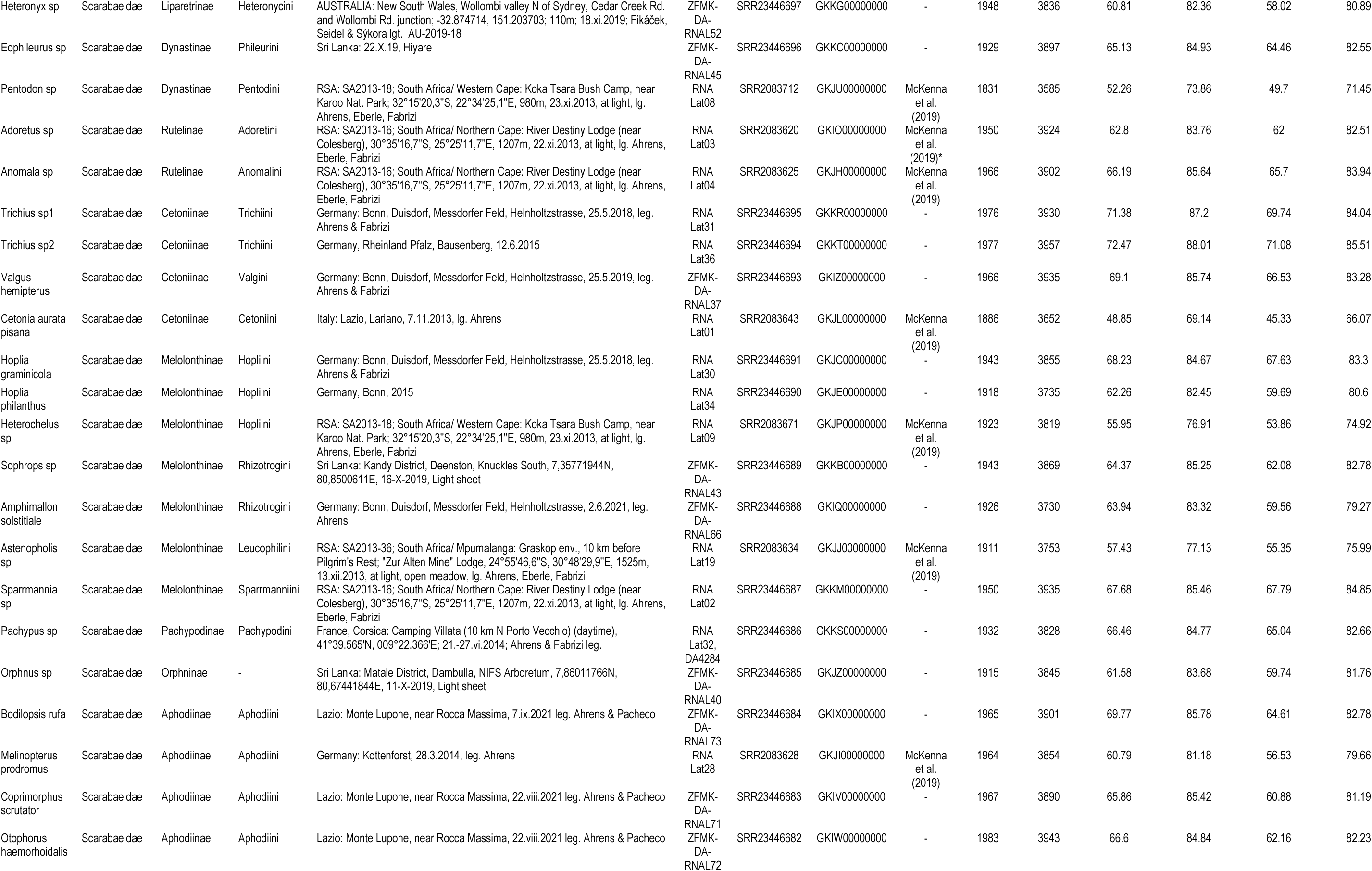
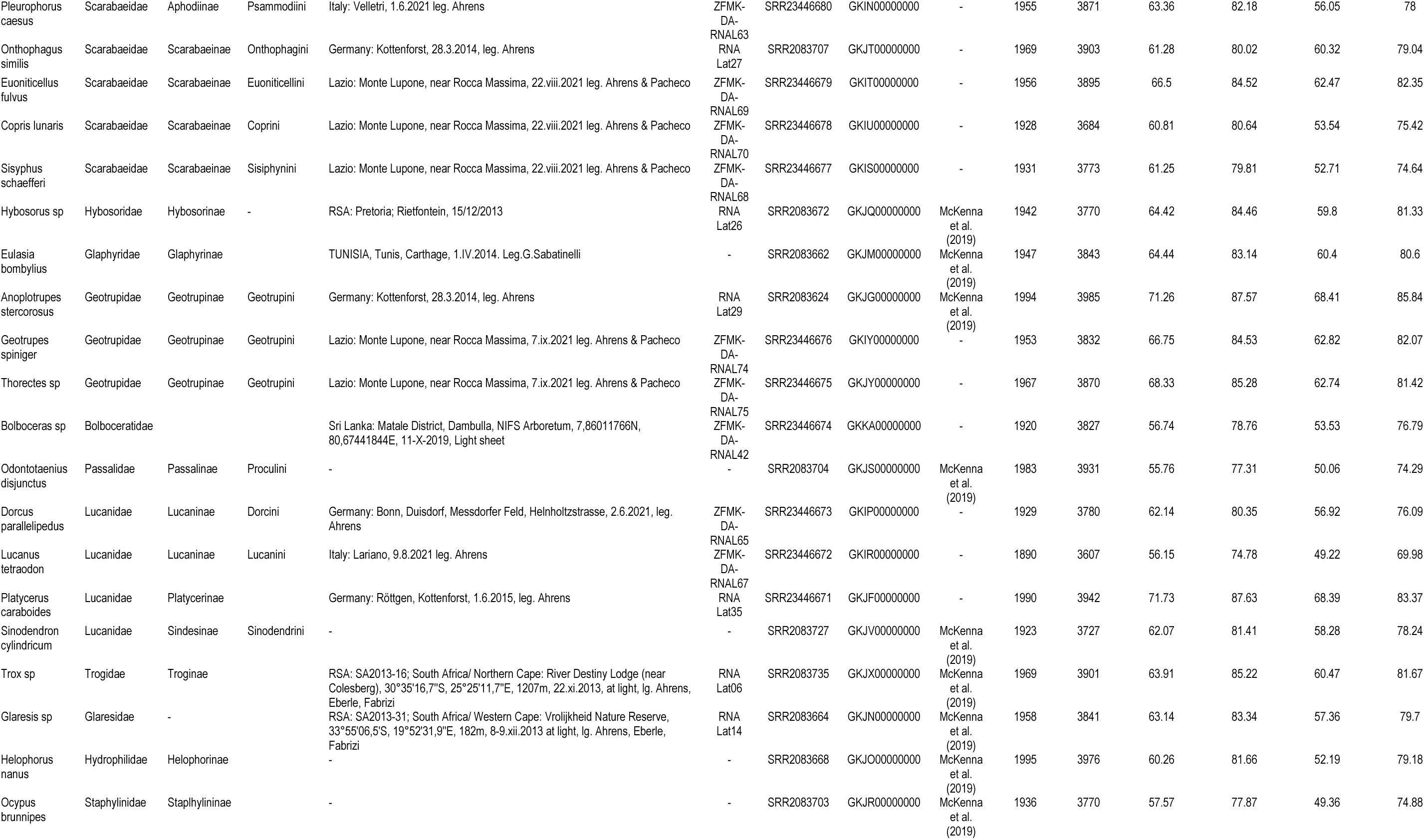
Overview on samples included in this study, referring to taxon name, current systematic placement, collection site, voucher number, accession numbers of sequence data repositories. The table also includes information on the number of recovered genes (Endopterygota USCOs vs. Coleoptera-specific genes), alignment completeness [in %] (Endopterygota USCOs, hmmalign vs. MAFFT vs. Coleoptera-specific genes, hmmalign vs. MAFFT).

**Table S2.** Likelihood tests regarding the monophyly of different groupings using the Endopterygota USCO dataset, with constraint trees using a variety of resampling tests in IQ-TREE using the RELL approximation including bootstrap proportion (BP), Kishino-Hasegawa test (KH), Shimodaira-Hasegawa test (SH), expected likelihood weights (ELW), and the approximately unbiased (AU) test.

| Tree | logL | deltaL | bp-RELL | p-KH | p-SH | c-ELW | p-AU |
| --- | --- | --- | --- | --- | --- | --- | --- |
| **hmmalign NT12** |  |  |  |  |  |  |  |
| **1: Dung scarabs + Pleurosticti** | -13896174 | 0 | 1 | 1 | 1 | 1 | 1 |
| 2: Hybosoridae + Pleurosticti | -13896819 | 645.24 | 0 | 0 | 0 | 3.11E-134 | 0.00029 |
| 3: Hybosoridae + Dung scarabs | -13896694 | 519.73 | 0 | 0 | 0 | 3.74E-61 | 2.72E-05 |
| **hmmalign NT123** |  |  |  |  |  |  |  |
| 1: Dung scarabs + Pleurosticti | -40490187 | 1800.4 | 0 | 0 | 0 | 0 | 4.53E-06 |
| **2: Hybosoridae + Pleurosticti** | -40488386 | 0 | 1 | 1 | 1 | 1 | 1 |
| 3: Hybosoridae + Dung scarabs | -40490262 | 1875.5 | 0 | 0 | 0 | 0 | 1.55E-59 |
| **hmmalign AA** |  |  |  |  |  |  |  |
| **1: Dung scarabs + Pleurosticti** | -11819613 | 0 | 1 | 1 | 1 | 1 | 1 |
| 2: Hybosoridae + Pleurosticti | -11820703 | 1090.2 | 0 | 0 | 0 | 9.97e-321 | 2.94E-06 |
| 3: Hybosoridae + Dung scarabs | -11820134 | 520.84 | 0 | 0 | 0 | 5.54E-51 | 1.97E-72 |
| **MAFFT NT12** |  |  |  |  |  |  |  |
| **1: Dung scarabs + Pleurosticti** | -12558587 | 0 | 1 | 1 | 1 | 1 | 1 |
| 2: Hybosoridae + Pleurosticti | -12559281 | 693.31 | 0 | 0 | 0 | 2.08E-162 | 1.29E-06 |
| 3: Hybosoridae + Dung scarabs | -12559109 | 521.89 | 0 | 0 | 0 | 1.21E-84 | 4.85E-06 |
| **MAFFT NT123** |  |  |  |  |  |  |  |
| 1: Dung scarabs + Pleurosticti | -36782425 | 1721.7 | 0 | 0 | 0 | 0 | 1.31E-52 |
| **2: Hybosoridae + Pleurosticti** | -36780703 | 0 | 1 | 1 | 1 | 1 | 1 |
| 3: Hybosoridae + Dung scarabs | -36782464 | 1760.9 | 0 | 0 | 0 | 0 | 4.16E-06 |
| **MAFFT AA** |  |  |  |  |  |  |  |
| **1: Dung scarabs + Pleurosticti** | -10023091 | 0 | 1 | 1 | 1 | 1 | 1 |
| 2: Hybosoridae + Pleurosticti | -10024195 | 1104.1 | 0 | 0 | 0 | 0 | 3.72E-84 |
| 3: Hybosoridae + Dung scarabs | -10023664 | 572.39 | 0 | 0 | 0 | 1.79E-81 | 0.00017 |
| **hmmalign NT12** |  |  |  |  |  |  |  |
| **1: Adoretus + Anomala** | -13896166 | 0 | 0.999 | 0.999 | 1 | 0.999 | 0.999 |
| 2: Adoretus + Dynastinae | -13896412 | 246.1 | 0.001 | 0.0007 | 0.0008 | 0.00106 | 0.00122 |
| 3: Anomala + Dynastinae | -13896558 | 391.65 | 0 | 0 | 0 | 3.93E-34 | 3.22E-05 |
| **hmmalign NT123** |  |  |  |  |  |  |  |
| **1: Adoretus + Anomala** | -40488384 | 0 | 0.989 | 0.989 | 1 | 0.989 | 0.99 |
| 2: Adoretus + Dynastinae | -40488602 | 217.46 | 0.0106 | 0.0115 | 0.017 | 0.0106 | 0.0106 |
| 3: Anomala + Dynastinae | -40488773 | 388.8 | 0 | 0 | 0 | 3.37E-19 | 3.80E-07 |
| **hmmalign AA** |  |  |  |  |  |  |  |
| **1: Adoretus + Anomala** | -11819613 | 0 | 0.646 | 0.641 | 1 | 0.646 | 0.638 |
| 2: Adoretus + Dynastinae | -11819645 | 31.7 | 0.354 | 0.359 | 0.501 | 0.354 | 0.362 |
| 3: Anomala + Dynastinae | -11819998 | 384.88 | 0 | 0 | 0 | 1.07E-68 | 7.50E-47 |
| **MAFFT NT12** |  |  |  |  |  |  |  |
| **1: Adoretus + Anomala** | -12558590 | 0 | 0.836 | 0.892 | 1 | 0.836 | 0.917 |
| 2: Adoretus + Dynastinae | -12558699 | 109.33 | 0.0747 | 0.0915 | 0.15 | 0.0749 | 0.111 |
| 3: Anomala + Dynastinae | -12558691 | 101.52 | 0.0891 | 0.108 | 0.179 | 0.089 | 0.132 |
| **MAFFT NT123** |  |  |  |  |  |  |  |
| **1: Adoretus + Anomala** | -36780719 | 0 | 0.63 | 0.699 | 1 | 0.63 | 0.754 |
| 2: Adoretus + Dynastinae | -36780770 | 50.741 | 0.246 | 0.301 | 0.44 | 0.246 | 0.348 |
| 3: Anomala + Dynastinae | -36780805 | 85.785 | 0.124 | 0.189 | 0.293 | 0.124 | 0.219 |
| **MAFFT AA** |  |  |  |  |  |  |  |
| 1: Adoretus + Anomala | -10023128 | 37.129 | 0.328 | 0.338 | 0.477 | 0.329 | 0.336 |
| **2: Adoretus + Dynastinae** | -10023091 | 0 | 0.67 | 0.662 | 1 | 0.669 | 0.687 |
| 3: Anomala + Dynastinae | -10023296 | 204.98 | 0.0019 | 0.0078 | 0.0131 | 0.00188 | 0.00412 |
| **hmmalign NT12** |  |  |  |  |  |  |  |
| **1: Hopliini + Dynastinae etc.** | -13896163 | 0 | 1 | 1 | 1 | 1 | 1 |
| 2: Melolonthinae + Dynastinae etc. | -13897228 | 1064.9 | 0 | 0 | 0 | 9.88e-324 | 0.00032 |
| 3: Melolonthinae + Hopliini | -13896914 | 751.4 | 0 | 0 | 0 | 4.33E-189 | 3.21E-05 |
| **hmmalign NT123** |  |  |  |  |  |  |  |
| **1: Hopliini + Dynastinae etc.** | -40488268 | 0 | 1 | 1 | 1 | 1 | 1 |
| 2: Melolonthinae + Dynastinae etc. | -40490887 | 2618.5 | 0 | 0 | 0 | 0 | 5.34E-05 |
| 3: Melolonthinae + Hopliini | -40490321 | 2052.5 | 0 | 0 | 0 | 0 | 2.04E-48 |
| **hmmalign AA** |  |  |  |  |  |  |  |
| **1: Hopliini + Dynastinae etc.** | -11819612 | 0 | 1 | 1 | 1 | 1 | 1 |
| 2: Melolonthinae + Dynastinae etc. | -11820457 | 844.27 | 0 | 0 | 0 | 1.40E-213 | 1.57E-70 |
| 3: Melolonthinae + Hopliini | -11820241 | 628.59 | 0 | 0 | 0 | 7.87E-98 | 1.71E-37 |
| **MAFFT NT12** |  |  |  |  |  |  |  |
| **1: Hopliini + Dynastinae etc.** | -12558590 | 0 | 1 | 1 | 1 | 1 | 1 |
| 2: Melolonthinae + Dynastinae etc. | -12559724 | 1133.6 | 0 | 0 | 0 | 0 | 2.87E-07 |
| 3: Melolonthinae + Hopliini | -12559436 | 845.57 | 0 | 0 | 0 | 1.33E-231 | 2.56E-57 |
| **MAFFT NT123** |  |  |  |  |  |  |  |
| **1: Hopliini + Dynastinae etc.** | -36780491 | 0 | 1 | 1 | 1 | 1 | 1 |
| 2: Melolonthinae + Dynastinae etc. | -36783179 | 2688.9 | 0 | 0 | 0 | 0 | 3.67E-05 |
| 3: Melolonthinae + Hopliini | -36782640 | 2149.8 | 0 | 0 | 0 | 0 | 2.82E-43 |
| **MAFFT AA** |  |  |  |  |  |  |  |
| **1: Hopliini + Dynastinae etc.** | -10023092 | 0 | 1 | 1 | 1 | 1 | 1 |
| 2: Melolonthinae + Dynastinae etc. | -10024095 | 1003.6 | 0 | 0 | 0 | 2.89E-300 | 3.51E-44 |
| 3: Melolonthinae + Hopliini | -10024017 | 925.64 | 0 | 0 | 0 | 1.60E-254 | 1.47E-41 |

